# Species-wide genomics of kākāpō provides transformational tools to accelerate recovery

**DOI:** 10.1101/2022.10.22.513130

**Authors:** Joseph Guhlin, Marissa F. Le Lec, Jana Wold, Emily Koot, David Winter, Patrick Biggs, Stephanie J. Galla, Lara Urban, Yasmin Foster, Murray P. Cox, Andrew Digby, Lydia Uddstrom, Daryl Eason, Deidre Vercoe, Tāne Davis, Kākāpō Recovery Team, Jason T Howard, Erich Jarvis, Fiona E. Robertson, Bruce C. Robertson, Neil Gemmell, Tammy E. Steeves, Anna W. Santure, Peter K. Dearden

## Abstract

The kākāpō is a critically endangered, intensively managed, long-lived nocturnal parrot endemic to Aotearoa New Zealand. We generated and analyzed whole-genome sequence data for nearly all individuals living in early 2018 (169 individuals) to generate a high-quality species-wide genetic variant callset. We leverage extensive long-term metadata to quantify genome-wide diversity of the species over time and present new approaches using probabilistic programming, combined with a phenotype dataset spanning five decades, to disentangle phenotypic variance into environmental and genetic effects while quantifying uncertainty in small populations. We find associations for growth, disease susceptibility, clutch size, and egg fertility within genic regions previously shown to influence these traits in other species. Finally, we generate breeding values to predict phenotype and illustrate that active management over the past 45 years has maintained both genome-wide diversity and diversity in breeding values, and hence, evolutionary potential. We provide new pathways for informing future conservation management decisions for kākāpō, including prioritizing individuals for translocation and monitoring individuals with poor growth or high disease risk. Overall, by explicitly addressing the challenge of small sample size, we provide a template for the inclusion of genomic data that will be transformational for species recovery efforts around the globe.

## Introduction

Evidence-based conservation management of critically endangered species is crucial as human activities push more and more species to the brink of extinction. The increasing availability of population genomic and long-term phenotypic datasets for species of conservation concern offers exciting opportunities to inform conservation management ^1–4^ and enhance conservation outcomes. Critically endangered species, by definition, have small population sizes and are subject to strong genetic drift ^5–8^. For intensively managed species, the ultimate goal is to minimize the loss of genetic diversity to maintain long-term evolutionary potential. Despite the recent explosion of whole-genome sequence data for many threatened species ^9^, few studies have monitored genome-wide diversity at a population scale over time, nor have they linked remaining genetic variation to traits of conservation interest. Here we use whole-genome sequencing of nearly the entire extant population of kākāpō (*Strigops habroptilus*), combined with an extensive long-term, multi-generational, phenotypic and life-history dataset to develop methods and approaches to transform the conservation management of this taonga (treasured) species. We provide the first empirical population-level assessment of genome-wide diversity over multiple generations in an intensively managed, long-lived bird, using an up-to-date, high-quality reference genome and high-quality variant dataset. We then leverage this data to demonstrate that it is possible to identify candidate loci associated with fitness-related traits in critically endangered species, which can be used to determine breeding values, and to deploy this data to support conservation management decisions.

Kākāpō are a large, nocturnal, lek-breeding, and flightless parrot endemic to Aotearoa (New Zealand). In the past, these birds had a broad distribution over the three main islands of Aotearoa ^10,11^ (Figure 1a), but the arrival of humans and invasive mammalian predators decimated the species. Early conservation efforts were thwarted by these invasive predators ^12,13^. By 1976, kākāpō were thought to be functionally extinct, with fewer than 15 males present in Ata Whenua (Fiordland; ^14^). But the discovery of a population including females on Rakiura (Stewart Island) in 1977 rekindled conservation efforts ^15^. Most known birds were relocated to predator-free offshore islands to stabilize the rapidly-declining population ^13^. The current population of 252 kākāpō (adults & juveniles as of 10 Aug 2022) is derived from a population of 51 in 1995 which is when the formal Kākāpō Recovery Programme began, marking the beginning of intensive, research-informed management led by the Department of Conservation Kākāpō Recovery Team in partnership with Ngāi Tahu, a tribal group in Te Waipounamu (the South Island of Aotearoa New Zealand) ^16,17^. Only one male kākāpō, named Richard Henry, survived to breed from the Fiordland population. Kākāpō are now confined to five offshore islands where human disturbance is minimized, and invasive mammalian predators are absent (Figure 1a).

Kākāpō have a low reproductive rate owing to multiple factors, including high hatching failure ^18^ attributed to reduced egg fertility and embryo failure ^18^, irregular breeding synchronized with mass-fruiting (masting) of certain tree species such as rimu (*Dacrydium cupressinum*), and small population size ^19^. Intensive management, observation, and the early application of genetic markers and later reduced-representation sequencing, enabled a rich pedigree that has been used for management with the dual aim of minimizing inbreeding and maintaining genetic diversity ^20–23^. Integrating these types of data with management of fitness-related traits remains an unsolved problem ^1,3^, not unique to kākāpō, but provides an opportunity to improve active conservation outcomes.

To enhance the genomic-informed management of this critically endangered species, genomic resources were developed for kākāpō, including a reference genome ^24,25^ and whole-genome sequence data (Kākāpō 125+ Consortium). Reference genomes and associated genomic resources function as living documents, with each new study or analysis providing an opportunity to update the genome, its associated annotation, and the genomic data for each individual ^26–29^. For example, new variant calling methods, especially those that incorporate machine learning models, can be used for increased accuracy ^30–34^.

Dense and accurate genotyping opens the pathway to better inform conservation-focused management. Traditionally, conservation programs for intensively managed species sought to maximize population viability and genome-wide diversity by minimizing mean kinship (Box 1), but some programs are considering the incorporation of functional diversity ^35^. Breeding values (the ‘genetic merit’ of an individual) and genome-wide association studies (GWAS) provide a two-pronged approach for better understanding functional trait variability ^36^. At an individual level, linking genomic variation to variation of key fitness traits and inferring breeding values can provide additional information for prioritizing resources, such as urgent veterinary interventions. Further, it allows for enhanced monitoring of phenotypes such as growth rates based on expected ranges, determined from genetic contributions of parental genotypes (Box 2). At a population level, GWAS can infer the number and effect size of genetic variants that contribute to differences between individuals. This genetic ‘architecture’ significantly impacts small populations’ ability to adapt in response to selection ^37^.

Small sample sizes often prove a challenge for detecting variants associated with phenotypic traits and accurate inference of breeding values ^38,39^. Here, we develop and combine several approaches to overcome and quantify uncertainty in small sample size genomic analyses, which are inherent to critically endangered species, including: i) simulation of phenotypes to optimize methodology for heritability estimates; ii) probabilistic programming to model uncertainty to estimate genotypic and environmental contributions toward phenotypic variance; and iii) cross-validation of breeding values. Although studies of similar-sized populations have proven successful with GWAS (such as in plants), none have been as genetically depauperate as kākāpō ^19,25,40,41^. Crucially, in partnership with the Kākāpō Recovery Team, our foundational models provide predictions that can be tested on and improved by data from subsequent generations as the population increases, and hence are well-placed to be incorporated into the team’s evidence-driven conservation management toolbox. By analyzing the genomes of nearly all living individuals as of 2018, we have developed new approaches that will be used to accelerate kākāpō recovery and can be readily applied to other threatened and treasured species with relict populations.

## Methods

### Data Access and Cultural Considerations

Genomic reads, variant data, and phenotypic data are taonga of Ngāi Tahu and sensitive for the Department of Conservation. Raw and processed data, such as variant files, are available via application to the Aotearoa Genomic Data Repository https://data.agdr.org.nz/.

### Scripts and Workflows

Scripts, workflows, Jupyter notebooks, and other methodology resources are available at the GitHub repo for this paper: https://github.com/GenomicsAotearoa/Kakapo.

### Sample Collection and Sequencing

Kākāpō are currently (October 2022) managed across five predator-free islands: Whenua Hou (Codfish Island), Pukenui/Anchor Island, Te Kākahu-o-Tamatea/Chalky Island, and Pearl Island, located offshore of the South Island of Aotearoa/New Zealand, and Te Hauturu-o-Toi/Little Barrier Island located offshore of the North Island (Figure 1a). Between 2015 and 2018, high-throughput short-read libraries were created for 169 individuals, 125 of which were living at the time. Samples from live birds were collected as part of routine monitoring activities by the Kākāpō Recovery Team. DNA extraction and sequencing was completed as described for modern samples in Dussex et al. ^25^. Sequencing was paired-end with a goal of 30x coverage across three individual sequencing runs, and actual mean of 23x coverage. Mean depth, range, and mean mapping quality after filtering are available in Table S1.

### Variant Calling

Reads were pre-processed using fastp v0.20.0 ^42^ to correct truncated read files and AdapterRemoval v2.2.4 ^43^ to remove sequencing adaptors. Read qualities were checked with FastQC ^44^ via MultiQC ^45^. Reads were aligned to the kākāpō reference genome bStrHab1.2.pri (NCBI BioProject: PRJNA489135, RefSeq Accession: GCF_004027225.2) using bwa ^46^ and saved as CRAM files using samtools ^47^.

We employed DeepVariant v0.9.0 ^33^ to call variants in our dataset. DeepVariant was initially run using the default WGS model. We then utilized single nucleotide polymorphisms (SNPs) identified from confident regions of calls plus a Mendelian inheritance-corrected SNP set, calculated using a custom script, set to train a kākāpō-specific DeepVariant model. Confident regions were identified by base and read quality >= 10 and coverage between 15x and 1000x, inclusive. The set of Mendelian inheritance-corrected SNPs was created by examining loci incompatible with Mendelian inheritance within 16 nuclear families containing both parents and at least three full siblings. Where an offspring genotype was incompatible with the high-confidence calls of both parents, we corrected the offspring genotype. DeepVariant was then run after randomly downsampling confident regions to 100%, 80%, and 75%. Samples were shuffled at each step to prevent overfitting. Training was performed with a learning rate of 0.001 and batch size of 306, using the inception v3 model, starting from the original WGS checkpoint. After ~360k training steps, reads were randomly dropped to keep 100%, 70%, 60%, and 30% of reads to improve lower-depth region accuracy, and training was continued until ~500k training steps. The final accuracy for SNPs and indels were between 93 and 94% (Fig S1). Finally, variants were called for all samples and regions using the kākāpō-specific model. We used the Genomic Variant Call Format (gVCF) format, which provides complete statistics for all genome regions, allowing for additional variants from future generations to be incorporated. The resulting gVCFs were merged with GLNexus v.1.2.2 ^30^.

To determine the performance of our custom DeepVariant model, we calculated final Mendelian inheritance errors from DeepVariant and compared these with Mendelian Inheritance errors from a parallel run of GATK v4.1.4.1 ^48^ (Supplemental Methods, Fig. S2). To account for prediction differences, we filtered on population SNP quality and individual-level genotype quality. Except for the highest level of filtering, our custom DeepVariant model outperformed GATK in each instance.

### Population Statistics

Variants for the population genomic analyses were filtered in VCFtools v0.1.14 to remove indels, invariant sites, select biallelic SNPs, and require >90% of samples to be genotyped at each SNP ^49^. Unless otherwise stated, no minor allele frequency filtering was applied in an effort to capture rare alleles unique to Richard Henry and his descendants. Nucleotide diversity, Tajima’s D, pairwise F_ST_ (calculated in 10kb windows), and heterozygosity (per site) were all estimated using VCFtools v0.1.14. Nucleotide diversity, Tajima’s D, and Heterozygosity were estimated for: all individuals combined, Fiordland birds and descendants (Richard Henry and his descendants), Stewart Island birds (excluding Richard Henry descendants), all Founders in the kākāpō pedigree who were successfully sequenced, all Offspring, and three generations of offspring (Table 1). Offspring generations were determined based on age, with three generations selected for analysis because each had >15 birds that were separated by approximately seven years. Pairwise F_ST_ was estimated between Fiordland birds and Stewart Island birds, Founders and Offspring, Founders and each generation separately, and also between each of the generations. Runs of homozygosity (ROH) for each group were calculated with the BCFtools v1.13-23-g80f6aa8 RoH Hidden Markov model using the filtered SNPs with a minimum minor allele frequency (MAF) of 0.05 and no more than 20% missing, with unlimited buffer size and number of overlapping sites as specified by parameters -b 0 -I ^47,50^.

**Table 1.** Relevant groupings for population statistics in kākāpō. Categories with founding individuals do not have an age mean or range as the age of founders is unknown. (1) Founders are defined here as original birds in the kākāpō pedigree known to have produced offspring and that were sequenced.

| **Population groupings** | **n** | **Mean age (days)**<br>**(age range)** | **Mean Tajima's D** | **Mean Heterozygosity observed (expected)** |
| --- | --- | --- | --- | --- |
| All individuals | 169 | NA | 0.997 | 0.236 (0.236) |
| Fiordland | 8 | NA | 0.921 | 0.399 (0.306) |
| Stewart Island | 161 | NA | 1.760 | 0.228 (0.226) |
| Founders | 35<sup>1</sup> | NA | 0.559 | 0.238 (0.238) |
| Offspring | 128 | 4288 (27 - 14225) | 0.979 | 0.237 (0.234) |
| Generation 1 | 18 | 6561<br>(6548 - 6574) | 1.116 | 0.212 (0.203) |
| Generation 2 | 28 | 3993<br>(3973 - 4014) | 1.349 | 0.221 (0.215) |
| Generation 3 | 30 | 1441<br>(1421 - 1463) | 0.971 | 0.234 (0.231) |

To explore the population structure, we conducted a principal component analysis (PCA) with plink v2.00a3LM based on the SNPs filtered using a minimum MAF of 0.05 and no more than 20% missing calls ^51^. We further examined structure across the genome with local PCA of all individuals (Fig 2, S70, S71) using non-overlapping windows of 450 SNPs, using the python implementation of lostruct, and examining the first 10 PCA dimensions ^52,53^. This method identified 3,062 non-overlapping windows. The distance between all PCA windows was calculated with the get_pc_dists function, with the dimensionality of this 3062×3062 matrix reduced using multidimensional scaling (MDS) analysis from SciKit-Bio using the fsvd method and 6 dimensions ^54^. Across the genome, outliers for MDS dimensions 1 and 2 were identified by examining regions at least 2 standard deviations from the mean.

### Phasing and recombination

SNPs were phased in AlphaPeel v0.1.0 using the pedigree ^55^. Phased outputs for each chromosome in each individual were then filtered on i) having at least 20% of SNPs phased per individual for that chromosome, ii) fewer than 75% of phased SNPs falling in sliding windows classified as uncertain, and iii) average length of segments of consistently phased SNPs at >150 SNPs. Recombination was called on the filtered chromosomes when contiguous blocks of SNPs switched from having a high probability of being inherited from one grandparent to the other. SNPs with a probability of origin from one grandparent >90% were recoded to a binary, and switches in the segment of origin were identified using the changepoint package (ver 2.2.2) in R v3.6.3 ^56,57^.

### Founder Relatedness

As relationships between founders were unknown, the kākāpō studbook (pedigree) was enhanced by incorporating genomic-based estimates of founder relatedness ^35^. The R_XY_ estimator ^58^ was chosen after comparisons with several other estimators for precision with known first-order relationships — including KING ^59^, KGD ^60^, KGD with a correction for self-relatedness (as per ^61^), and TrioML ^62^— as outlined in ^63^. The final SNP data set used to estimate relatedness included a stringent filtering protocol using BCFtools and VCFtools to include only biallelic SNPs, coverage between 3 and 100, a minimum GQ score of 10, minimum allele frequency cutoff of 0.05, and a genotyping rate >90%. This SNP set was additionally filtered for linkage disequilibrium using BCFtools with an r^2^ of 0.8 and a sliding window of 1000 sites, resulting in 8,407 SNPs. Comparisons of relatedness estimators used — and filtering strategies chosen — can be found in the supplement (Fig S3).

### Variant Effects Census

A custom Perl script was used to convert the chromosomal names from the NCBI kākāpō genome to those equivalent to those used in the variant mapping. A custom Python script was then used to make the NCBI kākāpō genome annotation GFF3 file compatible with SnpEff v.1.2.3 64 by removing 138 gene models that were identified as unreasonably long or generated SnpEff errors. Once finished, SnpEff was run on the kākāpō reference genome bStrHab1.2.pri NCBI accession GCF_004027225.2 and NCBI annotation release 101, our SNP sets, and the cleaned GTF file.

### Phenotype Collection and Processing

All data collected during kākāpō management since 1975 are stored in a database held by Aotearoa New Zealand’s Department of Conservation Kākāpō Recovery Programme. Kākāpō are monitored intensively, particularly during breeding seasons, when chicks are given weight and health checks daily to weekly. Adults are captured at least annually for health checks and transmitter changes. The database contains weights, health status, behavioral, and morphological traits recorded during captures, as well as disease and reproductive history of each individual.

Phenotypes examined in this study are: egg shape index (ratio of width to length), clutch size, growth rate during the first 60 days, proportion of fertile eggs, embryo survival, and aspergillosis susceptibility. Save for egg shape index, the phenotypes are important for fitness. Egg shape index averaged across all measured eggs for each mother was analyzed as it is expected to have high heritability and can thus serve as a realistic benchmark of our ability to detect quantitative heritability ^18,65–68^. To capture uncertainty in the small sample sizes for the phenotypes and to provide information to the conservation team regarding factors influencing these phenotypes, we corrected each phenotype for random effects using probabilistic programming, excluding egg shape and aspergillosis susceptibility (Table S1; Supplemental Methods).

For the other phenotypes, important factors to be accounted for were determined using domain knowledge combined with ANOVA using the scikit learn library v0.24.2 ^69^, and manual experimentation to find the best fit model. These factors included the year, location (island site), hand-reared status, and the availability of ripe rimu fruit, a key food source for breeding. Hand-rearing is known to cause a change in growth rates ^70^. Rimu fruit is the preferred food for chicks, but supplemental feeding of nesting females is known to have an effect on growth rates ^68,71^. Generative models for each of these phenotypes were fitted using variational inference and implemented in Tensorflow Probability 0.14.1 utilizing Tensorflow 2.8.0 ^72,73^, or the Tensorflow model API fit function, minimizing the negative log-likelihood of the model. Chick growth to 60 days was fitted to a Gompertz curve following the method of Crispim et al. ^74^ with the predicted mean of three variables utilized in downstream analyses (body weight at 60 days; *M*, growth speed; *a*, and the growth-rate coefficient; *b*) ^74,75^. For all remaining phenotypes, the predicted means of the final distribution of corrected phenotypes were used in downstream analyses. The specifics of methods for each phenotype can be found in the supplemental materials, and code is available for each phenotype in Jupyter notebooks ^76^.

### Phenotype Heritability Simulations

The kākāpō population is small, genetically depauperate, and highly structured, and this is reflected in the genomic variation. We expect these features to have affected the allele frequency and effect size distribution in our population. As a consequence, we conducted simulations to determine the most appropriate methods to use and the utility of each for our heritability analyses. We simulated quantitative phenotypes based on the observed kākāpō SNP calls. Phenotypes were simulated across a range of heritabilities and a number of causal SNPs using R code as suggested in the hibayes package ^77–81^, modified for our use. The hibayes package was then used to run heritability analyses across all Bayesian Alphabet methods. We also calculated heritability for the simulated datasets using a TASSEL 5 kinship-based mixed linear model (with P3D option on for computational efficiency) ^82^. These simulations support BayesC and TASSEL as the most appropriate methods for heritability estimates for our dataset (Supplemental Materials: Heritability Simulation). Results are plotted in Figures S4 - S6.

### Phenotype Heritability, Association, and Breeding Values

Heritability was calculated for our corrected phenotypes using TASSEL (MLM model with P3D off) ^83,84^ as well as with Bayesian Alphabet methods implemented in hibayes. We conducted genome-wide association analysis using the package TASSEL 5. Phenotypes, as processed above, were analyzed in TASSEL 5 using a mixed linear model (MLM) with P3D off, with minimum observations of 50 samples per site and MAF of 0.02. Putatively associated genes were chosen from the genes found in the top 1,000 SNPs per trait followed by a literature search.

Breeding value parameter optimization and cross-validation was performed using KAML using weight at 60 days (*M* parameter of the Gompertz curve) from our processed dataset (covariates, environment, removed; Supplemental Methods). Finally, we calculated genomic-based breeding values for alle processed phenotypes with KAML using the optimized parameters. GWAS QQ and Manhattan plots, as well as predicted breeding values, are found in Figures S8 - S68, and supplemental tables S3 - S12.

## Results

### Population Diversity

In total, we identified 2,102,449 SNPs and 417,571 indels across 169 sequenced birds. Filtering for less than 20% missing called sites results in 1,923,224 high-quality SNPs and 8,325 indels. Further filtering for minimum allele frequency (MAF) of 0.05 results in 1,257,633 high-quality SNPs and 7,334 indels. The SNPs of the filtered callset have a Mendelian inheritance error rate of 1.28%. Ts/Tv ratios for filtered and unfiltered variant sets were 2.18 and 2.16, respectively. Many Richard Henry-lineage (n=8) specific alleles are lost with MAF filtering, thus, most analyses exclude the MAF filter.

Global PCA analysis was performed for the population as a whole and for the founders using the MAF filtered dataset (Fig. S69, S70). As expected, the primary outlier from the entire population is the sole Fiordland bird, Richard Henry, on PC1, which accounts for 38.2% variance explained for founders, and 43.5% for the entire population. Jean and Barnard, two additional founders, are the outliers on PC2 (accounting for 12.8% variance for founders, and 10.5% for the entire population) (Fig S7). Local PCA analysis revealed 3 clusters of outlier regions (> 2 s.d. of mean) on MDS1, clustered on chromosome 2, 5, and 7. MDS2 contained 3 clusters as well, on chromosome 2, 6 and 7 (Fig. 2).

The dataset excluding MAF filtering was used to estimate population genomic statistics. In the latest generation (Gen 3, as determined by youngest age), heterozygosity was up slightly from that of the founding population (mean 0.234 vs 0.233) and higher than in previous generations, with a mean of 0.212 and 0.221 for Gens 1 and 2. Tajima’s D ^85^, useful for comparing the number of heterozygotes to the number of segregating sites, increased for all offspring compared with the founders, but decreased in subsequent generations (Table 1). Runs of homozygosity (ROH) are a useful metric to assess demographic history and types of inbreeding ^86^. For example, inbreeding events that occurred more distantly in a geneology are expected to have shorter runs of homozygosity than recent inbreeding events, as recombination reduces the length of ROHs over time ^87–89^. Using the MAF filtered dataset, founders have a mean ROH length of 442,092 bases (306 - 115,478,914 bases). Offspring have a mean ROH length of 419,093 bases (371 - 73,089,505 bases).

Locations of putative recombination sites were identified by following haplotypes through the pedigree. After filtering for phasing quality, 4257 chromosomes (excluding microchromosomes, due to their small size) in 169 birds were scanned for recombination. A total of 2969 recombinations were identified. These occurred in the highest density at the ends of chromosomes and overlapped with the regions of greatest SNP density. Recombinations were also observed to occur at the borders where local population structure changed (Fig. 2).

Relatedness of the original founding individuals of the population are seen in Fig S6. While average relatedness amongst founders was low (average ± SD = 0.03 ± 0.06), our approach detected 37 relationships between founders with an R > 0.125 (first cousins), and two pairwise estimates approximating first-order relationships (R=0.5, parent-offspring or siblings).

### Variant Effect Census

Using the filtered variant calls, SnpEff identified 509 high-impact (0.012%), 29,273 moderate-impact (0.693%), and 52,269 low-impact (1.237%) variants. Of these, 58.7% were silent mutations, 0.2% were nonsense mutations, and 41.0% were missense mutations. The unfiltered variant set contained 5,692 high-impact (0.065%), 55,494 moderate-impact (0.637%), and 91,373 low-impact (1.049%) variants. The missense to silent ratio for filtered and unfiltered was 0.699 and 0.776, respectively.

### Phenotypes

Kākāpō phenotypes were pre-processed to account for possible confounding factors, with heritability assessed using Bayesian Alphabet methods and a mixed linear model (MLM), run through GWAS as an MLM, and breeding values estimated. Phenotypes were fitted using probabilistic programming to fit the models with uncertainty, and the predicted mean individual effects were used in the downstream analyses.

The phenotypes we assessed consisted of egg shape index, chick growth, aspergillosis susceptibility, clutch size, embryo survival, and proportion of fertile eggs. Heritabilities ranged from 0.093 to 0.158 via BayesC, and ~0 to 0.754 with TASSEL. Additional heritability estimates from other methods are available in Supplemental Table 2.

**Table 2.**
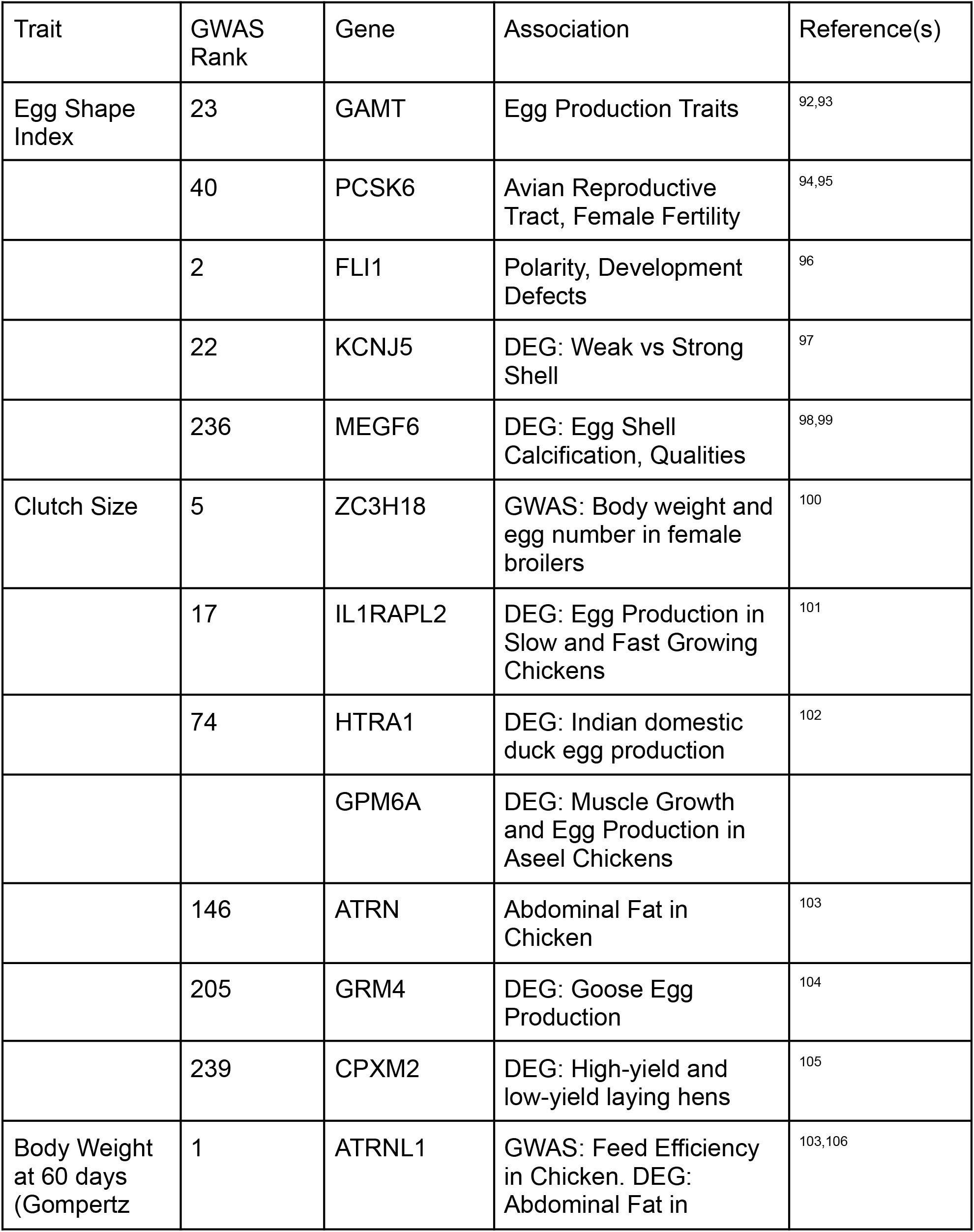
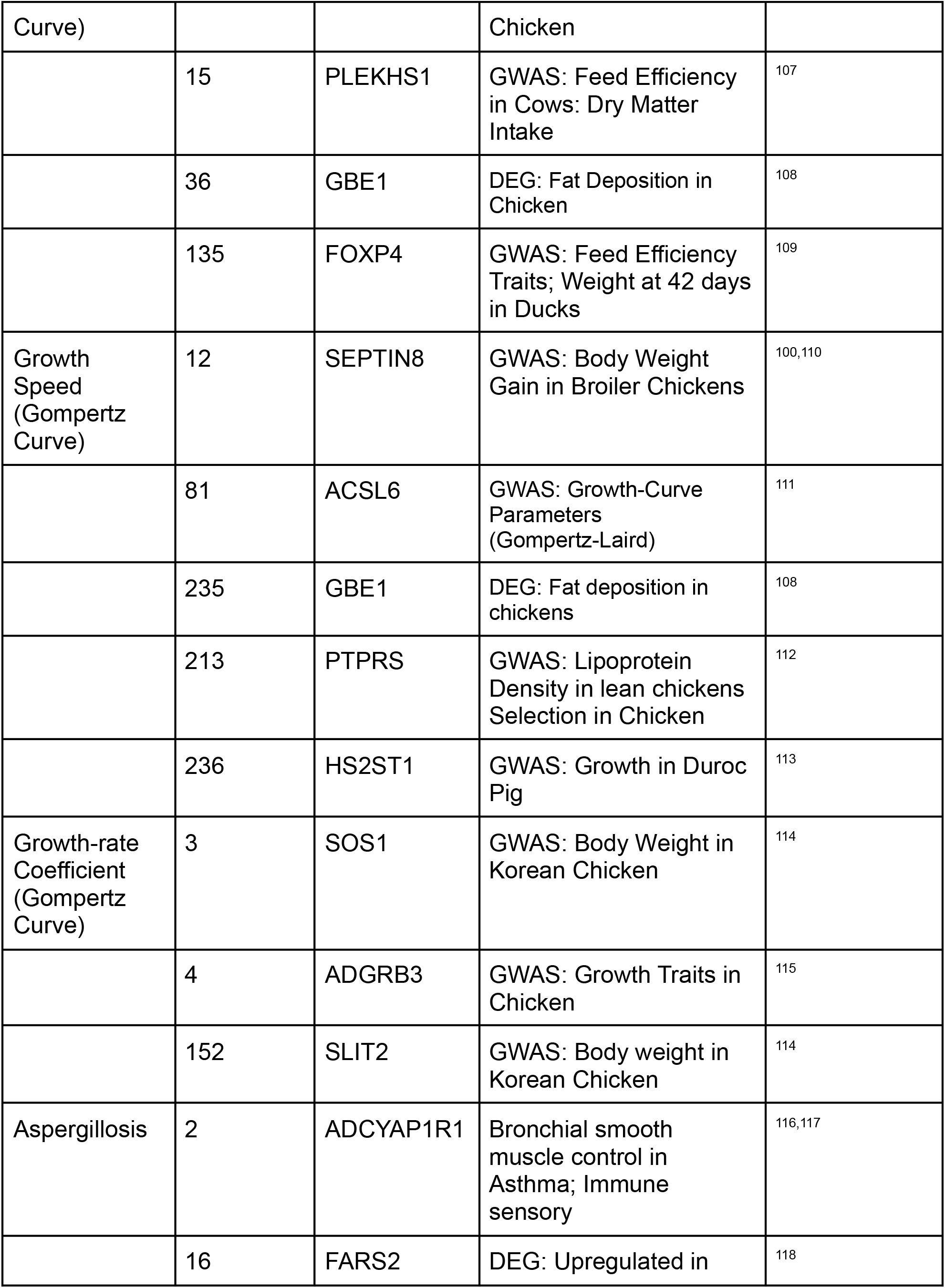
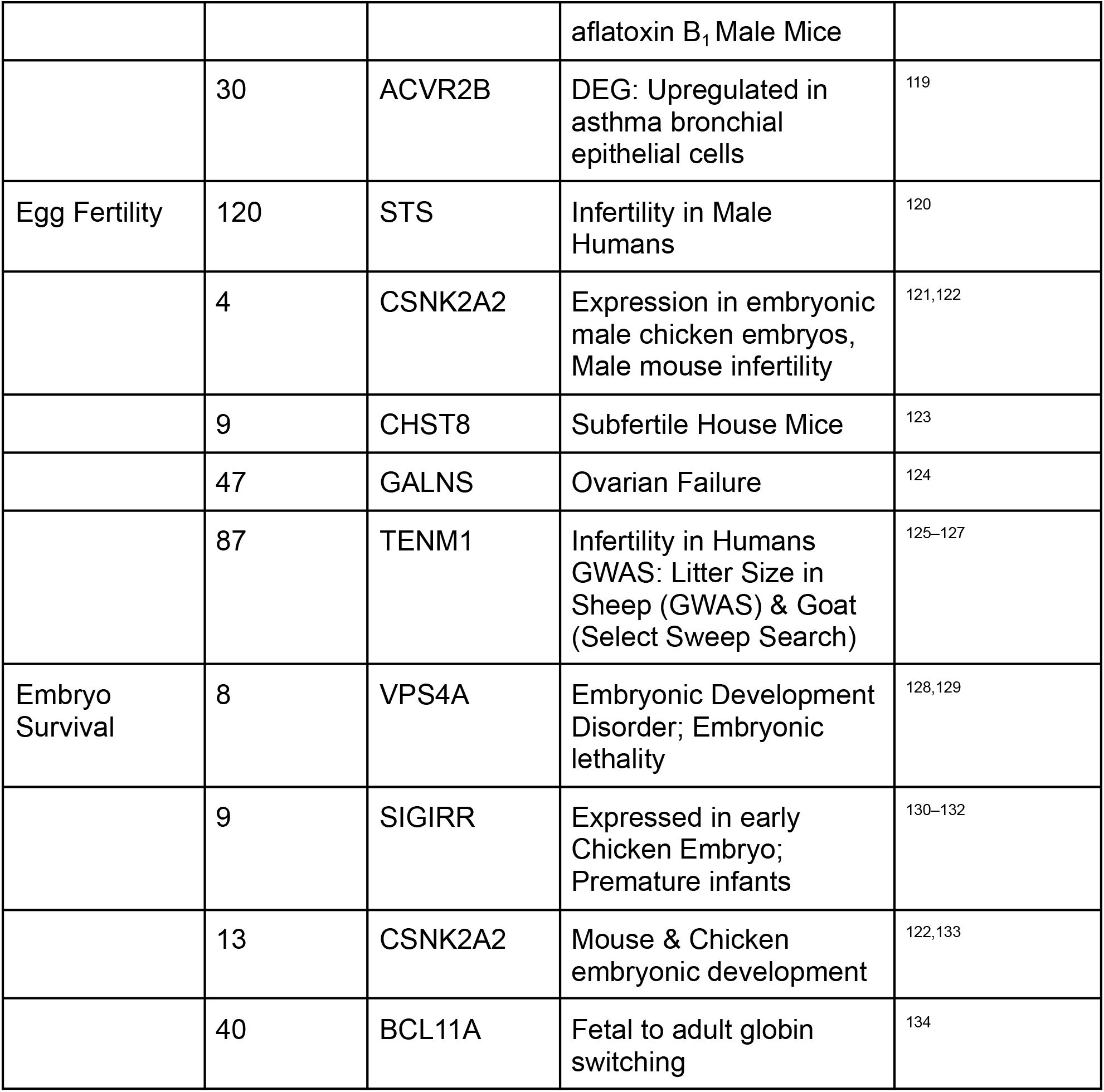
Selected genes from genome-wide association studies (GWAS) peaks associated with the phenotype of interest in kākāpō. ‘Association’ describes the association between the same gene and a related trait in another species from GWAS or differentially expressed gene (DEG) analysis.

Our GWAS analysis identified regions of the genome associated with variation in all traits. For each GWAS peak, representing a series of markers rising above the background -log10 p-values, we identified candidate genes close to the most highly associated markers. A summary of some genes of interest identified from these GWAS peaks is presented in Table 2, along with literature supporting possible relationships between the gene and phenotype in other species. Identifying genes putatively contributing to a phenotype from GWAS can lead to false associations ^90^, especially in small, highly-structured populations ^91^. However, we are confident that our dataset is enriched for true positives as we have identified loci that have been identified in other related species and independent experiments, including GWAS but also differential expression and quantitative trait locus (QTL) mapping, to be associated with the same or similar traits. The top 1000 GWAS results can be found in supplemental tables S5-12.

Genotype and phenotype data was then used to generate breeding values to support management decisions (Figure 3, Box 1). The breeding values of the founders compared to the subsequent generations (specified in Table 1) are generally consistent, with a similar mean and variance over time. Some traits show a slight depression in the oldest offspring generation, with the mean of the trait’s breeding values returning towards the original founder mean by the more recent generation (Supplemental Figures Sa-Sb). For example, body weight at 60 days has a mean of 0.500 for the founders, 0.417 in the oldest offspring generation, 0.415 in the following generation, and 0.469 in the youngest sequenced generation. The same effect is seen in egg shape index, with mean breeding values of 0.524, 0.500, 0.550, and finally 0.524. Descendants of Richard Henry, 7 in total, have different average breeding values compared to the rest of the population. This includes lower 60-day body weight (t-test p-value 0.09) and a lower egg shape index. Interestingly, Richard Henry’s descendants have higher fertile egg ratios and surviving embryos than the rest of the population, although they are often mated with unrelated birds, which may represent an interplay between the genetics of both parents.

To illustrate an example of our approach for each phenotype, chick growth for the first 60 days of life was fitted to a Gompertz curve, a three-parameter equation often used in animal studies. The Gompertz curve fits the body weight (here the 60 day weight, as parameter *M*), the time-scale parameter (*a*), and the growth rate (*b*). Cofactors were fitted for the effects of island, year, island by year, hand-reared status, and ripe rimu fruit availability. These variables explained ~73% of the total variance in the dataset. The remainder was then fitted per bird, which left 6.61% remaining mean absolute percentage error. The per bird contributions to the three growth parameters were run through heritability estimators, GWAS, and breeding value calculations. All three traits provided peaks on the Manhattan plot (Supplemental Figures S24-S44).

By examining GWAS peaks and some of the genes underlying them, we identified genes of interest associated with kākāpō chick growth and found supporting literature to give confidence to our results. Our peaks include genes associated with body weight gains in chicken GWAS (ATRNL1), body-weight growth in chicken GWAS (SEPTIN8), a known growth-established QTL in chicken (HS2ST1), lipoprotein density QTL (PTPRS), a gene differentially expressed between both high and low-fat chickens, and also fast and slow-growing chickens (GBE1), and a gene associated with backfat thickness in pigs (MKLN1) (Table 2). The intersection of our results with other studies support our GWAS results as likely enriched for true positives, and thus an association to the trait, despite our small, inbred population.

## Discussion

By leveraging species-scale genomic resources, we have created a high-quality variation callset for nearly the entire extant kākāpō population. This foundational building block will inform conservation specialists in ongoing management decisions, enable assessment of the efficacy of those decisions, as well as contribute more generally to our understanding of the impact of population bottlenecks on threatened species recovery. This trained DeepVariant model proved superior to GATK SNP calling, increasing the number of high-quality SNPs while more than halving Mendelian inheritance errors (from 3.38% to 1.28%). The quality of this variant dataset is critical to the analyses presented here and maximizes the accuracy of the information when analyzing genomic data for threatened species. Further, the trained model can be utilized for future generations of kākāpō, increasing the accuracy of variation calls without needing to train the data again.

Kākāpō have been under intensive conservation management using microsatellite and pedigree resources. By examining genome-wide diversity and inbreeding through subsequent generations of kākāpō, we illustrate that conservation management decisions based on pedigree data have successfully kept inbreeding to a minimum and maintained genetic diversity. Because pedigrees assume that founders are unrelated to one another, there is potential for pedigrees to underrepresent empirical relatedness between extant individuals ^35^. As demonstrated here, high-quality genomic variant data continue to inform pedigree resources by inferring founder-relatedness values, including first-order relationships between founders within the kākāpō pedigree. Since 2020, this genomic-informed pedigree has been used to inform the translocation of kākāpō among island populations to maximize genetic diversity for each. We anticipate this informed pedigree will continue to enhance translocations, strategic mate pairings, and/or artificial insemination of kākāpō in the future. Further, identifying recombination hotspots and linkage disequilibrium (Figure 2) allows for fine-grained simulations of future populations. Such simulations allow the accurate prediction of genetic diversity and inbreeding in future individuals, identify when and where new allele combinations may occur, and provide the basis for modeling the impacts of future conservation interventions on genetic diversity.

The most significant input into the genetic diversity of kākāpō is provided by Richard Henry, the single source of the Ata Whenua/Fiordland lineage and diversity in the remaining population. Previous research has suggested that Richard Henry carries a higher load of deleterious variants ^25^. Using our high-quality SNP dataset combined with the updated NCBI RNA-Seq-backed annotation, many previously identified deleterious loss-of-function variants are no longer detected (Tables S13-S15). This conclusion is related to the quality of the annotation used in that analysis and, to a lesser extent, the SNP calls used. The genetic variation brought into the kākāpō population through Richard Henry is thus likely valuable to the species’ long-term future. Because of this finding, we believe that high-quality annotation is crucial to accurately identify mutational load for downstream purposes. As the newest annotation is underpinned by only two tissue samples for RNA-Seq analysis, further RNA-Seq and reannotation are likely to improve inference for future kākāpō studies. The efficacy of proxy statistics for mutational load, such as Genomic Evolutionary Rate Profiling (GERP) scores ^135^, is unknown in managed populations, and these proxies are often isolated from fitness related traits ^136,137^. Richard Henry carries slightly higher mutational load as determined by GERP scores, but we show that he also carries values associated with higher fitness and fertility.

Importantly, genomic analysis can identify genetic variants associated with key fitness traits. These traits provide predictions for the genetic merit of individuals based on their genome-wide data. Here, we used probabilistic programming to identify trait associations while accounting for uncertainty generated by the environment and management interventions that contribute to phenotypic variance. This approach allowed us to gain maximum information from the small sample sizes inherent to studies of endangered species. We identified associations to genes contributing to fitness traits previously identified in chickens and other species, giving credence to the idea that our phenotypic studies are valid. Our phenotypic analyses have provided the ability to make predictions for key traits for future birds based on the pedigree, which will provide useful information to the conservation team to help with the prioritization of limited resources. For example, phenotypic prediction of health outcomes can prioritize early intervention and monitoring for birds at high risk without necessarily performing any selection or breeding. Crucially, these predictions can be made based either on genomic information for an individual or by inference from the genomic information from its parents. We envisage that this prediction framework will provide valuable insight as to when to allocate limited resources for illness and increased monitoring in the event of disease outbreaks.

## Conclusion

We have cataloged the majority of the genomic diversity in the kākāpō population and addressed and understood the limitations created by small sample sizes. With this data, conservation researchers and practitioners for kākāpō can co-develop approaches for better monitoring phenotypes and genomic statistics of interest. For example, key fitness traits can be more effectively monitored, including identifying birds at high risk for aspergillosis and inferring when individuals have abnormal growth rates, while expected values for parents can be used to better predict future offspring traits in the absence of offspring sequence information. The genetic variation described here can be used to identify regions of interest for future analysis projects and imputation of new generations, reducing sequencing costs as populations expand. This phenotypic and population genomic resource that accounts for fitness traits can be managed with approaches to minimize inbreeding to enhance both demographic rates and evolutionary potential in this species. Future work could fully simulate approaches to model potential management decisions accounting for phenotype, genotype, inbreeding, and other concerns which could be derived from this data.

Genomic sequencing can benefit threatened species conservation programs by helping identify loci of interest for traits, gauge the success of genetic diversity preservation, and resolve hidden relatedness. High-throughput sequencing, along with high-quality phenotypes and pedigrees open up additional avenues for future research projects, and the breadth of data processed here supports future and ongoing projects looking after the wellbeing of a threatened species. We highlight the value of viewing the genome and associated genomic resources, including sets of variants, as living ‘documents’ that can be continually improved upon and used for increasingly accurate predictions.

This project brings new information and context to an ongoing breeding program, updating results from previous work in this field ^21,22,25,67^. Overall the conservation program is successful, not just in population increases but also in maintaining genetic diversity. From the variation data presented here and the phenotypes collected over many years, we have developed routes that can help improve the management of kākāpō, and be repurposed for use in other species of similar conservation concern. Information gained from identified genes and loci can be used to pursue new methods of non-genetic intervention. New sequencing data and phenotypes can increase the accuracy of prediction of these techniques, but a crucial next step is to discuss with conservation managers, agencies and Indigenous guardians the use of information generated by these tools for the benefit of endangered species.

## Box 1

**Revealing founder relatedness to better inform conservation translocation decisions**

In kākāpō, conservation translocation decisions are made to ensure that all offshore predator-free island populations are healthy (e.g., managed for diseases which affect kākāpō, such as cloacitis), productive (e.g., appropriate sex ratio, age, hand-reared status), functional (e.g., behaviorally compatible individuals), and genetically diverse (i.e., maximized genome-wide diversity). Until recently, incorporating genetic diversity into conservation translocation decisions has been done using a pedigree validated with microsatellite-based parentage assignments ^21^. However, one shortcoming of most pedigrees–including the kākāpō pedigree–is that founders are assumed to be equally unrelated ^35^, leading to underestimated relatedness values. While attempts have been made to use microsatellites to estimate founder relatedness ^20^, microsatellites have been shown to produce poor estimates of relatedness in genetically depauperate species ^61,138^, like kākāpō.

Worldwide, many conservation programs struggle to incorporate founder relatedness into pedigrees ^139^, but thanks to the long-lived nature of kākāpō and the sustained efforts of the Kākāpō Recovery Team, genomes for almost all kākāpō founders were available, and we were able to use the high-density SNPs developed in this manuscript to estimate relatedness among 35 founding individuals, representing all but a singular breeding member of the original population. In 2020, these estimates were incorporated into the kākāpō pedigree to enable better informed decisions using mean kinship and the Mate Suitability Index (MSI) in PMx ^140^. This new genomic-informed pedigree provides the Kākāpō Recovery Team with more confidence to undertake conservation translocations, including the transfer of 11 females to Te Kakahu in 2020 to establish a new breeding island (Figure 2), and the identification of another 64 birds for transfer before the next breeding season.

**Figure 1.**
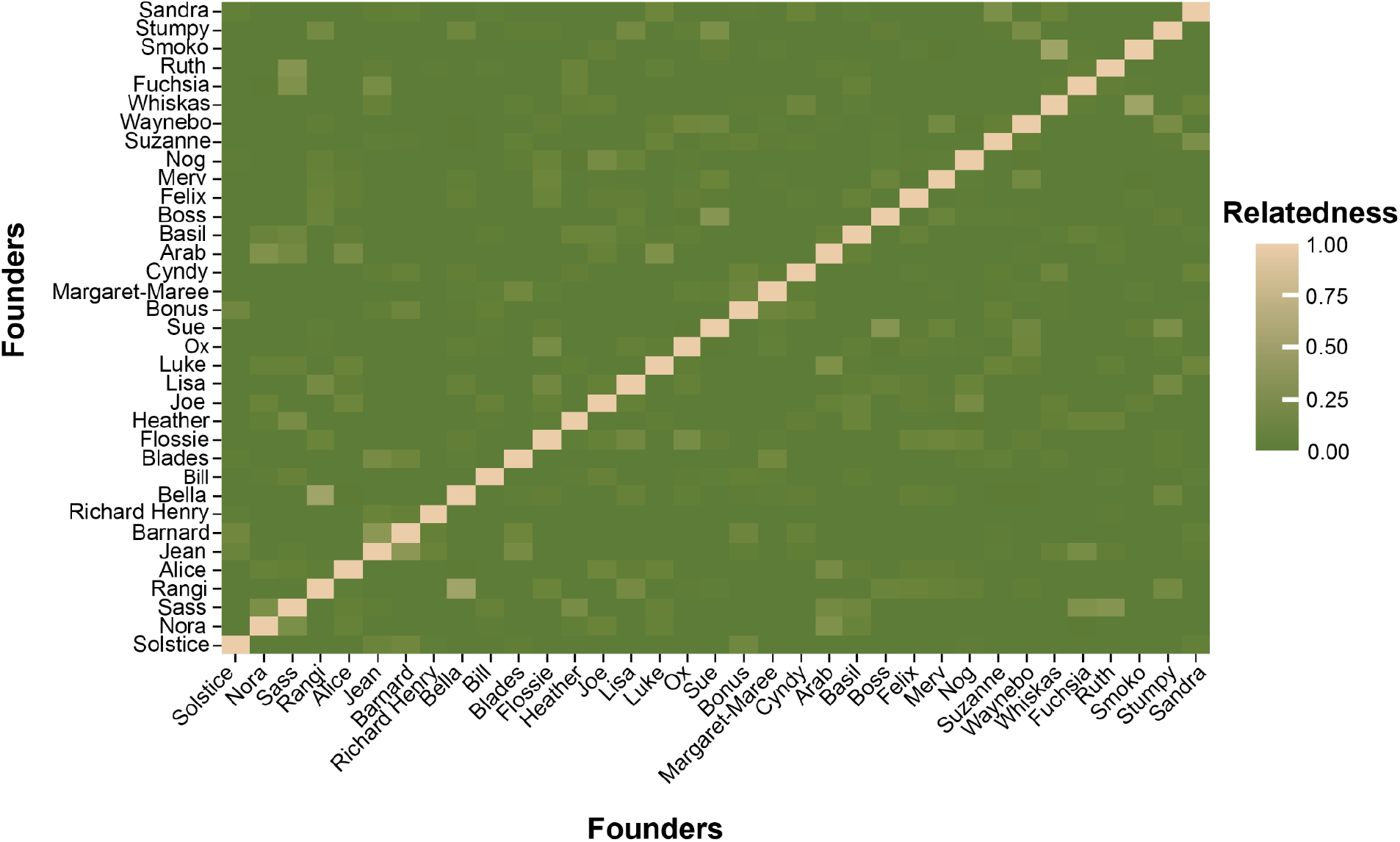
Heatmap of relatedness among 35 kākāpō founders, with relatedness being represented in scaled colors from unrelated (dark olive green) to more related (tan). The diagonal represents self-relatedness (*R* = 1).

**Figure 2.**
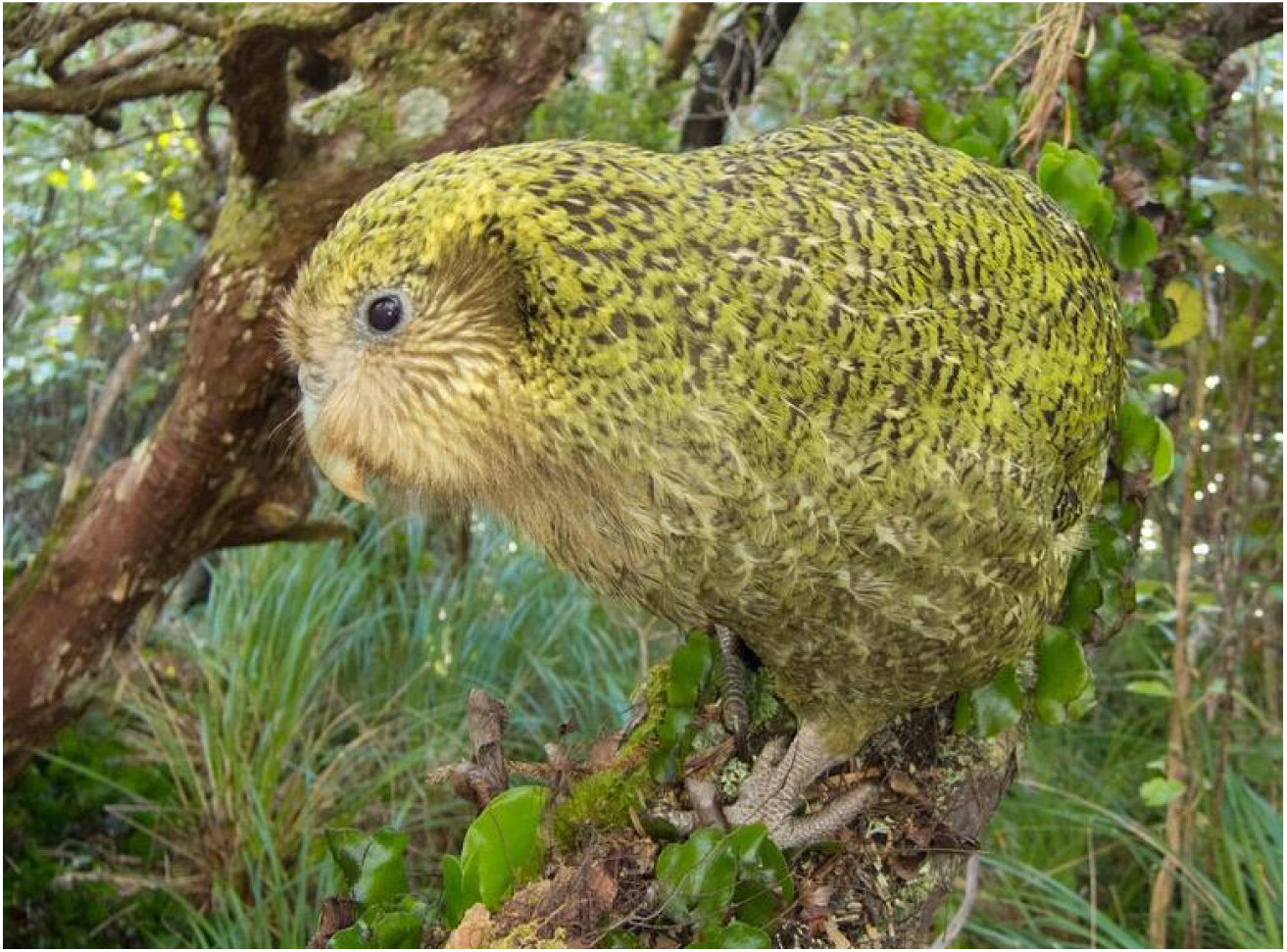
Image of Mahli, a female kākāpō that was translocated to Te Kakahu in 2020, thanks to the genomics-informed pedigree. Andrew Digby / New Zealand Department of Conservation

## Box 2

**Monitoring Growth for Early Vet Intervention**

Critically endangered, with breeding only occurring every 2-5 years and with an unusually long lifespan, each kākāpō chick is a valuable opportunity to increase the population. For this reason, eggs are frequently taken from the nest to be incubated until hatching ^70^, followed by close observation either in hand-rearing or in a natural nest to identify any medical issues or difficulties requiring vet care.

**Figure 1.**
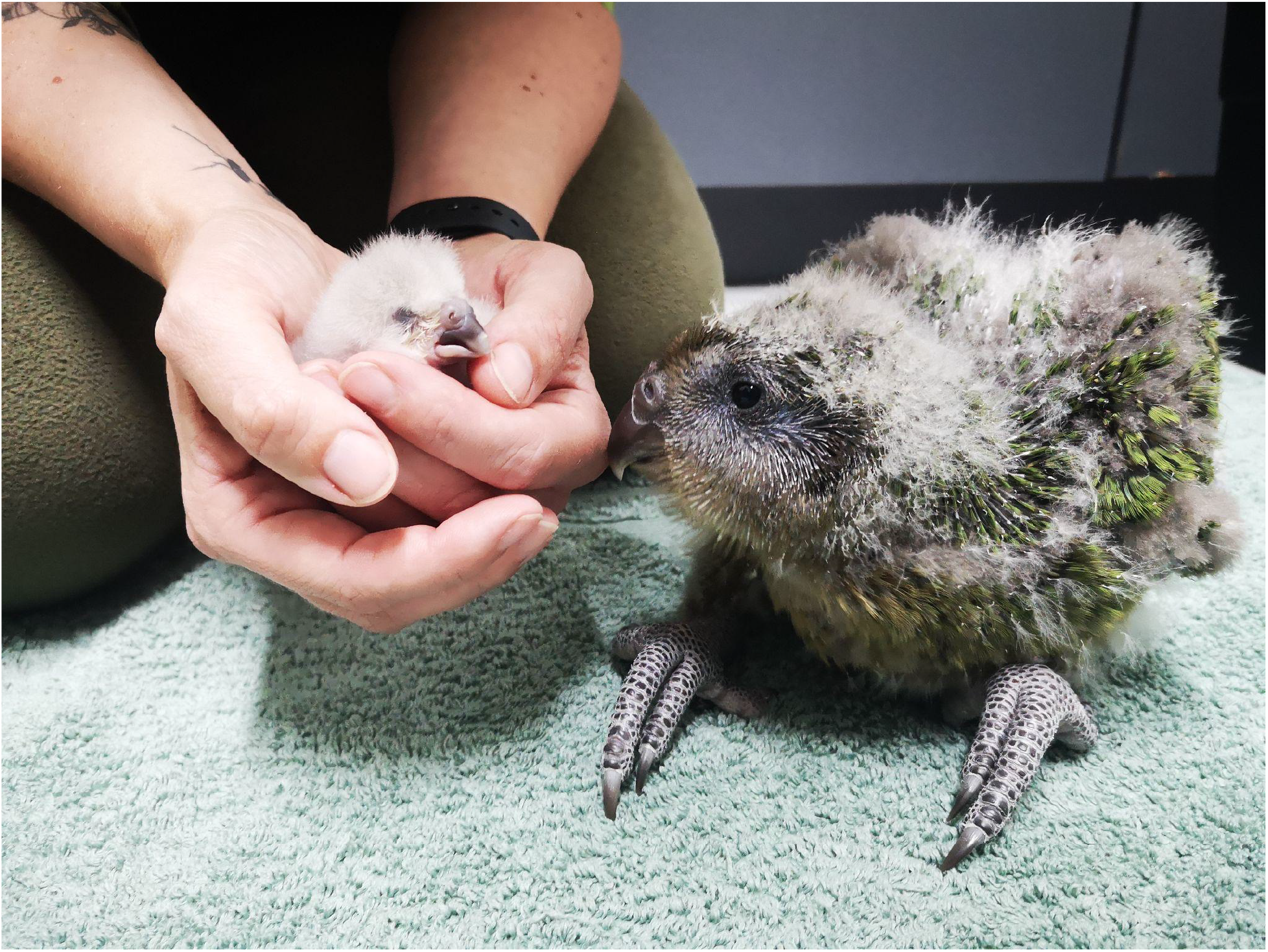
Close monitoring of chick weights assists conservationists in identifying individuals that may need additional monitoring or health assessment. Zephyr-A2-2022 (left), 2 days old, and Pearl-A2-2022 (right), 38 days old. Photo credit: Lydia Uddstrom

One promise of quantitative genetics is predicting traits from genotype, allowing conservation practitioners and veterinarians to identify deviations from the expected outcome and to apply early intervention if required. Offspring inherit trait values from their parents, and quantitative traits are expected to be close to the average of both parents, although traits are also affected by the environment. While these expected trait values, termed ‘breeding values’ in production systems, are often used to select ideal candidates for breeding, we can also use them to monitor expected growth more closely. Any chicks who grow abnormally from their expected growth rate, either faster or slower, can be prioritized for medical assessment and support. Given the tendency of many species to hide signs of illness as long as possible, having tools that allow for earlier intervention can assist conservation practitioners and veterinarians in giving the best chance for a chick to grow to adulthood.

Kākāpō chick growth in the first 60 days of growth can be modeled by a 3-parameter Gompertz curve. As the population increases, management intensity must decrease. Using parental informed growth curves, it is possible to reduce management for chicks that are doing well and check on them less frequently. Even in the absence of sequencing information on the chick, by utilizing the average of parental breeding values, and estimates of environmental factors, we can plot an expected growth rate alongside the actual growth rate for each chick, and this can be provided immediately to the Kākāpō Recovery Team.

**Figure 2.**
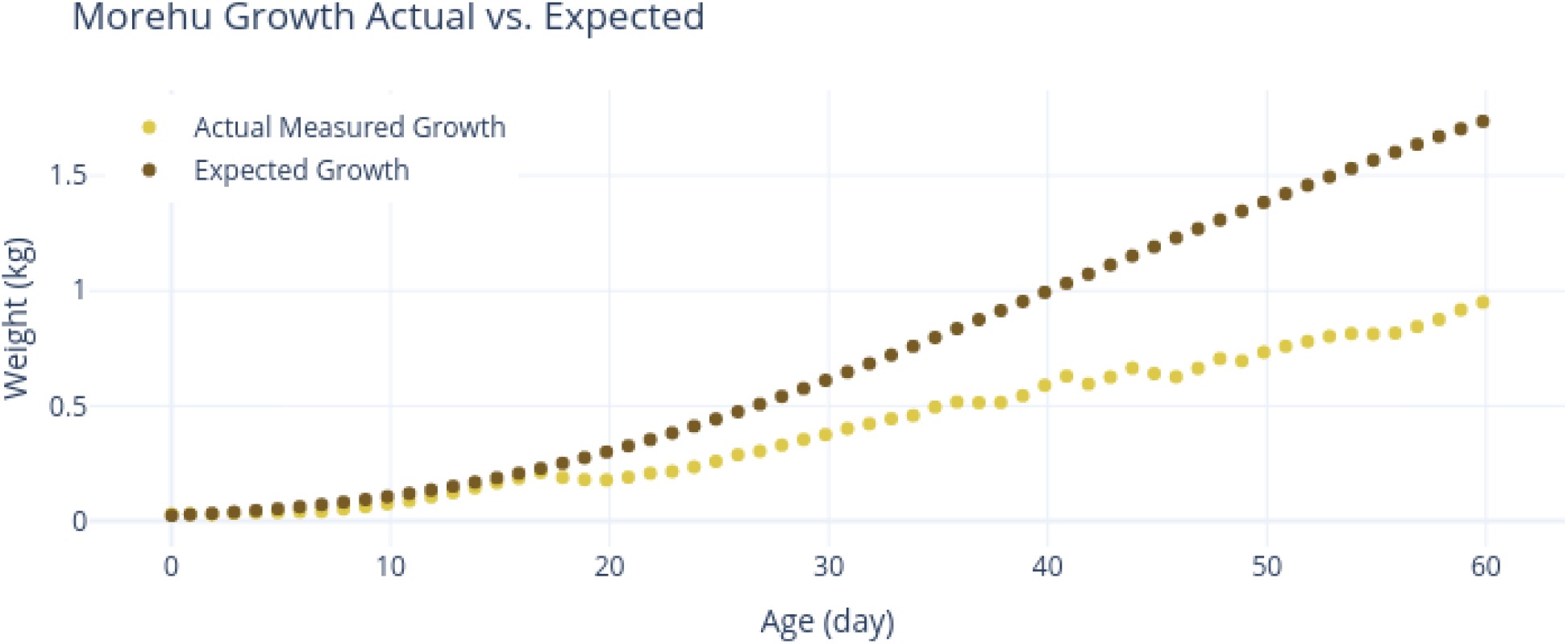
Growth curve of Morehu, a chick determined to be ill. Expected growth is derived from average parental phenotypes. Obvious illness began at day 18, however slow growth can be seen in this chart starting at day 4 and continuing until the more sudden drop at day 18.

**Figure 1.**
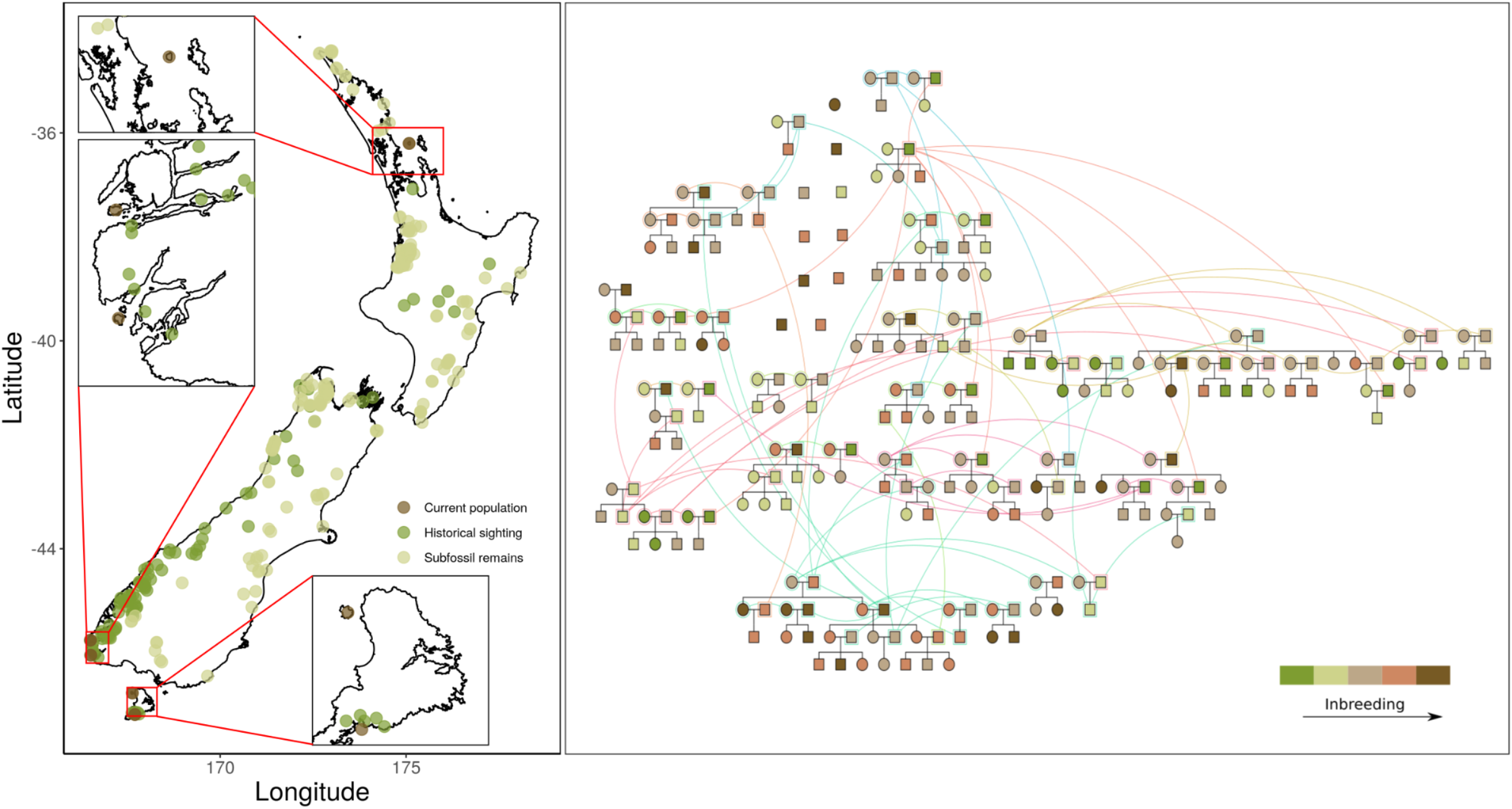
a) Historical and modern population distributions; Kākāpō are currently limited to five, predator-free islands. b) The pedigree of our sequenced population, with coloring according to inbreeding.

**Fig 2.**
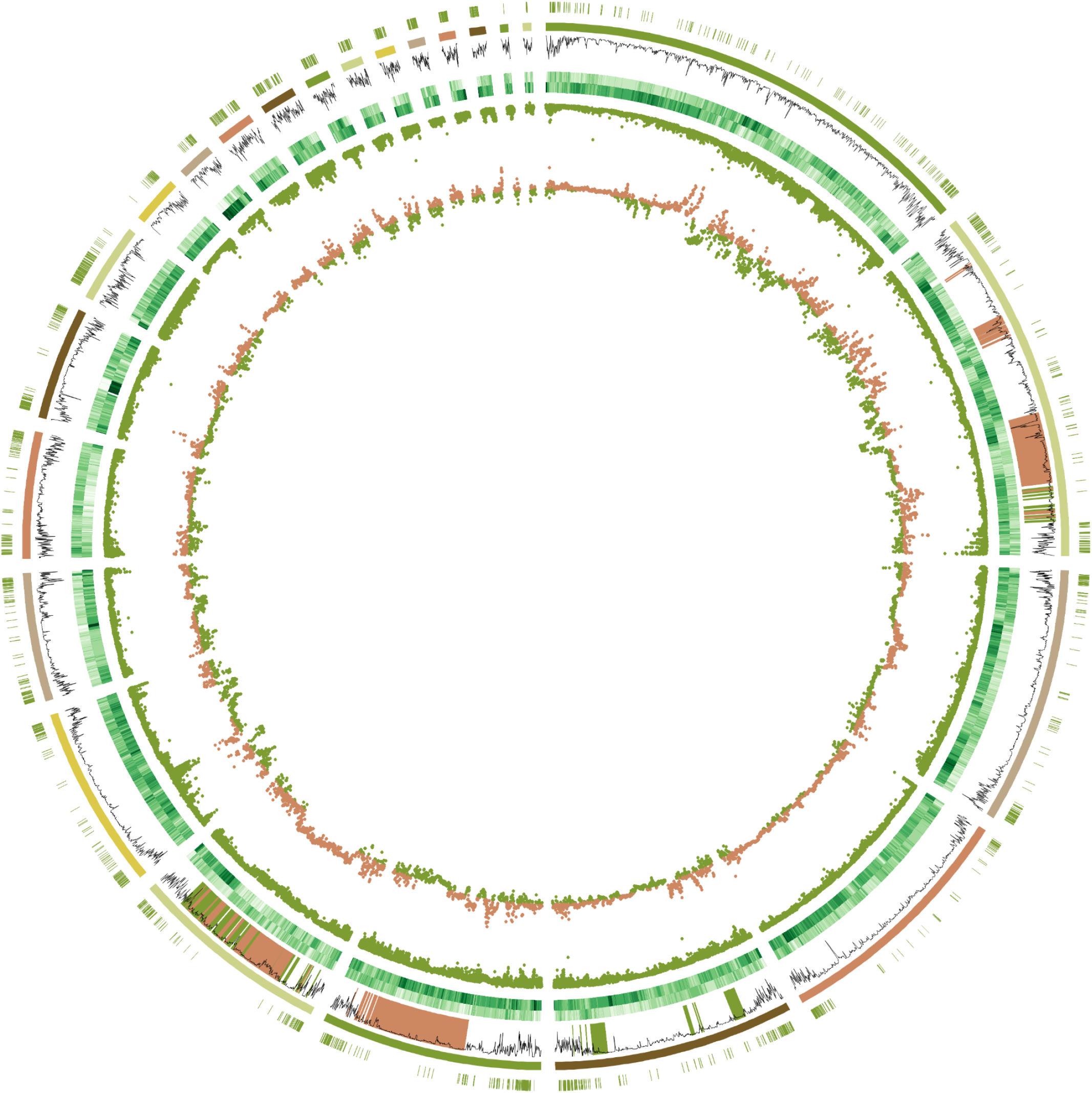
CIRCOS Plot of kākāpō population genomics. From outer to inner, the rings are as follows: the outer ring are locations of putative recombinations, followed by rings representing chromosomes, with Z and W sex chromosomes removed, with the largest being chromosome 1 (Table S2). The line plot is single nucleotide polymorphism (SNP) density per 100kb. Regions of extreme genetic structure are highlighted green (MDS1) and orange (MDS2). The dual-stacked green rings are runs of homozygosity for the founders (outermost) and offspring. The scatter plot in green following are the Fst values plotted for founders vs. offspring. The next scatter plot (orange and green) are heterozygosity scores for founders compared with the newest generation of birds in this analysis. Orange indicates higher rates of heterozygosity in that region in the founders, while green indicates higher rates in the offspring (Generation 3).

**Fig 3.**
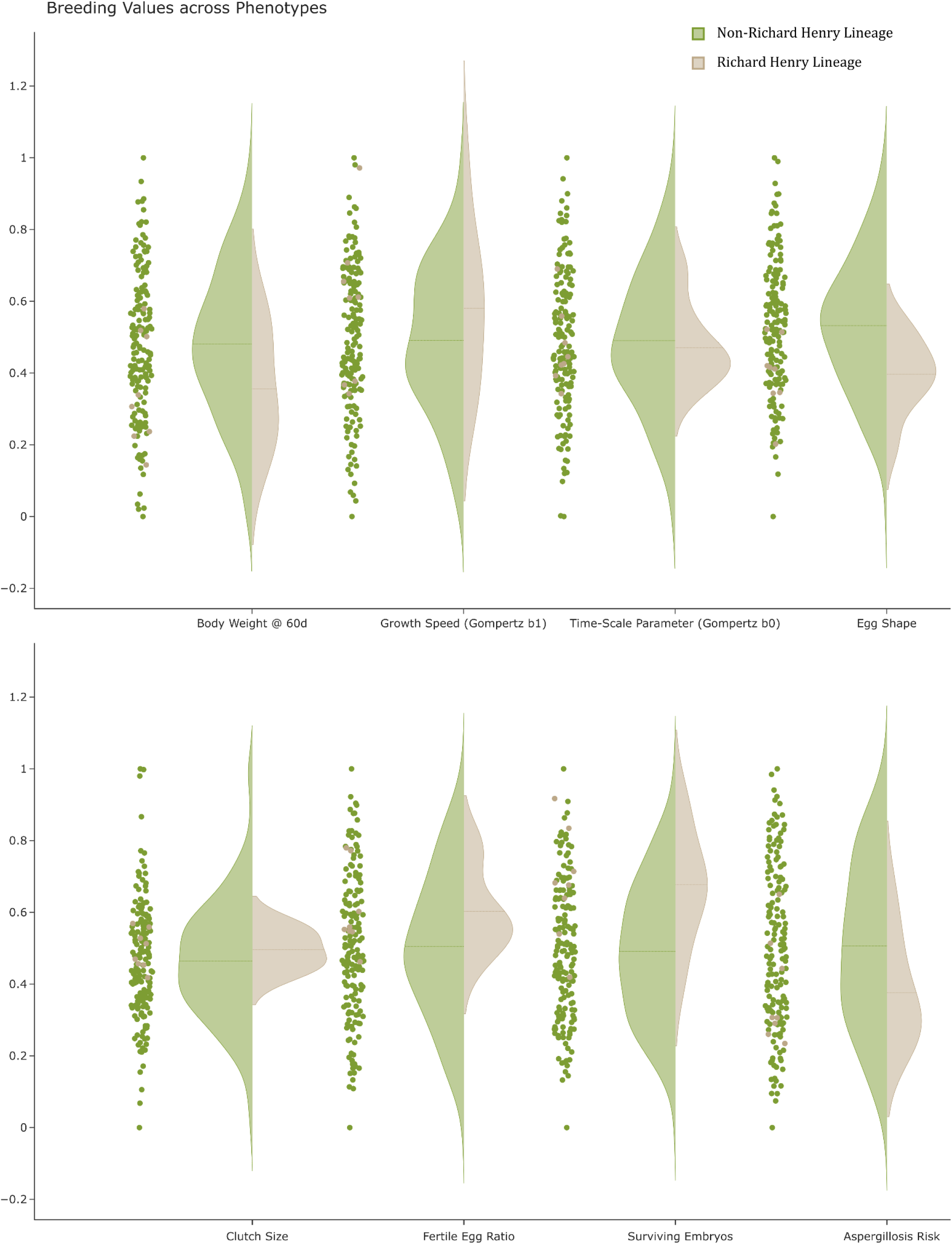
Violin plot of all calculated Breeding Values for each trait. Green represents all birds exclusive of Richard Henry and his lineage, while tan represents Richard Henry and lineage scores (n=8). Aspergillosis Risk is a risk score derived from case-control (see Supplemental Materials: Aspergillosis Susceptibility for details).

