## Supplemental Materials for "Species-wide genomics of kākāpō provides transformational tools to accelerate recovery"

#### DeepVariant Training

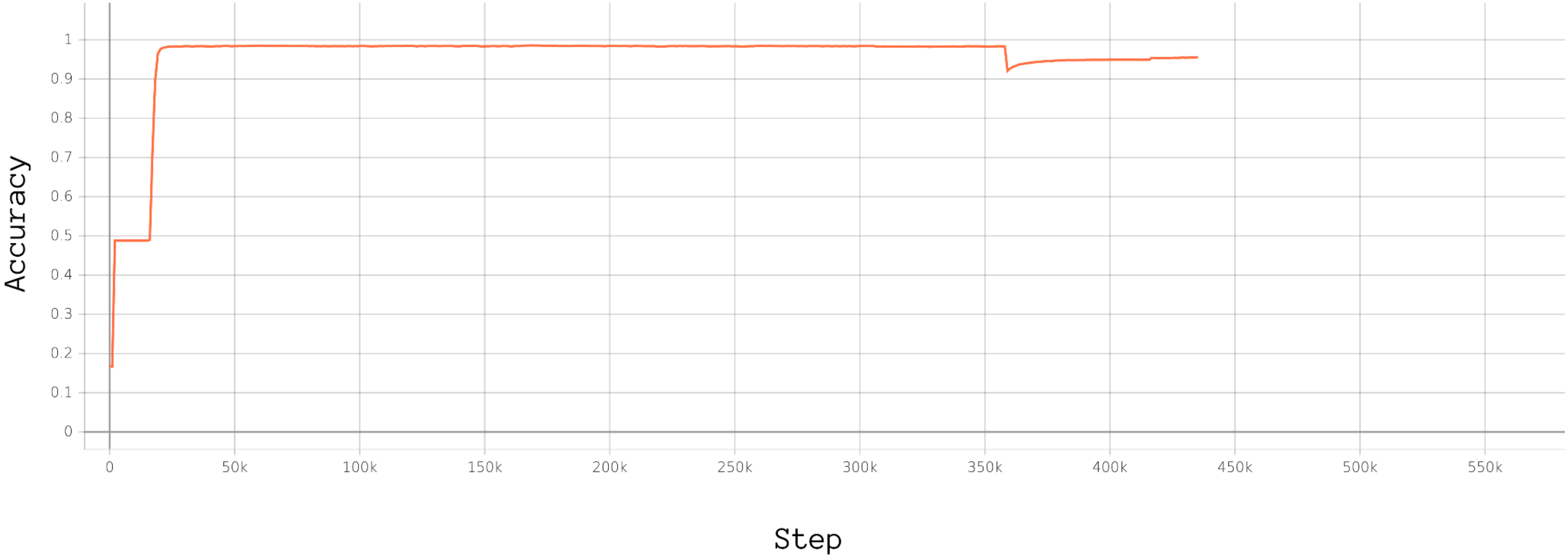


Fig S1. Accuracy of SNPs and indels across training batches for a trained DeepVariant model for kākāpō. The training was interrupted near 360k, and the downsampling was altered to improve SNP calling in low-coverage regions. See main text for details.

### Variant Calling Models

In total, three variant calling pipelines were tested, DeepVariant, which produced the final variant call, FreeBayes v1.2.0-dirty, and GATK 4.1.4.1. Initial quality checks for Mendelian inheritance error and genotype quality scores determined that FreeBayes performance was much lower than for DeepVariant and GATK, and so results from this pipeline were not used further. GATK was used to individually call variants on the full-read set. GATK genomic VCFs, which include statistics for each genomic position, were joint-called and combined using GLNexus; results were comparable but had a higher Mendelian inheritance error rate (3.38%) than DeepVariant (1.28%, Figure S2).

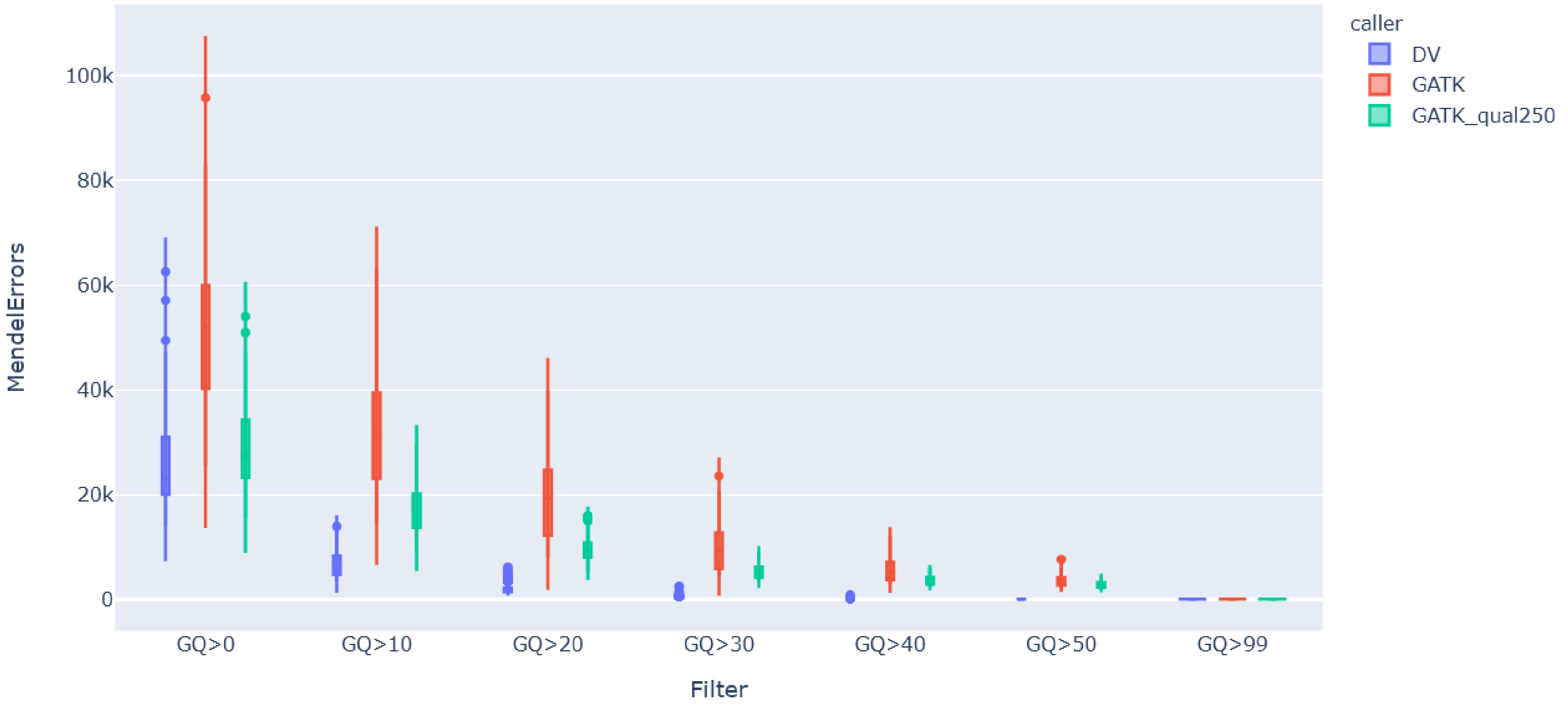


Fig S2. Mendelian inheritance errors were categorized by genotype quality (GQ) to compare a trained DeepVariant model for kākāpō, GATK, and GATK filtering on QUAL scores >= 250.

### Relatedness Evaluation

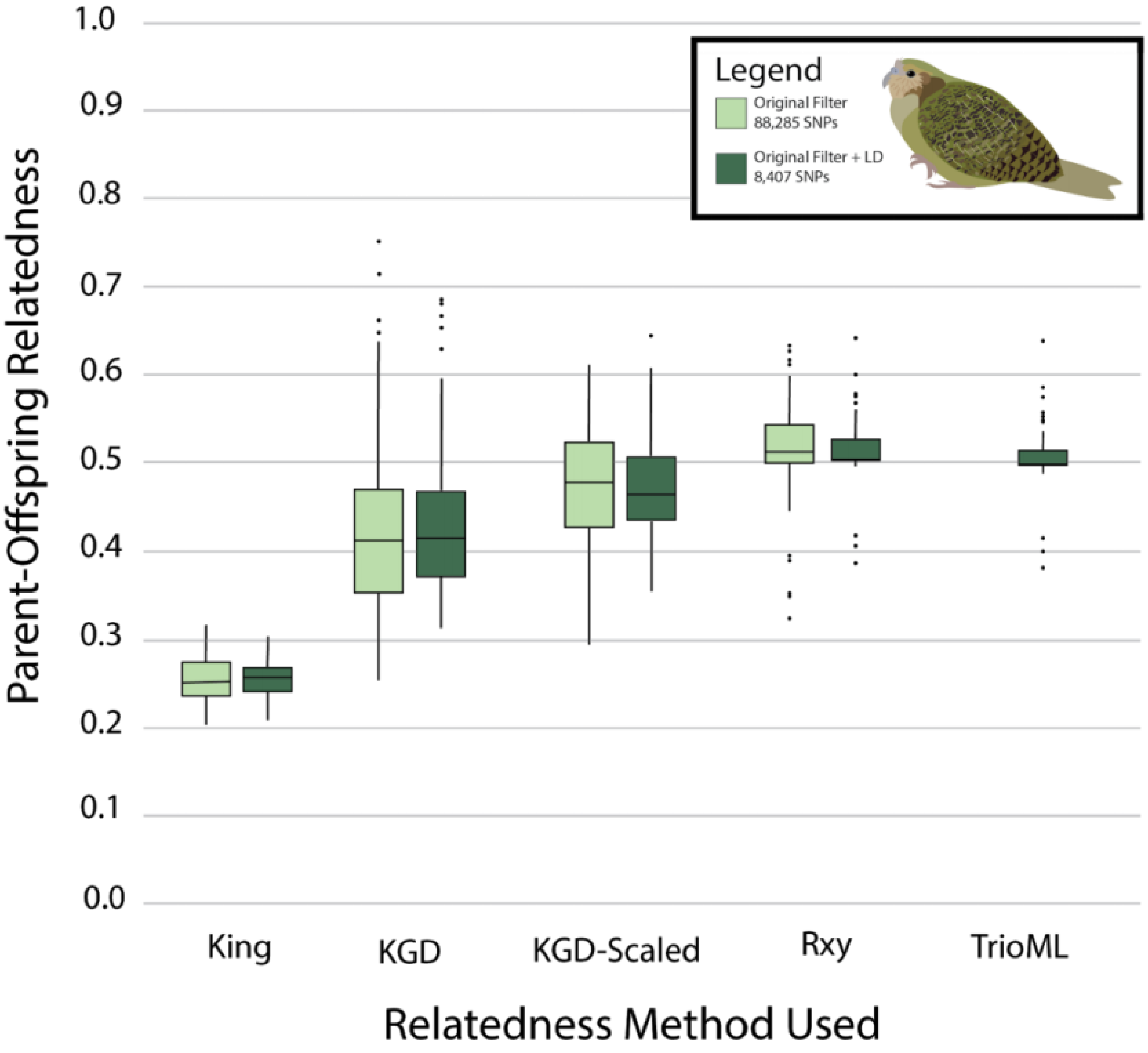


Fig S3. Comparison of known parent-offspring relatedness for kākāpō for multiple relatedness estimators.

Multiple relatedness estimators were tested to identify the best method for accurately measuring parent-offspring relatedness to update the studbook and calculate founder-relatedness, including KING (Waples *et al*. 2019, estimated through the package NGSrelateV2, Hanghøj et al. 2019), KGD (Dodds *et al*. 2015), KGD with a correction for self-relatedness (as per Galla *et al*. 2020), $R_{xy}$ (Hendrick & Lacy 2015, estimated through NGSrelateV2), and TrioML (Wang 2007, estimated through the R-programme *related*, Pew et al. 2015). Mother-offspring relationships (n=91) were used to evaluate the precision of relatedness estimators, as first order relationships between parents and offspring should be 0.5 and mother-offspring relationships were confirmed through both field observations and genetic verification. Rxy and TrioML most closely matched our expectation of parent-offspring relatedness of 0.5, with little variance in the estimate, but due to computational costs, Rxy was chosen as the best method for our dataset.

### Heritability Simulation

We assessed the accuracy of heritability estimates from Bayesian Alphabet methods and from TASSEL by simulating phenotypes with a range of heritabilities with different numbers of causal SNPs. Simulations were performed on the kākāpō genotypes using simulated phenotypes with R code modified from the hibayes tutorial, as seen in our GitHub repository: https://github.com/GenomicsAotearoa/Kakapo.

These simulations used the default number of iterations and burn-in (20,000 and 14,000, respectively). However, in our phenotypic analysis (Table S2), we set these parameters to 500,000 and 20,000. Simulations consisted of test (one of the available BayesAlphabet functions implemented in hibayes, along with exporting these results and running in Tassel with P3D on and off), simulated heritability ("0.01", "0.05", "0.10", "0.20", "0.30", "0.40", "0.50", "0.60", "0.70", "0.80", "0.90", "0.95", "0.99", "1.0"), and number of SNPs controlling phenotype (10, 100, 1,000, 10,000). For each combination of heritability and number of SNPs, we repeated the simulation 10 times, using a different random number generator for each replicate. The performance of each of the methods was quantified via the error and absolute error of the simulated versus estimated heritability.

#### Results

After inspection, BayesC and TASSEL's heritability estimates are used in the main text, as they offer the highest accuracy (Fig S4-5) while benefiting from both Bayesian and traditional analysis. In addition, neither appear to suffer extreme error bias at high or low heritabilities (Fig S6).

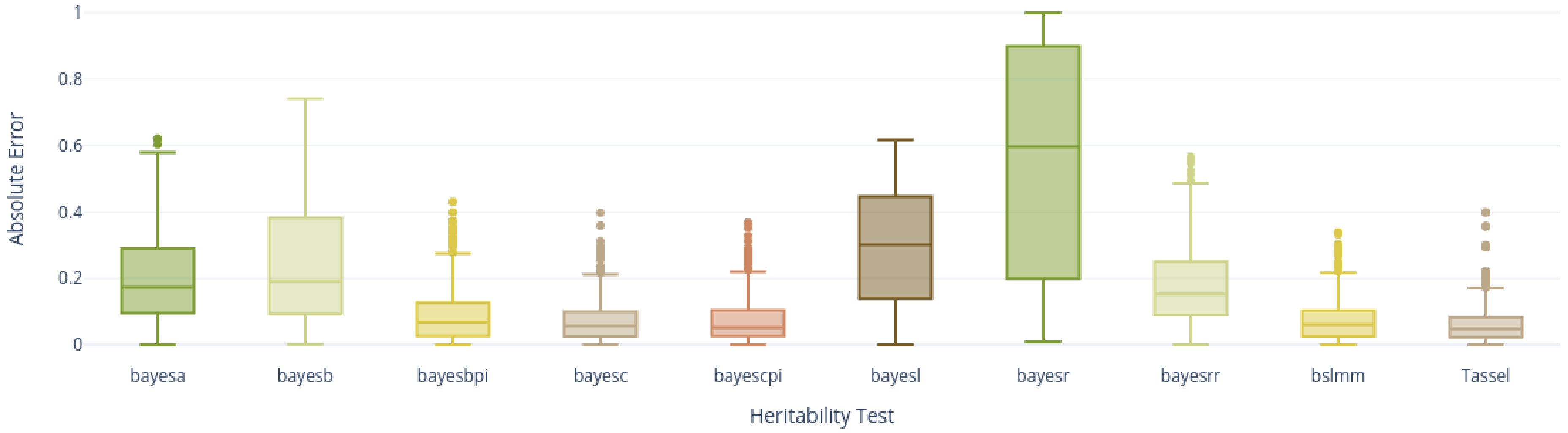


Fig S4. Absolute error by test.

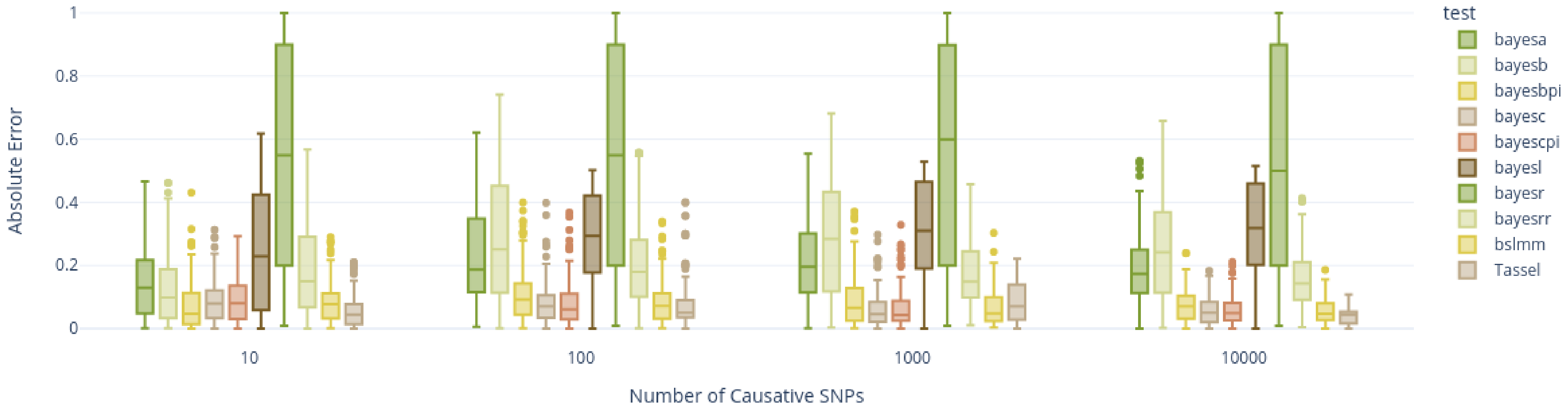


Fig S5. Absolute error by number of causative SNPs (k) per test.

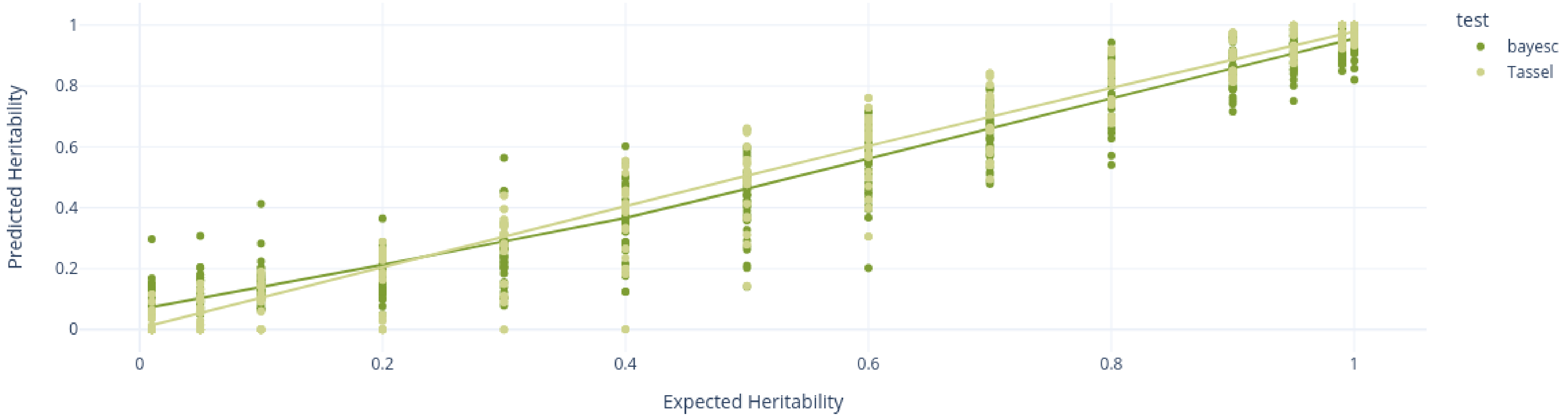


Fig S6. Expected vs Predicted heritability for BayesC and TASSEL. A Lowess trendline was fitted for each test.

### Breeding Value Calculation Optimization

Simulations were run for 10 replicates, with a 10% chance of a phenotype being masked (unknown phenotypes were not masked). We optimized the parameters for our dataset by testing the following: i) KAML cross-validation number (crv.num), ii) variance components estimation (vc.method), iii) top-SNP correlation threshold (cor.threshold), iv) reference GWAS toggle (ref.gwas), v) GWAS number of principal components used as covariates in the model (gwas.npc). Simulations were measured as the correlation between input values and predicted values of the masked phenotypes. Ideal parameters for our population were: crv.num = 10, vc.method = brent, cor.threshold = 0.90, ref.gwas = FALSE, and gwas.npc = 0.

### Phenotype Analyses

Jupyter notebooks are available on our GitHub repository: https://github.com/GenomicsAotearoa/Kakapo
Additional heritability estimates are available in supplemental table S1.

#### Egg Shape Index

Egg shape index was calculated as width/length, and the mean of these ratios was taken for all measured eggs of each mother. Ratios were 0 centred by subtracting the mean from the ratios. There were a total of 759 observations of egg ratios, with 58 mothers with recorded data. TASSEL predicted a heritability of 0.61, while BayesC estimates heritability was 0.49. Egg Shape Index QQ-plots, manhattan plots, and predicted breeding values are available in Figs S8 - S14.

#### Clutch Size

Clutch size phenotypes were available for 48 individuals in our population dataset with confirmed paternity, spanning the years 1981 - 2021 and included 122 entries, 103 of which were first clutches and 19 were second clutches. We adjusted the raw clutch size phenotypes for each breeding event for contributing factors and for repeated measures across a female's lifetime to give a single adjusted phenotype value for each individual for downstream analyses. To do so, we used TensorFlow probabilistic programming, which enables priors on the mean (termed the 'effect') and standard deviation (termed the 'scale') of the impact of each factor on the phenotype to be updated until the model converges. The model output is a set of distributions of the effects (and standard deviations thereof) for each of the factors on the phenotype. The model was trained with variational inference, which enables variation to be shared across different levels of each effect, which is important in datasets such as this one, where some categories of each factor have very few observations.

In consultation with the Kākapō Recovery Team and by exploring the impact of other possible factors on the raw phenotypes (i.e., via ANOVA - see script: Chick Weights Formatting.ipynb), as well as experimenting with models to find the best fit, the final model fitted was:

**Priors**

$$\begin{aligned}
\mu &\sim Uniform(-0.01,\ 0.01) \\
\sigma &\sim Softplus(Uniform(0.01, 1.0)) \\
output_scale &\sim Softplus(Normal(\mu, \sigma)) \\
rimu_effect_r &\sim Normal(\mu, \sigma) && \text{for r} = 0,\ 1 \\
clutch_effect_c &\sim Normal(\mu, \sigma) && \text{for c} = 0,\ 1 \\
bird_effect_b &\sim Normal(\mu, \sigma) && \text{for b} = 0 \ldots 48 \\
intercept &\sim Normal(2.9016395,\ 0.7289093)
\end{aligned}$$

**The Model**

$$\begin{aligned}
v &\sim intercept + bird_effect + riperimu_effect + clutch_effect \\
y &\sim QuantizedDistribution(Normal(v, output_scale))
\end{aligned}$$

**Where**

$$\begin{aligned}
r &= \text{rimu unripe (0) or ripe (1)} \\
c &= \text{first (0) or second clutch (1)} \\
b &= \text{mother bird, } 0 \ldots 48
\end{aligned}$$

The intercept priors on mean and standard deviation were $\mu$ = mean clutch size (2.902) and $\sigma$ = standard deviation (0.729) of the dataset.
Other prior means are uniformly drawn from random samples between -0.01 and 0.01.

The model was trained as outlined in the Jupyter notebook with loss function defined as the target log probability. The mean loss of the final 1000 steps was 150.319. Convergence was reached when log probability no longer decreased.

A single adjusted phenotype value was taken for each individual as the mean of the individual effect distribution from the model above. This phenotype was then used to estimate trait heritability and for GWAS and breeding value analyses. The heritability of the processed phenotypes was 0.18 from TASSEL and 0.16 from BayesC. Heritabilities estimated with additional methods are listed in Supplemental Table 3.

Predicted model variables are found in Figure S15, with the effects of each individual bird in Figure S16. Genome-wide association analysis (GWAS) outputs, including QQ Plots and Manhattan plots, are found in Fig S17 - S22. Predicted breeding values are presented in Fig S23.

#### Growth Rate

Growth rate of chicks was fitted to a Gompertz curve for the first 60 days. Weights were fitted from 8,308 observations, measured from the years 1992 to 2019, for 157 birds, 11 different locations (referred to as islands), and included information on hand-rearing or not. Ripe Rimu was excluded from the dataset as this was captured by Island x Year effects (specified below). Twenty percent of observations, selected randomly, were kept out of the analysis to provide a validation dataset.

We used TensorFlow probabilistic programming to first infer the per-day scale (i.e., standard deviation) across chicks to capture the increasing variance in growth as the chicks get larger over time (in days). To do so, the standard deviation of growth was fitted to a line parameterized by 3 trainable variables (vl[2-4]) and the cumulative distribution function of a logistic distribution parameterized by 2 trainable variables (vl[0], vl[1]). The final formula is as follows:

**Variance Day Curve**

**Priors**

$$v_z \sim Uniform(0.0, 2.0) \quad for\ z\ =\ 0 \ldots 4 \tag{2.1}$$

**Model**

$$\begin{aligned} L &\sim Logistic(v_0,\ |v_1|) \\ f(x) &\sim v_4 * abs(v_3 + v_2 * P(L <= x))] \end{aligned} \tag{2.2}$$

**Where**

$$\begin{aligned} v_z &= \text{Trainable vector of variables} \\ x &= \text{Age, in days} \\ P(L <= x) &= \text{Cumulative Distribution Function for} \\ &\quad\ \text{distribution L at point x (day)} \end{aligned}$$

The formula was fitted to reduce the mean absolute percentage error, and trained with the Adam optimizer.

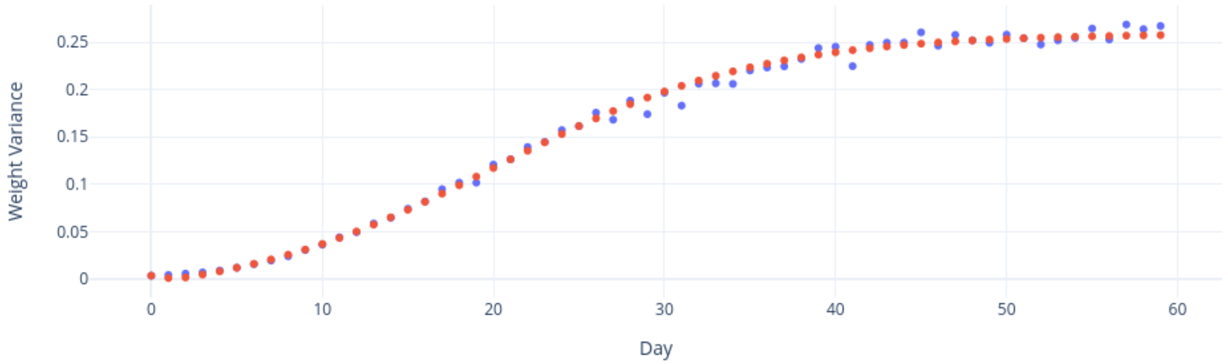


Fig. S7 The blue dots indicate the actual per-day variance, the red dots is the computed formula.

Once the per-day scale was calculated, the following model was then fit to adjust individual growth for contributing factors across an individual's lifetime to give a single adjusted phenotype value for each individual for downstream analyses. To do so, we used TensorFlow probabilistic programming, which enables priors on the mean (termed the 'effect') and standard deviation (termed the 'scale') of the impact of each factor on the phenotype to be updated until the model converges. The model output is a set of distributions of the effects (and standard deviations thereof) for each of the factors on the phenotype. The model was trained with variational inference, which enables variation to be shared across different levels of each effect, which is important in datasets such as this one, where some categories of each factor have very few observations.

In consultation with the Kākāpō Recovery Team and by exploring the impact of other possible factors on the raw phenotypes (i.e., via ANOVA), as well as experimenting with models to find the best fit, the final model fitted was:

#### Fit Intercepts

**Priors**

$$
\begin{aligned}
r &\sim Uniform(0.1, 0.2) \\
\sigma_r &= 1e-8 + Softplus(r) \\
a_{intercept} &\sim Normal(r,\ \sigma_r^2) \\
b_{intercept} &\sim Normal(r,\ \sigma_r^2) \\
M_{intercept} &\sim Normal(r,\ \sigma_r^2)
\end{aligned} \tag{2.3}
$$

**The Model**

$$
\begin{aligned}
w &= M \exp(-\exp(a - bx)) \\
y &= Normal(w,\ o)
\end{aligned} \tag{2.4}
$$

**Where**

$$
\begin{aligned}
x &= \text{Age, in days} \\
w &= \text{Weight, as defined by Gompertz} \\
o &= \text{Output of (2) with day as input}
\end{aligned}
$$

**Fit Effects**

**Priors**

$$
\begin{aligned}
&\text{for v in } a, b, M: && \text{For each variable in the Gompertz curve} \\
&\quad \text{for } p = 1 \ldots P: && \text{for each random-effect group} \\
&\qquad \text{for } c = 1 \ldots |C_p|: && \text{for each category of group } p \\
&\beta_{vpc} \sim Normal(r,\ \sigma_r^2)
\end{aligned}
\tag{2.5}
$$

**Model**

for $i$ in 1..N : For each data point i

$$
\begin{aligned}
M_i &= M_{intercept} + \sum_{p=1}^{P} \beta_{M,C_p(i)} \\
a_i &= a_{intercept} + \sum_{p=1}^{P} \beta_{a,C_p(i)} \\
b_i &= b_{intercept} + \sum_{p=1}^{P} \beta_{b,C_p(i)} \\
g(M, a, b, x) &= M \exp(-\exp(a - bx)) \\
y_i | x_i, M_i, a_i, b_i &\sim Normal(g(M_i, a_i, b_i, x_i),\ f(x_i))
\end{aligned}
\tag{2.6}
$$

**Where**

$$
\begin{aligned}
P &= \text{number of random-effect groups} \\
|C_p| &= \text{Number of categories for group } p \\
N &= \text{Number of birds in the dataset} \\
C_p(i) &= \text{Category (under group } p\text{) of the } i\text{th sample} \\
x &= \text{Age, in days} \\
f(x) &= (2.2)
\end{aligned}
$$

Random-effect groups included sex (2 levels), year(x), island(x), island by year(x), hand-reared status(2), and ripe rimu status(2)

This part of the model, bird effect for a,b,M are fixed as 0.

Where hr is hand reared status, as provided by the Kākāpō Recovery Team, Year, Island, and Chick are self-explanatory and IslandBy is Island by Year, to capture interaction between the two effects.

The intercept explained ~68% of the data, mean absolute percentage error (MAPE) of 33%. The remaining cofactors (sex, hand-reared status, year, island, island by year) reduced MAPE to ~29% (explaining about 3% of the data). Chicks reduced MAPE to ~6.59% (the dataset left out used for validation was 6.45%). The means are taken for each of the three variables (M, a, b) for each bird to be used as phenotypes in downstream analyses.

#### Fit Individuals

**Priors**

for k in 1..157: Number of birds

$$
\begin{aligned}
M_k &\sim Normal(r, \sigma_r^2) \\
a_k &\sim Normal(r, \sigma_r^2) \\
b_k &\sim Normal(r, \sigma_r^2)
\end{aligned}
\tag{2.7}
$$

**Model**

for $i$ in 1..N : For each observation i

$$
\begin{aligned}
M_i &= M_{intercept} + \sum_{p=1}^{P} \beta_{M,C_p(i)} + M_k( \\
a_i &= a_{intercept} + \sum_{p=1}^{P} \beta_{a,C_p(i)} + a_k(i) \\
b_i &= b_{intercept} + \sum_{p=1}^{P} \beta_{b,C_p(i)} + b_k(i) \\
g(M_i, a_i, b_i, x_i) &= M_i \exp(-\exp(a_i - b_i x)) \\
y_i | x_i, M_i, a_i, b_i &\sim Normal(g(M_i, a_i, b_i, x_i),\ f(x_i)
\end{aligned}
\tag{2.8}
$$

**Where**

$N$ = Each observation of age (days), sex, year, island, weight, island by year, ha

There are a total of 8,308 observations used

$p_i$ = Bird for individual i

Growth rate GWAS results and predicted breeding values are found in Figures S24-44.

#### Fertile Eggs

Fertile egg phenotype data, as determined by the conservation management team, consisted of 122 observations from 48 mothers and 37 confirmed fathers, across 6 islands, across 15 years (ranging from 1981 - 2021) and was calculated as the ratio of reported fertile eggs to the total clutch size. Each mother had between 1 and 6 observations (mean; $\mu = 2.54$, standard deviation; $\sigma = 1.54$). Fathers had between 1 and 14 observations ($\mu = 3.30$, $\sigma = 2.88$). Mating pairs were unique in 91 observations, and 12 pairs had repeated pairings, with between 2 and 6 pairing events ($\mu = 2.58$, $\sigma = 1.24$). The fertile egg ratio across all observations had $\mu = 0.89$ and $\sigma = 0.22$.

We adjusted the fertile egg ratio for each breeding event for contributing factors to give a single adjusted phenotype value for each female for downstream analyses. To do so, we used TensorFlow probabilistic programming, which enables priors on the mean (termed the 'effect') and standard deviation (termed the 'scale') of the impact of each factor on the phenotype to be updated until the model converges. The model output is a set of distributions of the effects (and standard deviations thereof) for each of the factors on the phenotype.

In consultation with the Kākāpō Recovery Team and by exploring the impact of other possible factors on the raw phenotypes (i.e., via ANOVA), as well as experimenting with models to find the best fit, the final joint distribution model fitted was as follows:

**Priors**

$$\begin{aligned}
\mu &\sim Uniform(0.0,\ 3.0) \\
\sigma &\sim 1e-8 + Softplus(Uniform(0.01, 1.0)) \\
bird_effect &\sim Normal(\mu,\ \sigma) && \textbf{For 1...48} \\
father_effect &\sim Normal(\mu,\ \sigma)) && \textbf{For 1...37} \\
output_scale &\sim Normal(\mu,\ \sigma) \\
intercept &\sim Normal(\mu,\ \sigma)
\end{aligned}$$

**The Model**

$$\begin{aligned}
loc &= intercept + bird_effect * father_effect \\
scale &= sigmoid(output_scale) \\
y &\sim Normal(loc, scale)
\end{aligned}$$

**Where**

$$\begin{aligned}
\textbf{sigmoid}(x) &= 1/(1+\exp(-x)) \\
\textbf{softplus}(x) &= log(exp(x)+1)
\end{aligned}$$

Scale is the sigmoid of the smallest value of bird_scale or father_scale, added with 1e-4, to prevent the scale from reaching zero. This was determined from multiple experiments to identify a working model. The loss function is the target log probability. The final loss is ~53. Model results and GWAS Results are found in Figs S45-53.

#### Embryo Survival

Embryo survival phenotype is from the same dataset as Fertile Eggs. Embryo survival is defined as (1 - (dead embryo count / fertile eggs)) per clutch, and the ratio across all observations had μ = 0.81 and σ =0.28.

**Priors**

$$\mu \sim Uniform(-0.01,\ 0.01)$$
$$\sigma \sim Uniform(0.5, 1.0)$$
$$bird\_effect \sim Normal(\mu,\ \sigma) \qquad \textbf{For } 1\ldots48$$
$$zfather\_effect \sim Normal(\mu,\ \sigma)) \qquad \textbf{For } 1\ldots37$$
$$bird\_scale \sim Softplus(Normal(\mu,\ \sigma)) \qquad \textbf{For } 1\ldots48$$
$$father\_scale \sim Softplus(Normal(\mu,\ \sigma)) \qquad \textbf{For } 1\ldots37$$
$$intercept \sim Normal(\mu,\ \sigma)$$

**The Model**

$$loc = intercept + bird\_effect * father\_effect$$
$$scale = sigmoid(minimum(bird\_scale, father\_scale))$$
$$y \sim Normal(loc, scale)$$
$$\textbf{sigmoid}(x) = 1/(1 + \exp(-x))$$
$$\textbf{softplus}(x) = log(exp(x) + 1)$$

We adjusted the embryo survival ratio for each breeding event for contributing factors to give a single adjusted phenotype value for each female for downstream analyses. To do so, we used variational inference TensorFlow probabilistic programming, which enables priors on the mean (termed the ‘effect’) and standard deviation (termed the ‘scale’) of the impact of each factor on the phenotype to be updated until the model converges. The model output is a set of distributions of the effects (and standard deviations thereof) for each of the factors on the phenotype.

In consultation with the Kākāpō Recovery Team and by exploring the impact of other possible factors on the raw phenotypes (i.e., via ANOVA), as well as experimenting with models to find the best fit, the final joint distribution model fitted was as follows:

Model training was performed as Fertile Eggs, and the average loss of the final 1000 steps was 154.87. GWAS Results are found in Figures S54 - S60.

### Aspergillosis Susceptibility

The 2019 Aspergillosis outbreak–restricted to a single island (Whenua Hou)–was of urgent concern, with 25 infections and 10 deaths at the time of the dataset analysis. As many of the infected chicks did not have sequence information available, parents were used as a proxy for these individuals. For each infected individual with genetic data (n = xx), we defined aspergillosis susceptibility as 1. In the absence of genotype information for an infected individual (n = xx), we assigned its two genotyped parents an aspergillosis susceptibility, defined as the ratio of the total number of their infected offspring to the total clutch size. Aspergillosis offspring ratios ranged from 0.08 to 0.75 for uninfected parents. Given that many individuals were not exposed to aspergillosis, uninfected individuals were assigned a missing phenotype value (rather than defined with susceptibility = 0) for the GWAS and breeding value calculations. Model and GWAS Results are found in Figures S61 - S68.

### Phenotype Results

#### Egg Shape Index

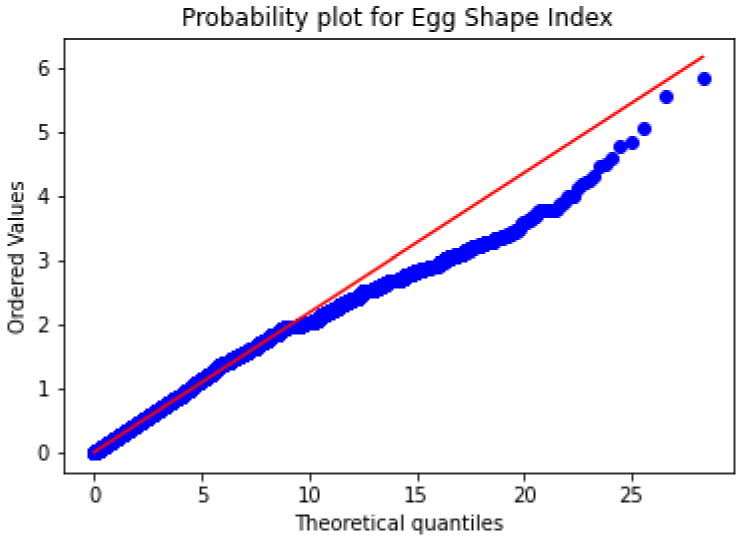


Fig S8. QQ-Plot for Egg Shape Index marker p-values.

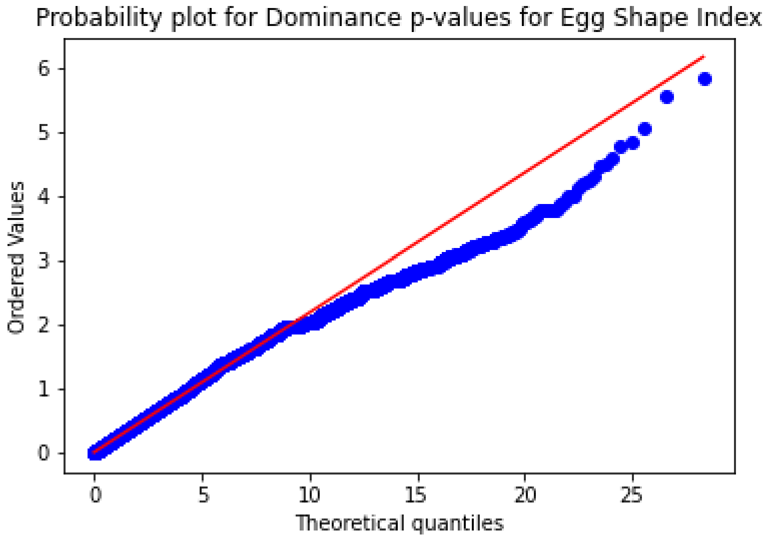


Fig S9. QQ-Plot for Egg Shape Index marker p-values, Dominance model.

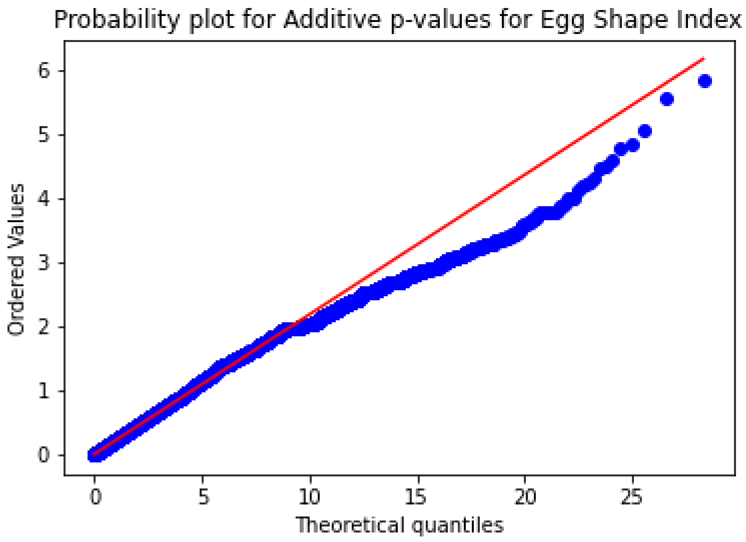


Fig S10. QQ-Plot for Egg Shape Index marker p-values, Additive model..

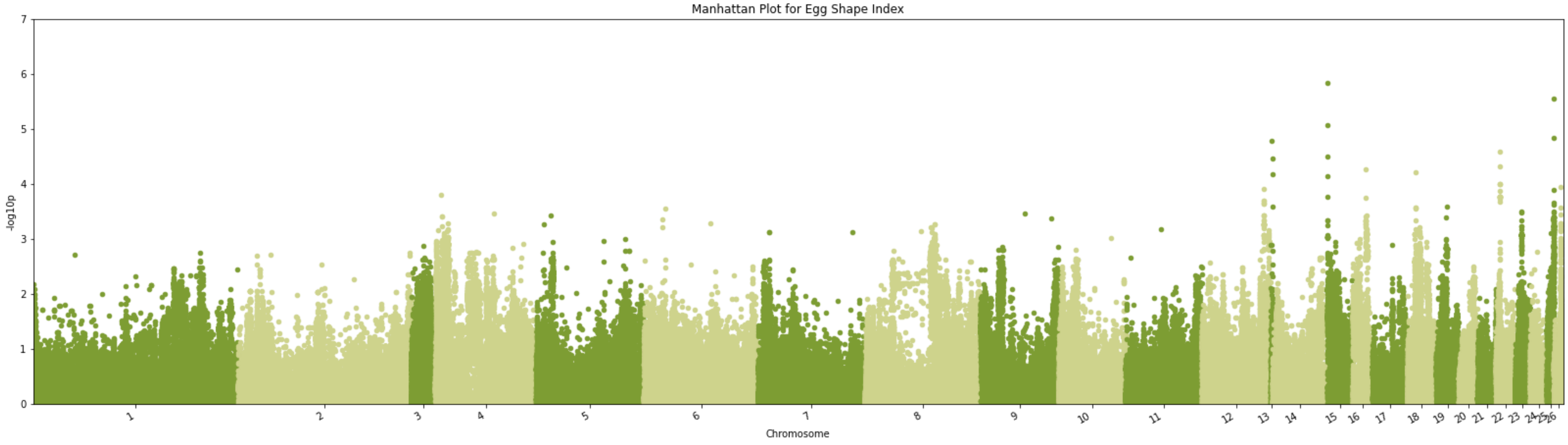


Fig S11. Manhattan plot of -log10(p) using the marker p-value from TASSEL GWAS for Egg Shape Index for chromosomes 1 through 26 (Supplemental Table S2 for Chromosome mapping).

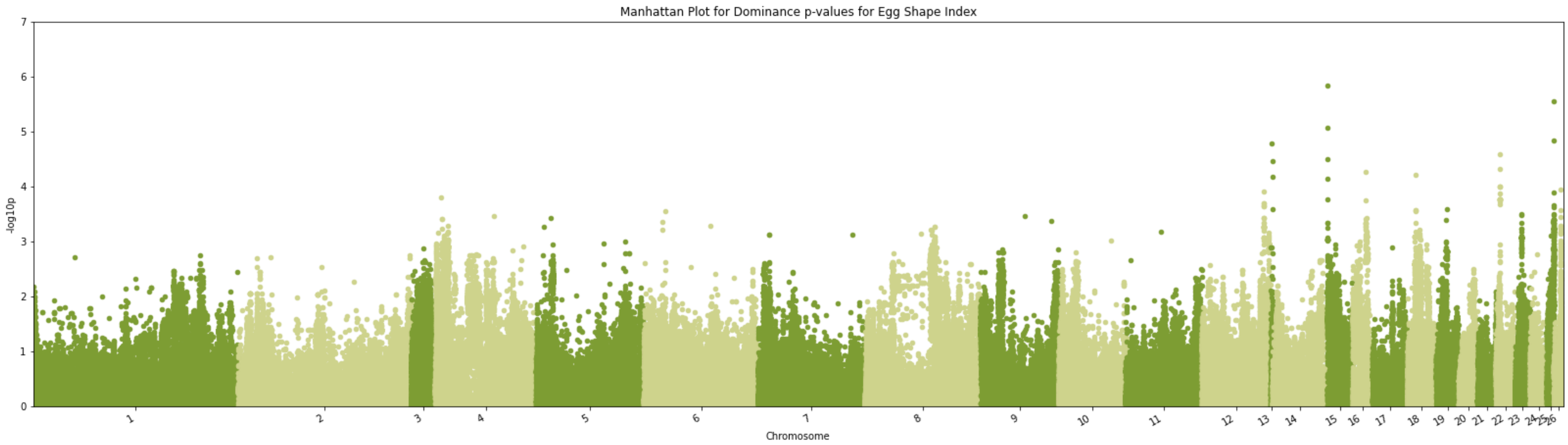


Fig S12. Manhattan plot -log10(p) using the marker p-value from TASSEL GWAS for Egg Shape Index for chromosomes 1 through 26 (Supplemental Table S2 for Chromosome mapping), Dominance model.

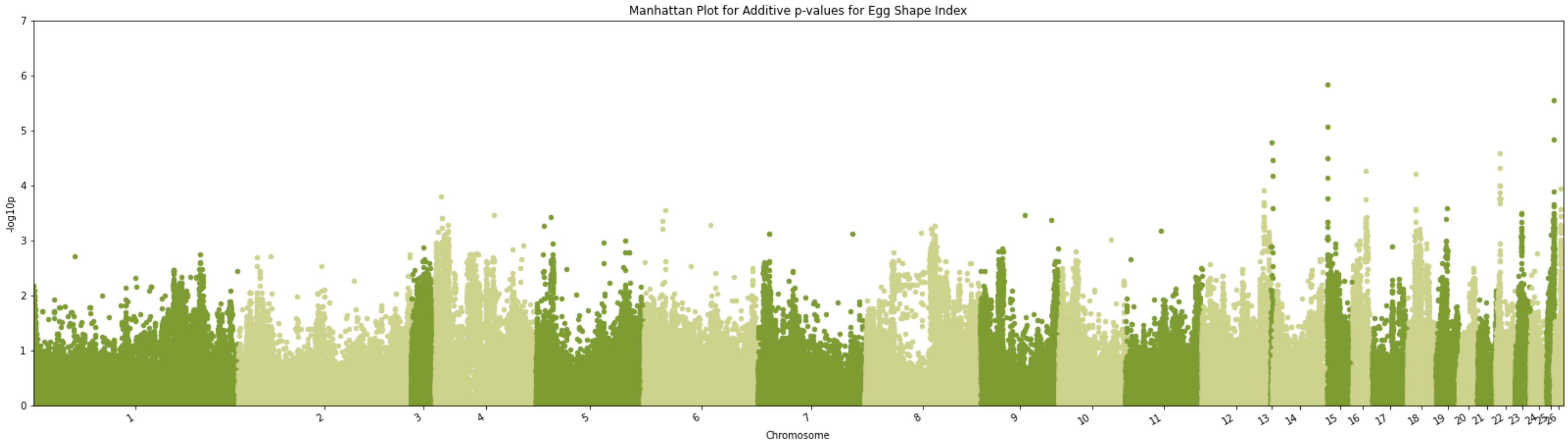


Fig S13. Manhattan plot -log10(p) using the marker p-value from TASSEL GWAS for Egg Shape Index for chromosomes 1 through 26 (Supplemental Table S2 for Chromosome mapping), Additive.

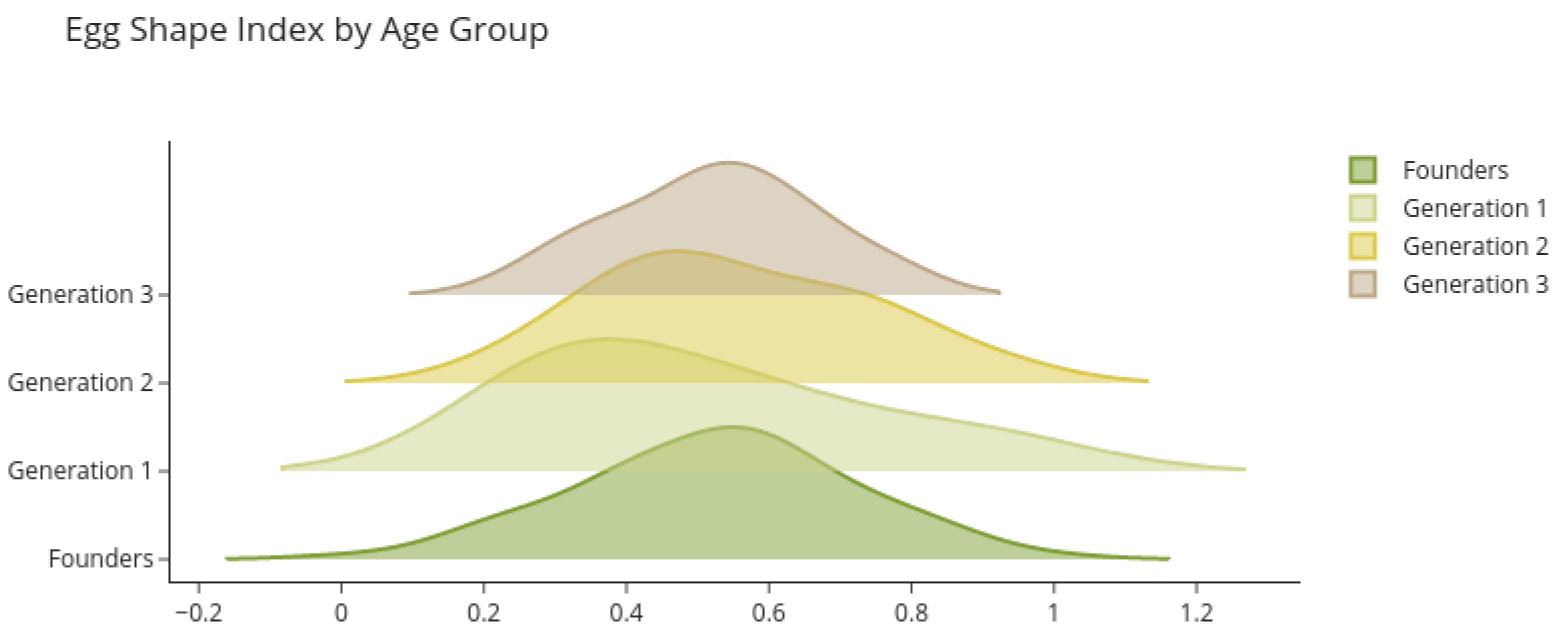


Fig S14. Predicted breeding values by age group (Table 1) for Egg Shape Index.

#### Clutch Size Figures

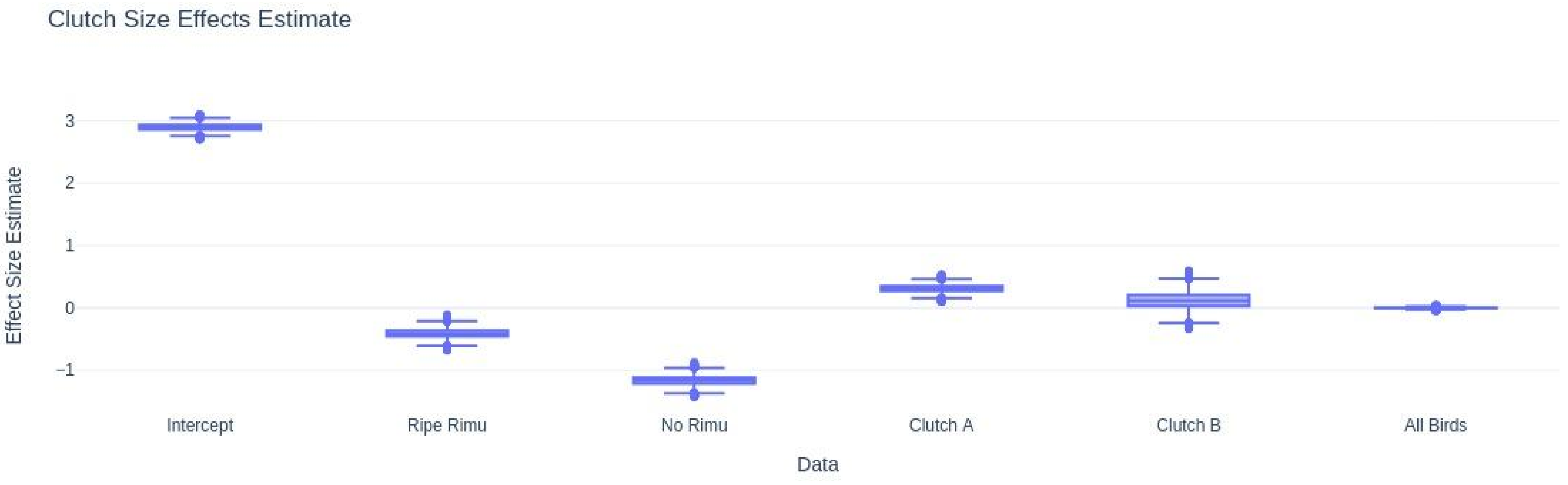


Fig S15. Clutch size effect estimates for the model.

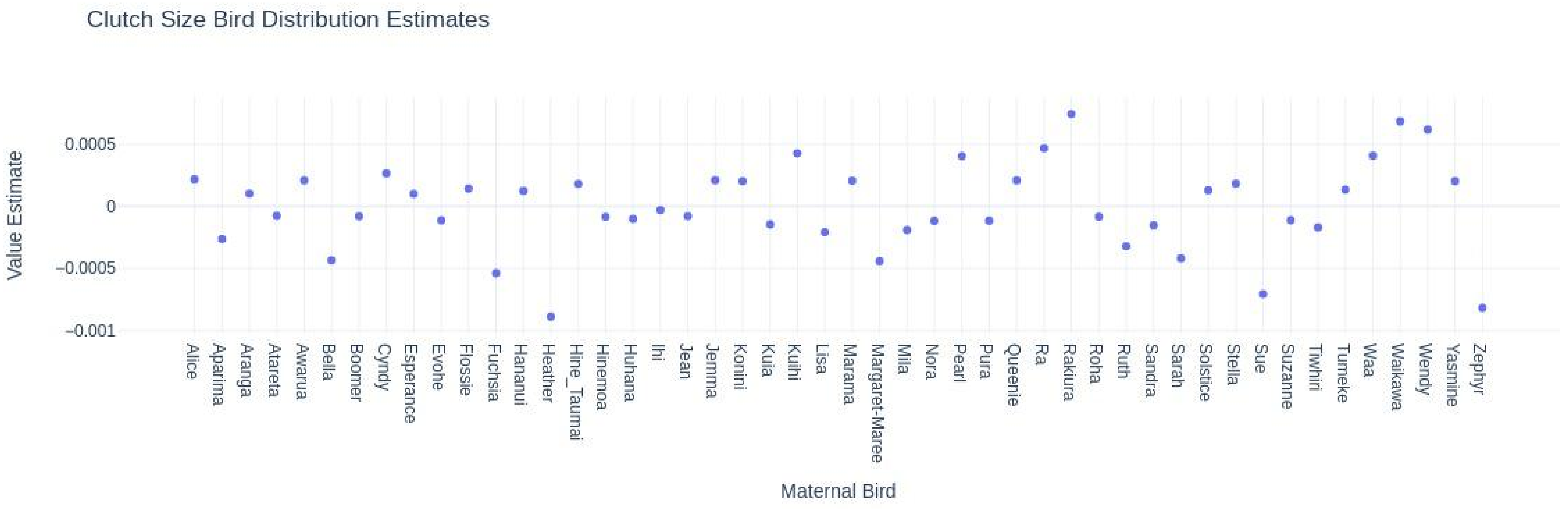


Fig S16. Clutch size effect estimates for mother birds.

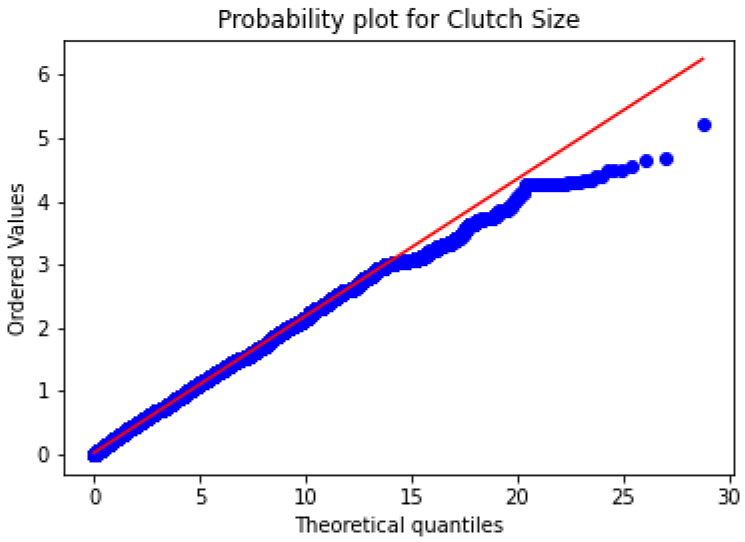

Fig S17. Clutch size QQ-plot for marker p-values.

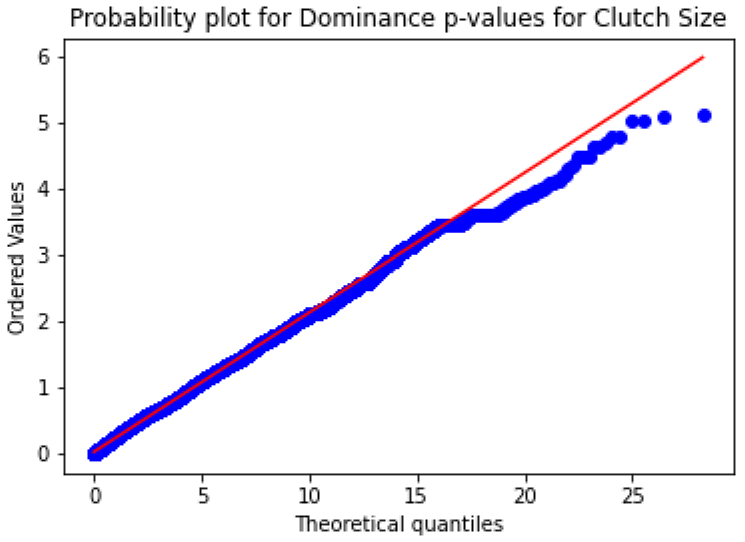

Fig S18. Clutch size QQ-plot for dominance p-values.

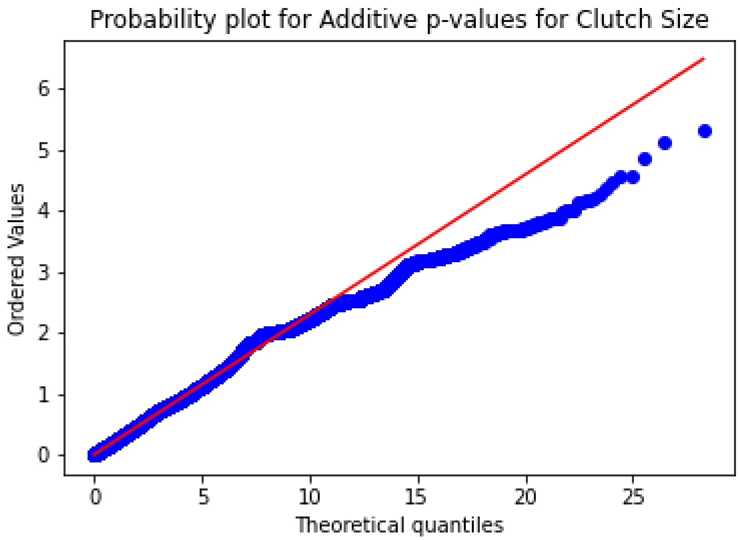


Fig S19. Clutch size QQ-plot for additive p-values.

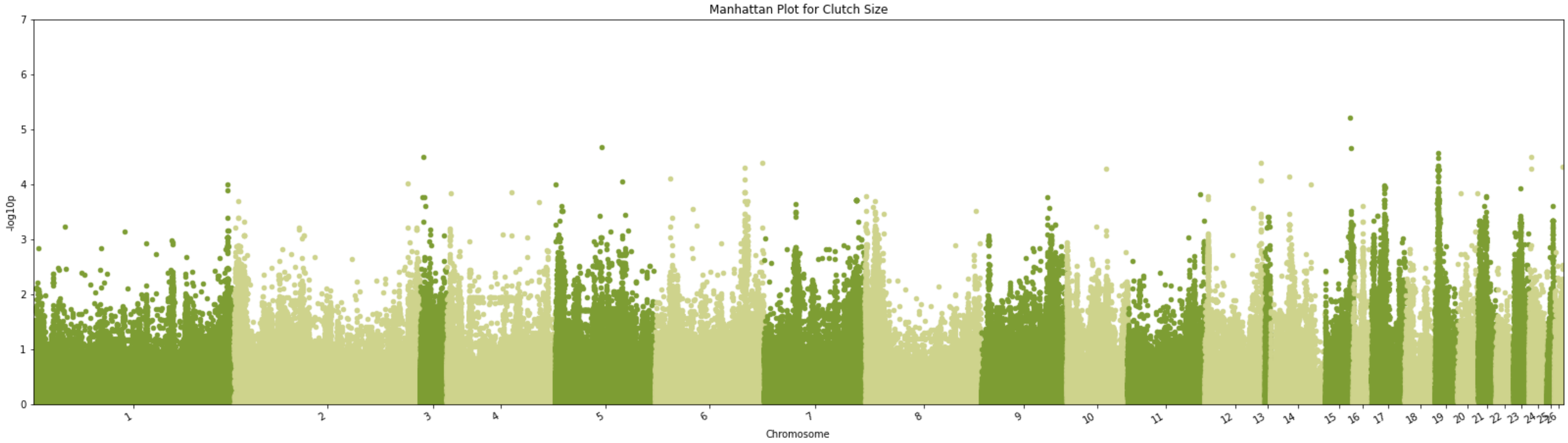


Fig S20. Manhattan plot -log10(p) using the marker p-value from TASSEL GWAS for Clutch Size for chromosomes 1 through 26 (Supplemental Table S2 for Chromosome mapping).

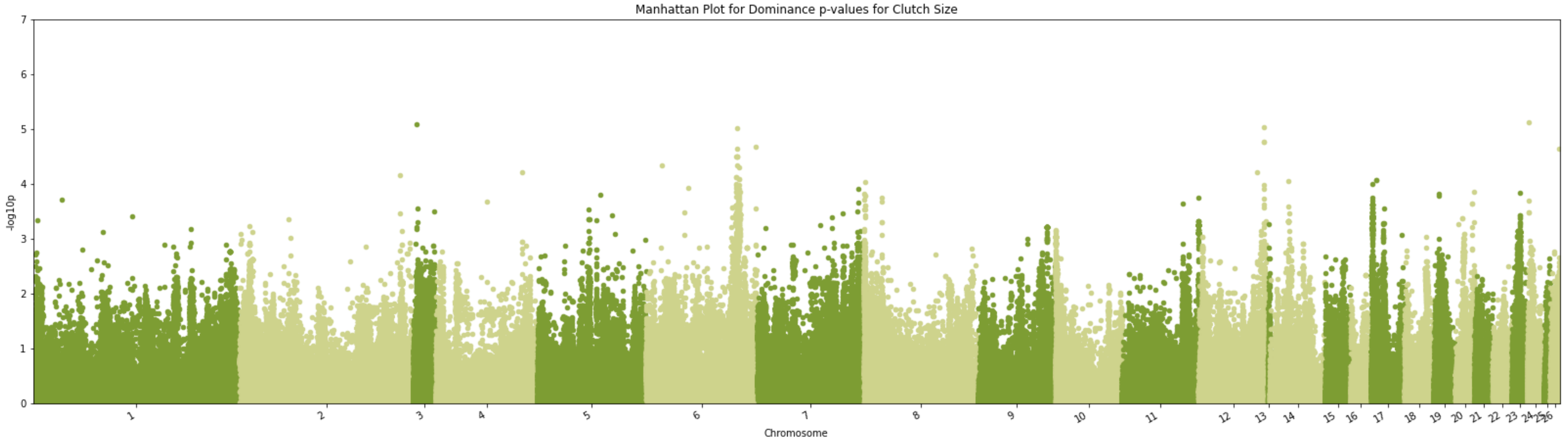


Fig S21. Manhattan plot -log10(p) using the marker p-value from TASSEL GWAS for Clutch Size for chromosomes 1 through 26 (Supplemental Table S2 for Chromosome mapping). Dominance model.

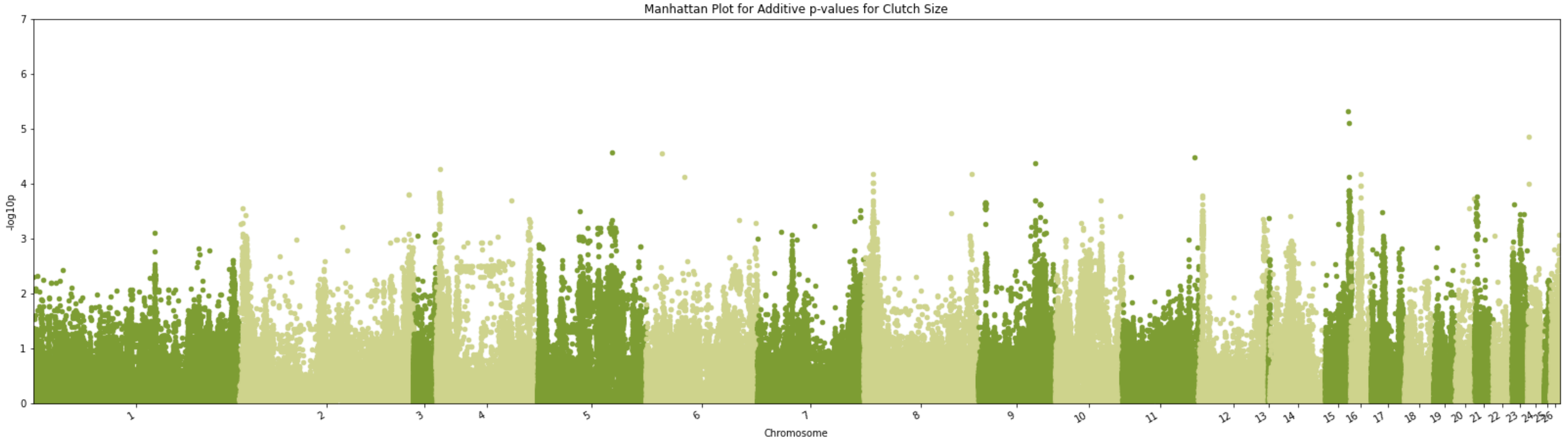


Fig S22. Manhattan plot -log10(p) using the marker p-value from TASSEL GWAS for Clutch Size for chromosomes 1 through 26 (Supplemental Table S2 for Chromosome mapping). Additive model.

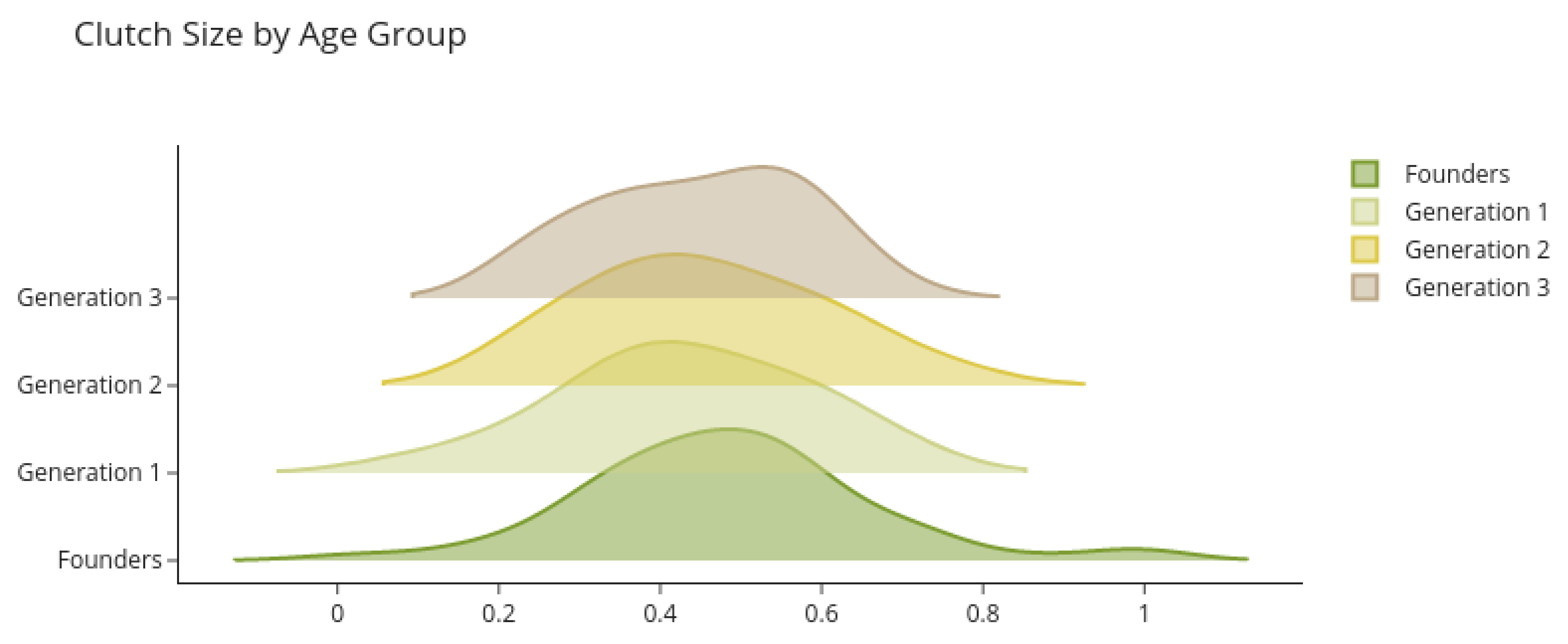


Fig S23. Predicted breeding values by age group for Clutch Size.

Growth Rate - M Parameter

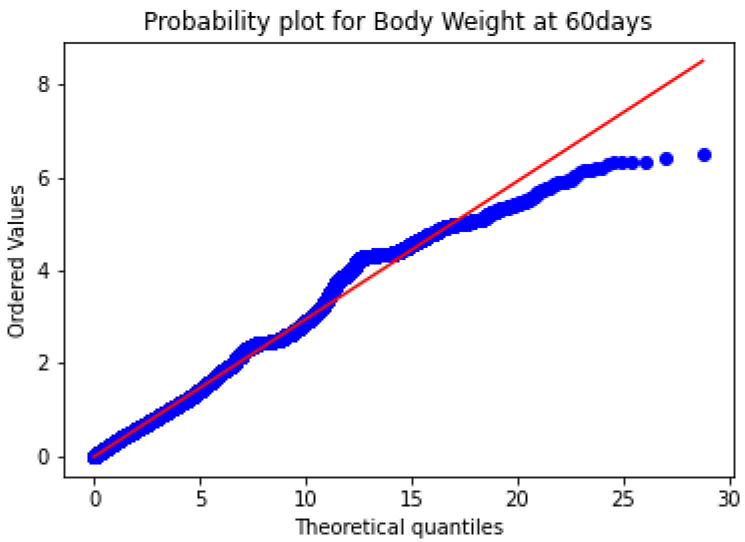


Fig S24. Body Weight at 60days QQ-plot.

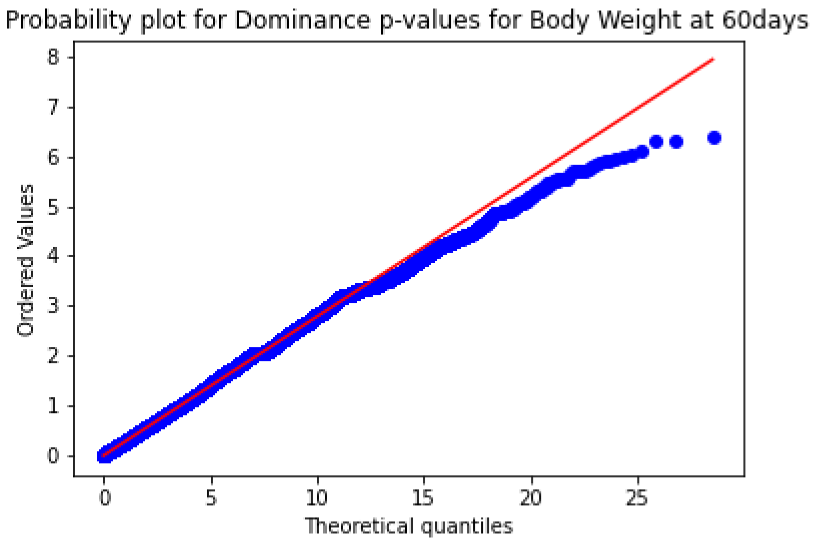


Fig S25. Body Weight at 60days QQ-plot. Dominance model.

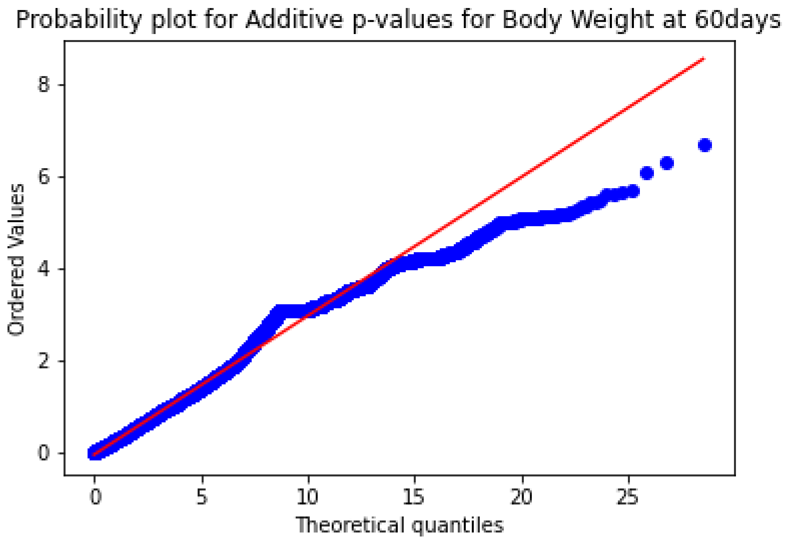

Fig S26. Body Weight at 60days QQ-plot. Dominance model.

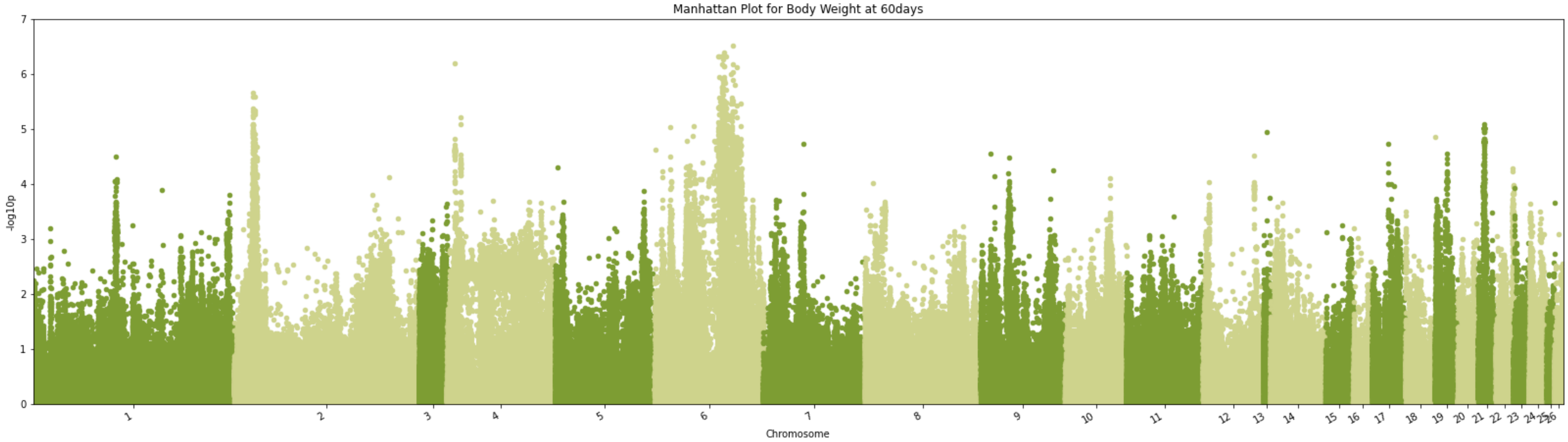

Fig S27. Manhattan plot -log10(p) using the marker p-value from TASSEL GWAS for Body Weight at 60 days for chromosomes 1 through 26 (Supplemental Table S2 for Chromosome mapping).

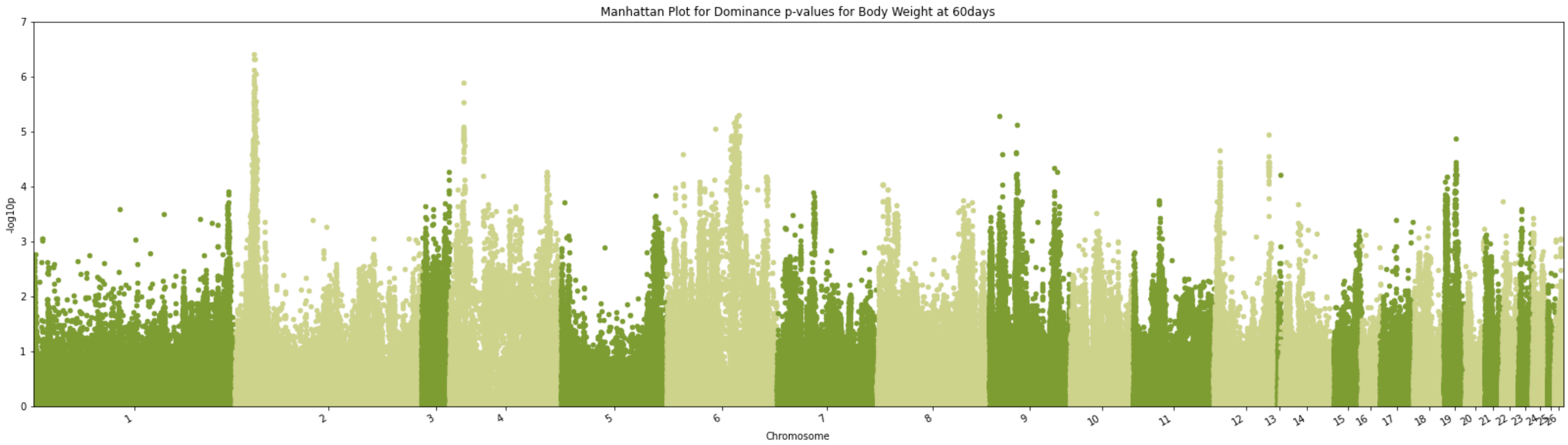


Fig S28. Manhattan plot -log10(p) using the marker p-value from TASSEL GWAS for Body Weight at 60 days for chromosomes 1 through 26 (Supplemental Table S2 for Chromosome mapping). Dominance model.

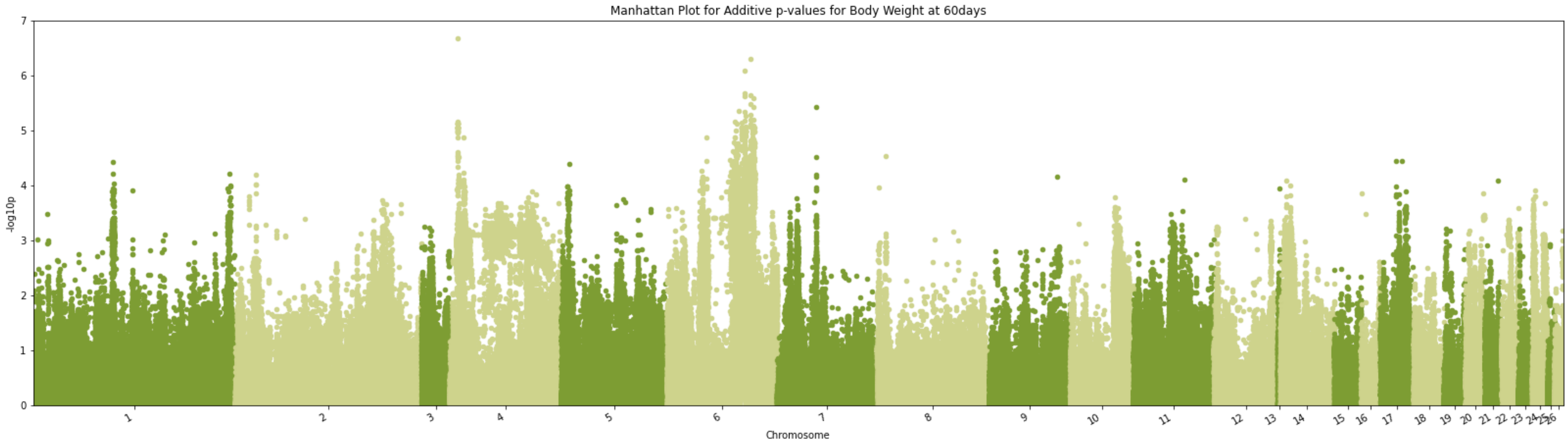


Fig S29. Manhattan plot -log10(p) using the marker p-value from TASSEL GWAS for Body Weight at 60 days for chromosomes 1 through 26 (Supplemental Table S2 for Chromosome mapping). Additive model.

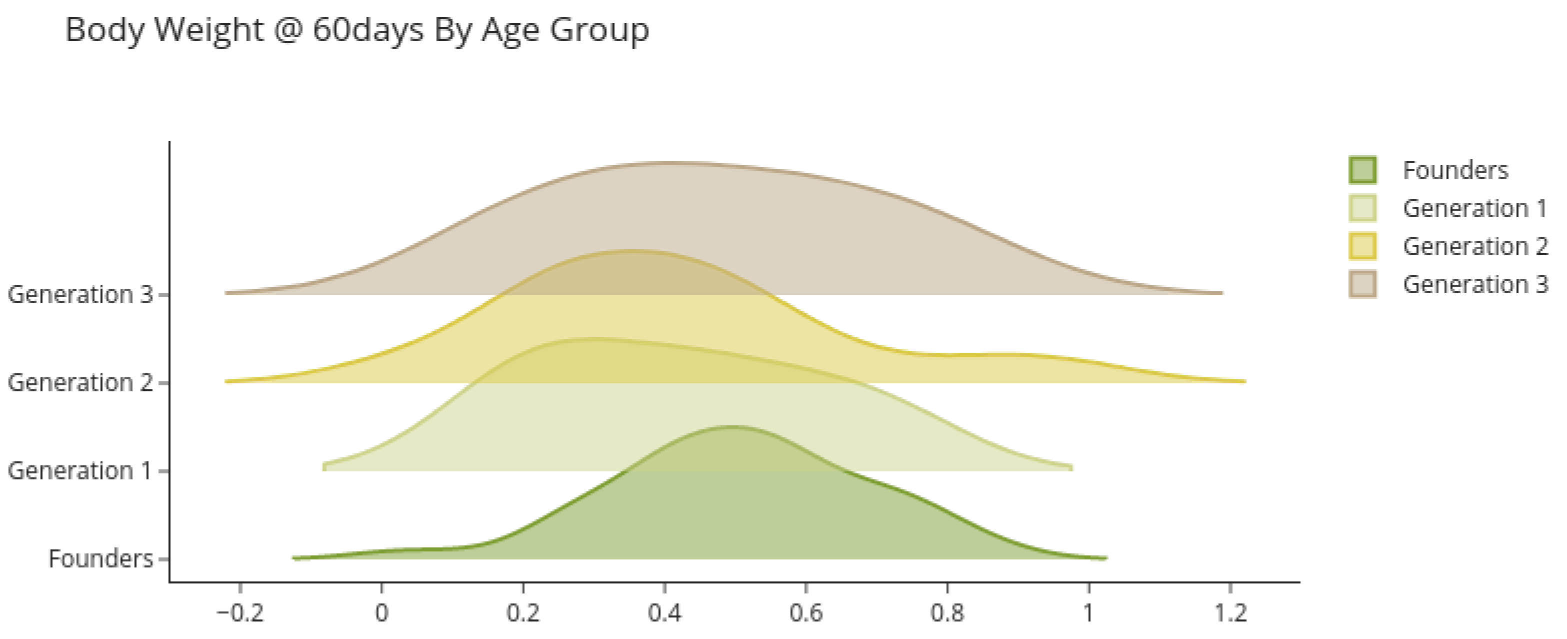


Fig S30. Predicted breeding values by age group for Body Weight at 60 days.

#### Growth Rate - a Parameter - Growth Curve Coefficient

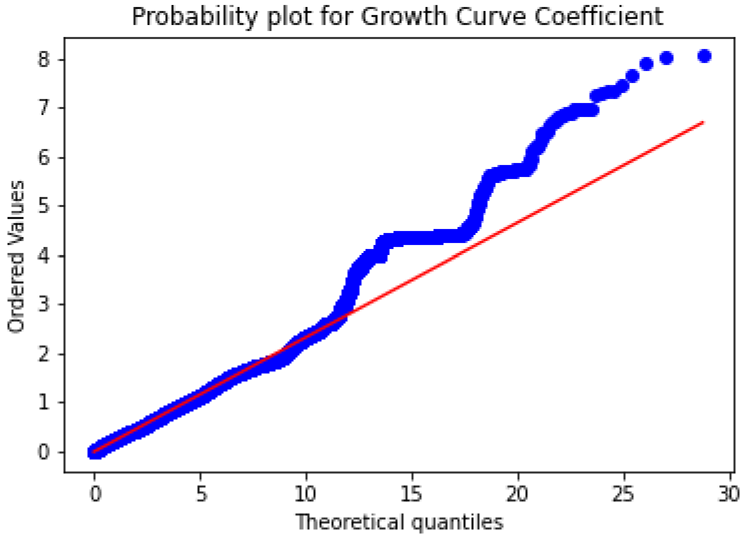


Fig S31. Growth Curve Coefficient (Gompertz Curve) QQ-Plot.

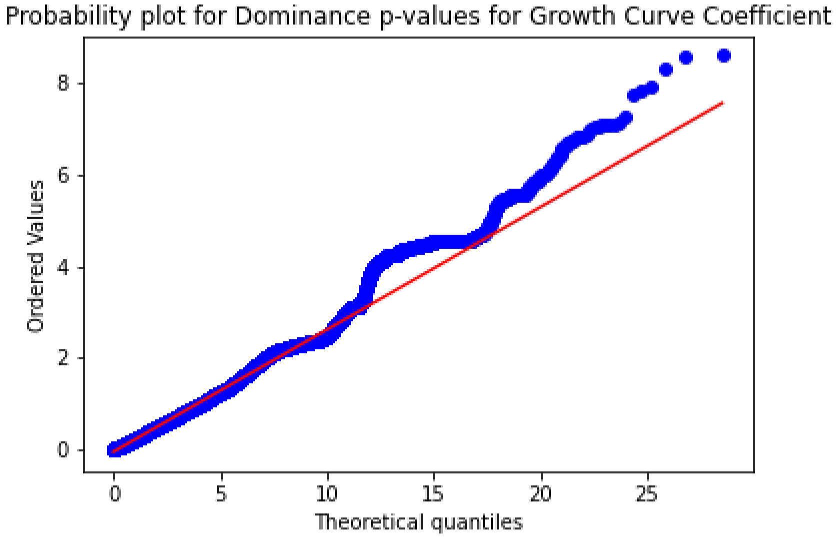


Fig S32. Growth Curve Coefficient (Gompertz Curve) QQ-Plot. Dominance model.

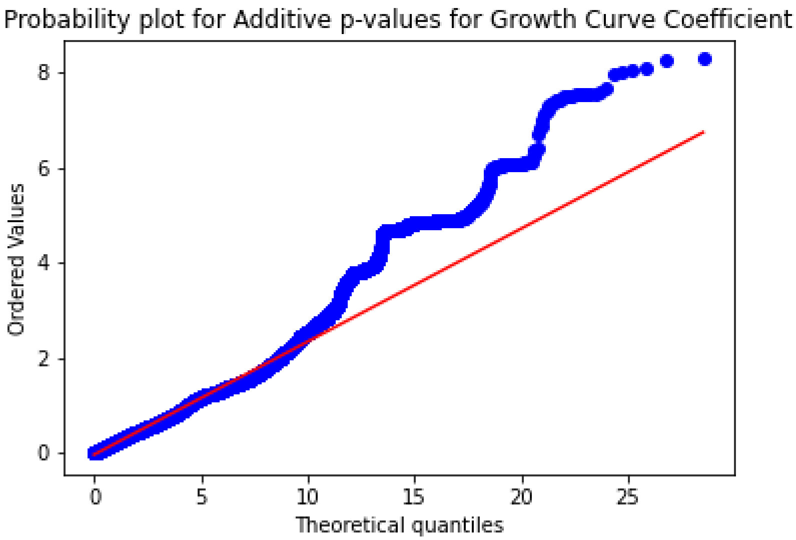

Fig S33. Growth Curve Coefficient (Gompertz Curve) QQ-Plot. Additive model.

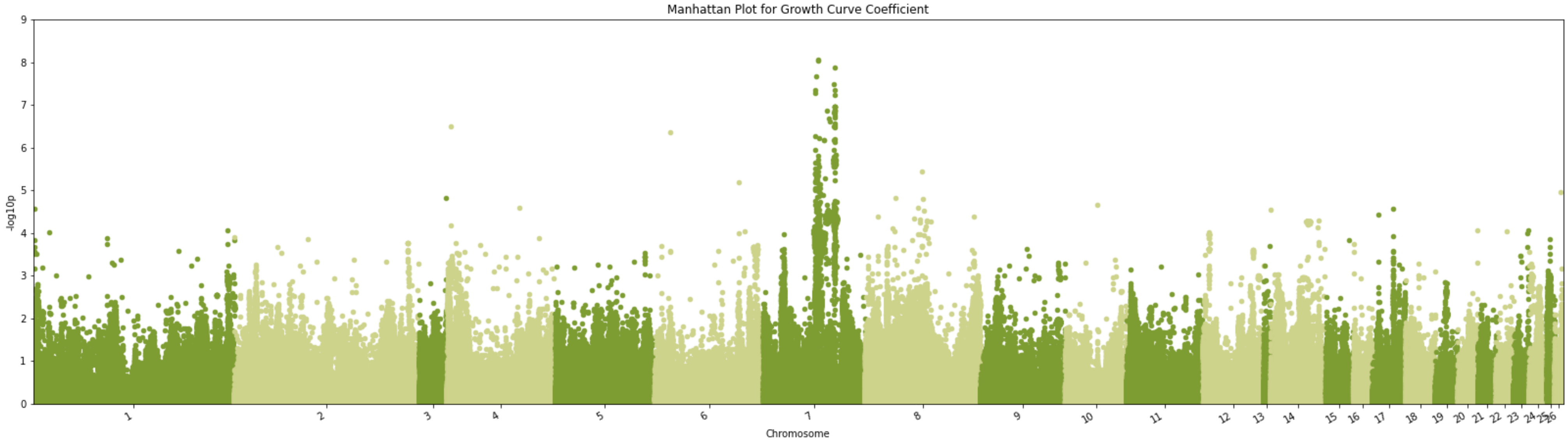

Fig S34. Manhattan plot -log10(p) using the marker p-value from TASSEL GWAS for Growth Curve Coefficient for chromosomes 1 through 26 (Supplemental Table S2 for Chromosome mapping).

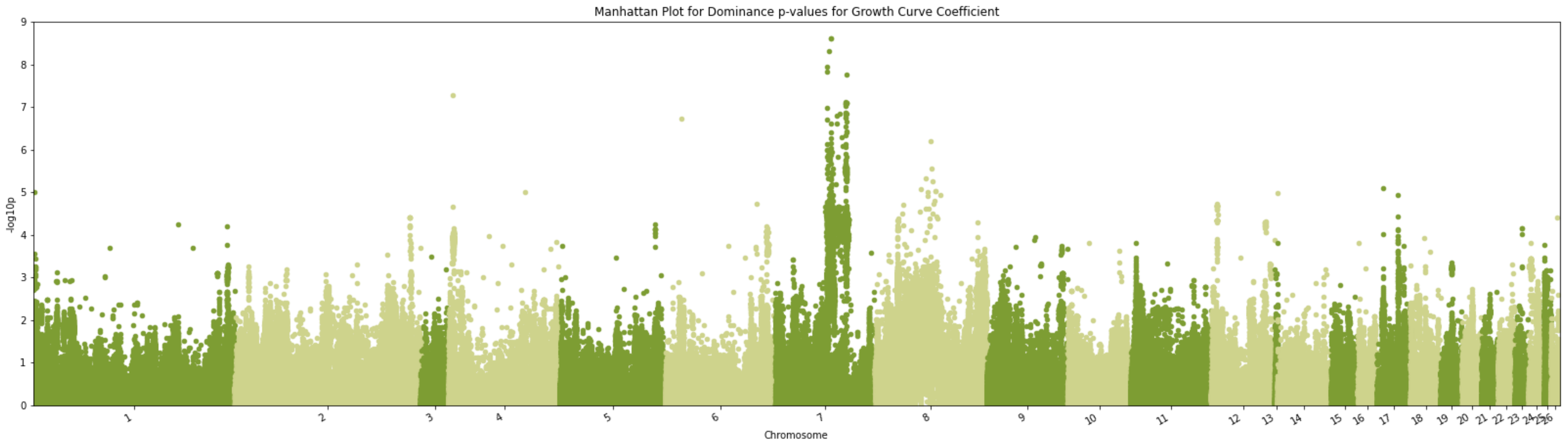


Fig S35. Manhattan plot -log10(p) using the marker p-value from TASSEL GWAS for Growth Curve Coefficient for chromosomes 1 through 26 (Supplemental Table S2 for Chromosome mapping). Dominance model.

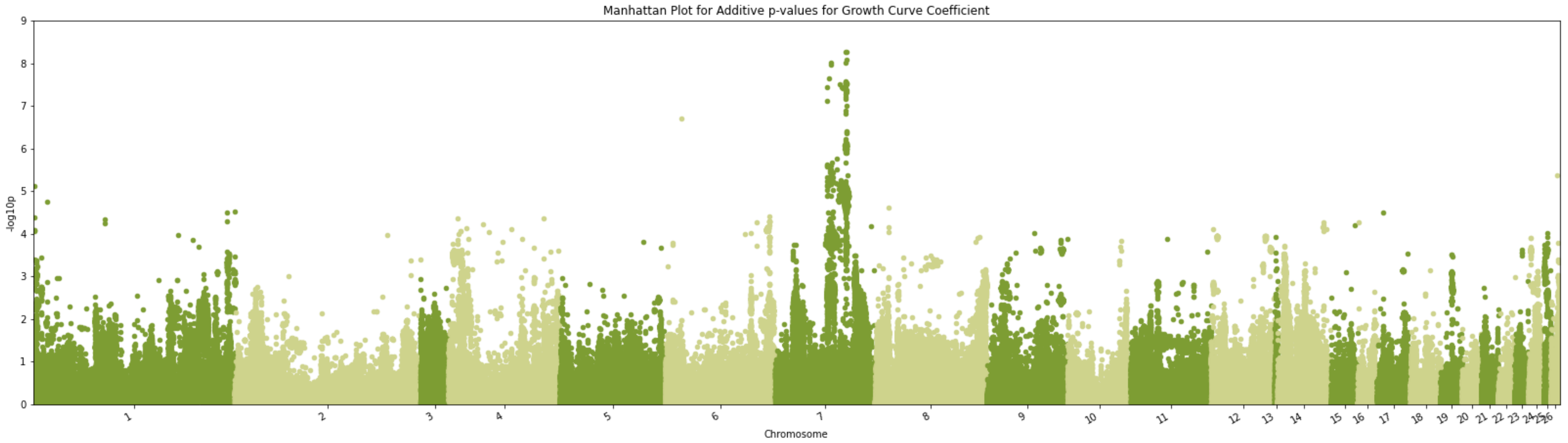


Fig S36. Manhattan plot -log10(p) using the marker p-value from TASSEL GWAS for Growth Curve Coefficient for chromosomes 1 through 26 (Supplemental Table S2 for Chromosome mapping). Additive model.

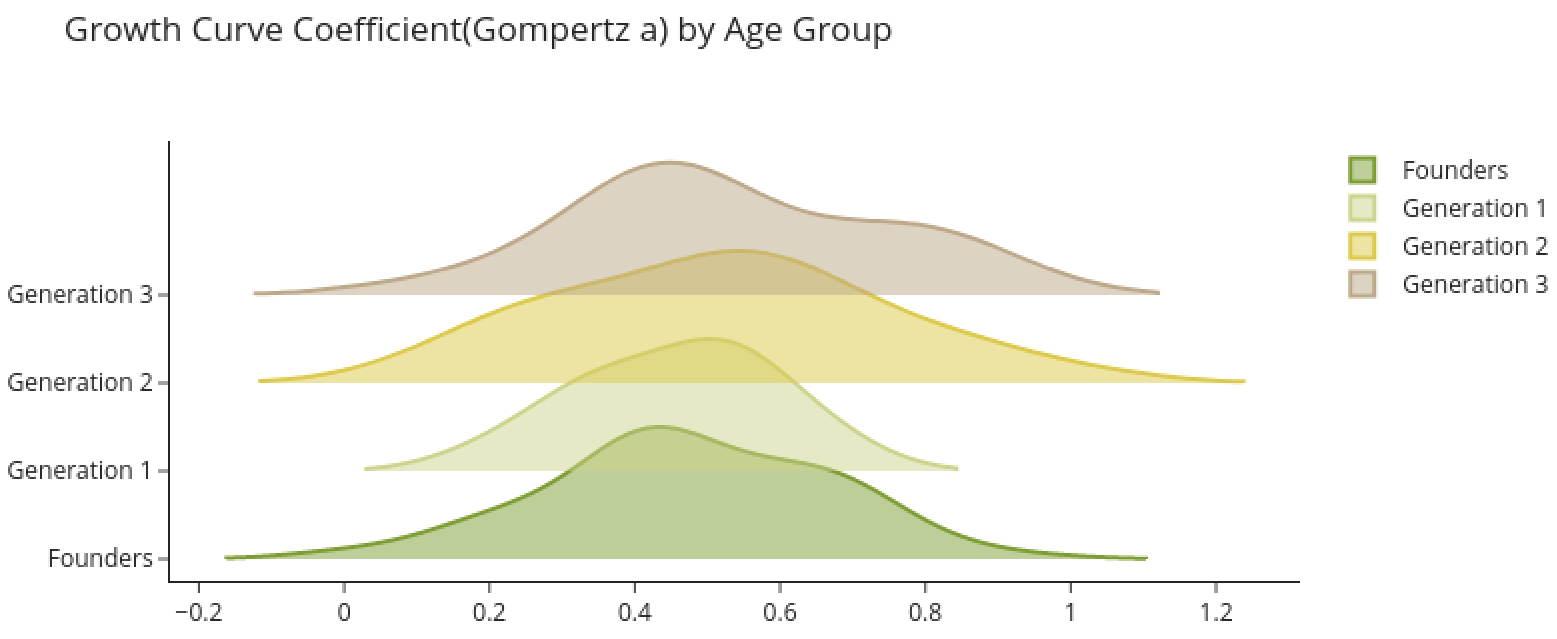


Fig S37. Predicted breeding values by age-group for Gompertz Growth Curve Coefficient.

Growth Rate - b Parameter - Growth Rate

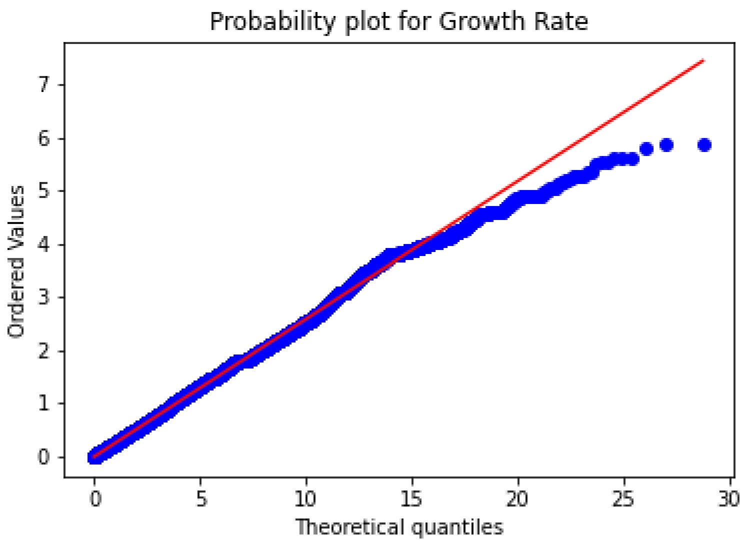


Fig S38. Growth Curve Growth Rate (Gompertz Curve) QQ-Plot.

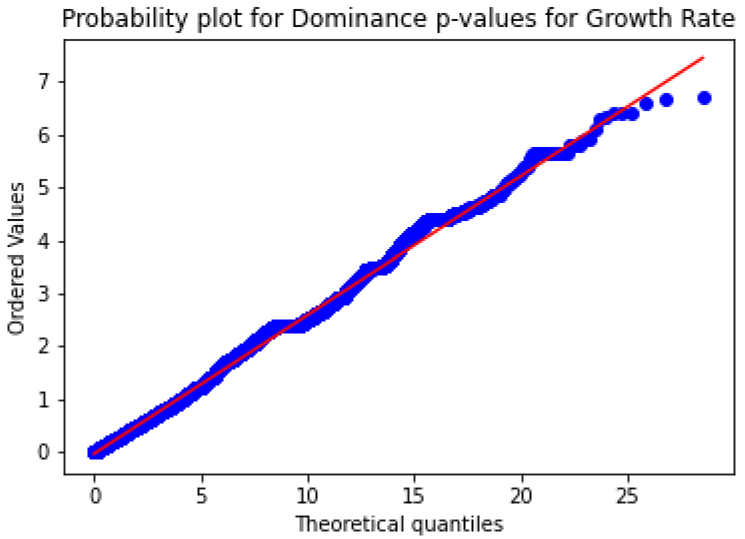


Fig S39. Growth Curve Growth Rate (Gompertz Curve) QQ-Plot. Dominance model.

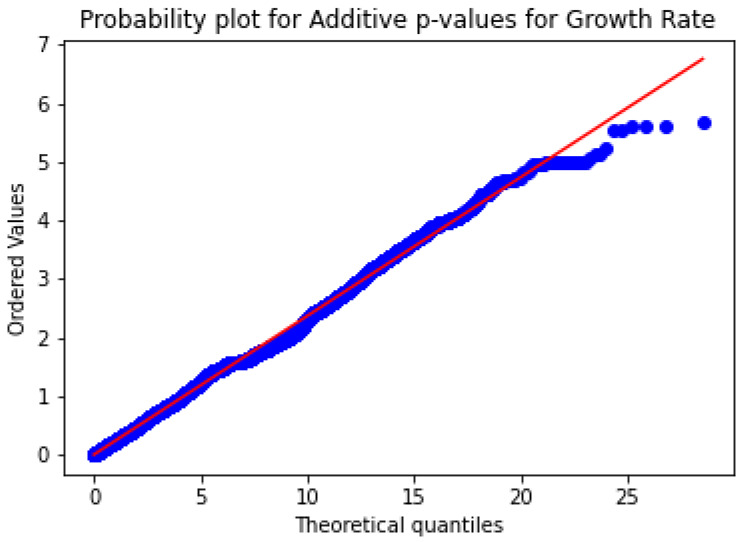


Fig S40. Growth Curve Growth Rate (Gompertz Curve) QQ-Plot. Additive model.

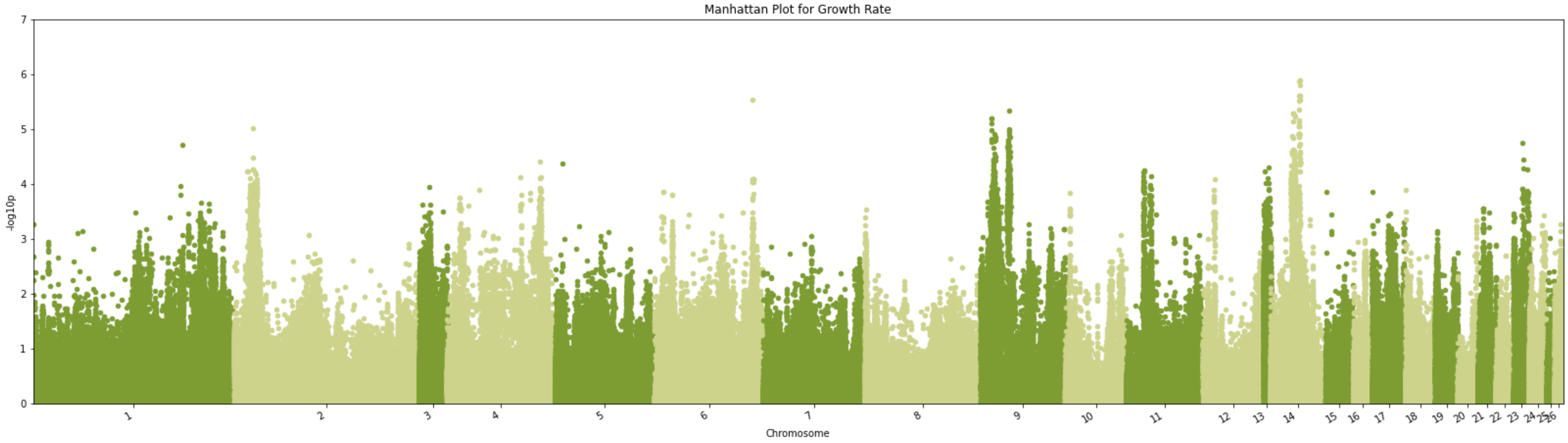


Fig S41. Manhattan plot -log10(p) using the marker p-value from TASSEL GWAS for Growth Curve Growth Rate for chromosomes 1 through 26 (Supplemental Table S2 for Chromosome mapping).

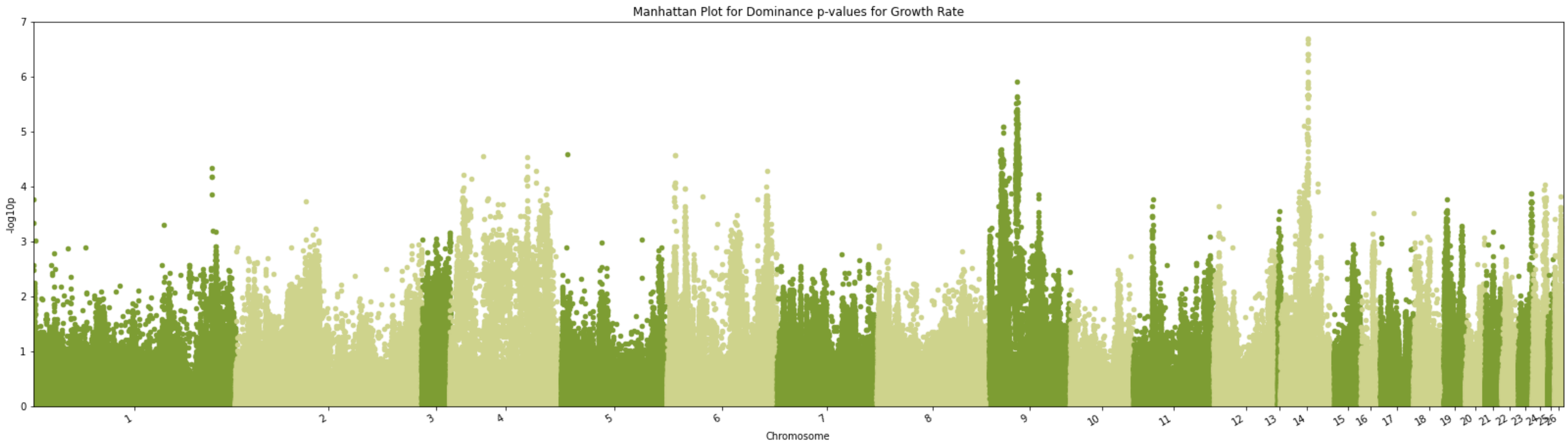


Fig S42. Manhattan plot -log10(p) using the marker p-value from TASSEL GWAS for Growth Curve Growth Rate for chromosomes 1 through 26 (Supplemental Table S2 for Chromosome mapping). Dominance model.

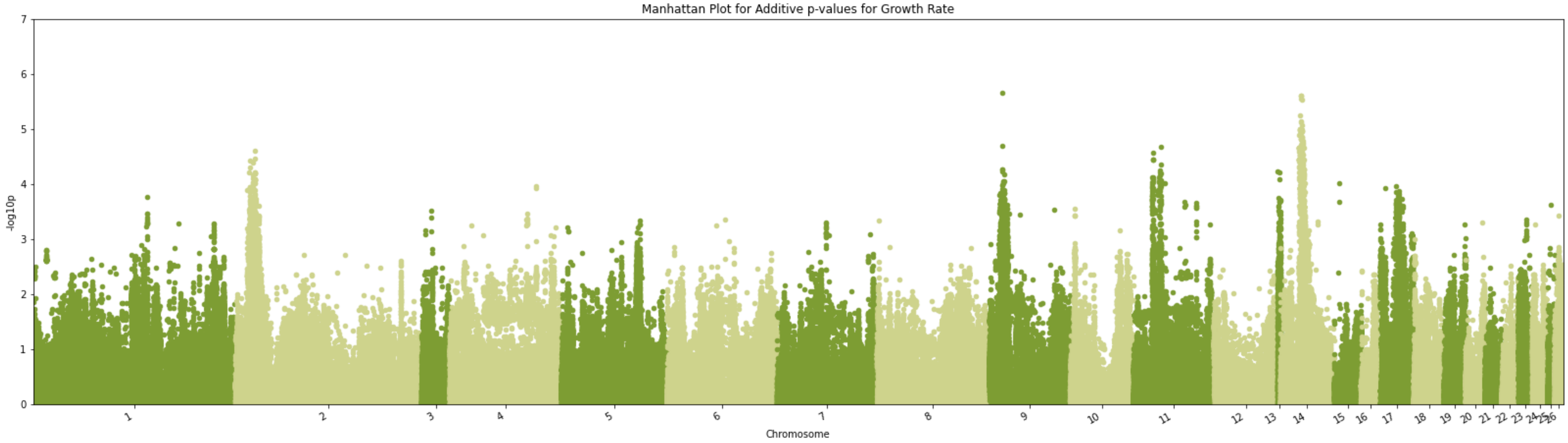


Fig S43. Manhattan plot -log10(p) using the marker p-value from TASSEL GWAS for Growth Curve Growth Rate for chromosomes 1 through 26 (Supplemental Table S2 for Chromosome mapping). Additive model.

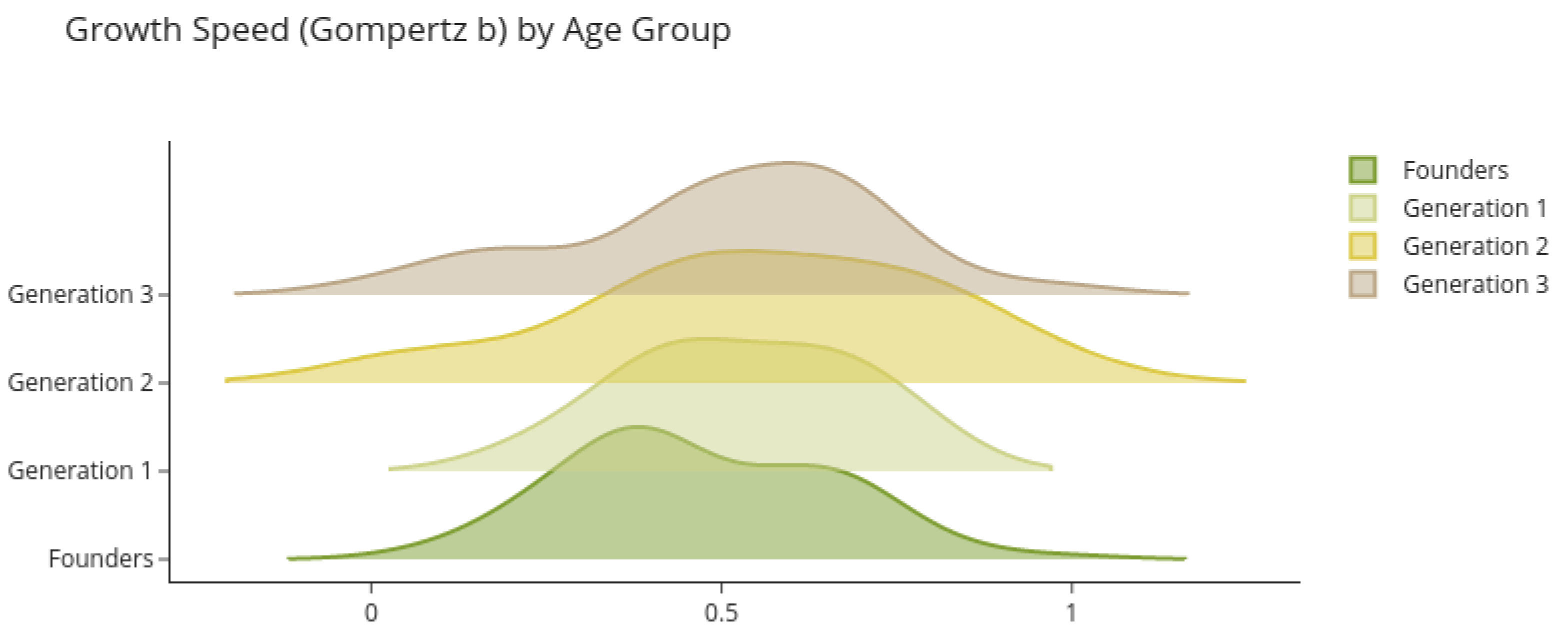


Fig S44. Predicted Breeding Values for Growth Speed of Gompertz Curve, by generation.

### Fertile Eggs

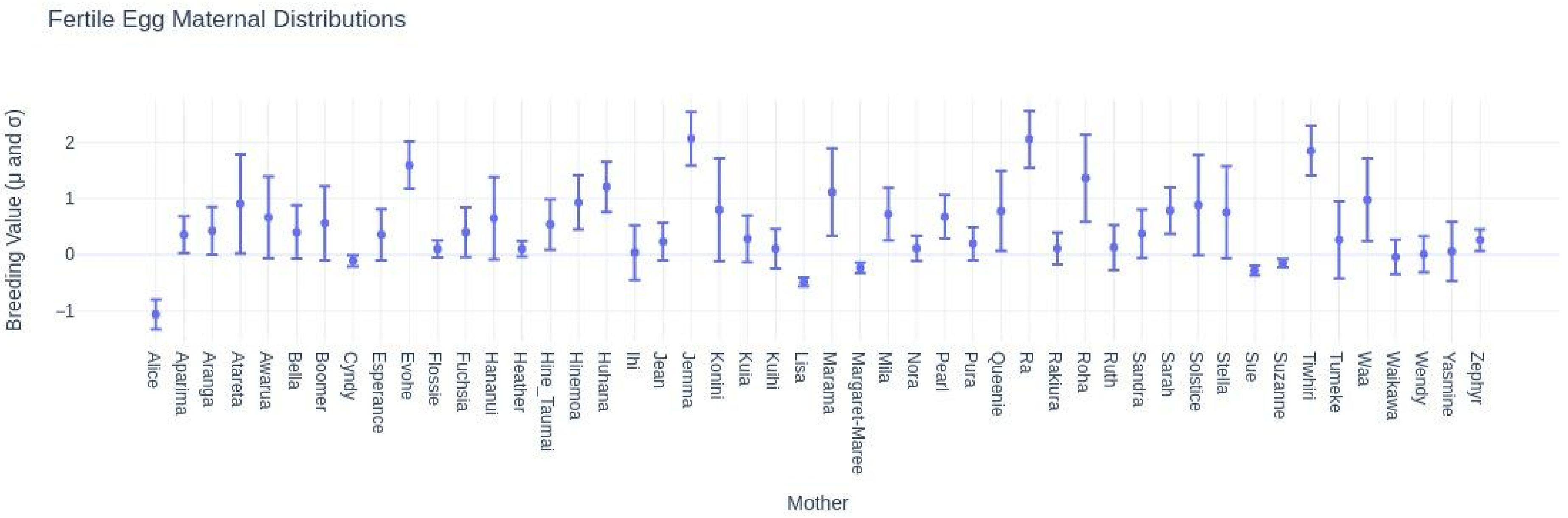


Fig S45. Distribution of maternal effect sizes in probabilistic model.

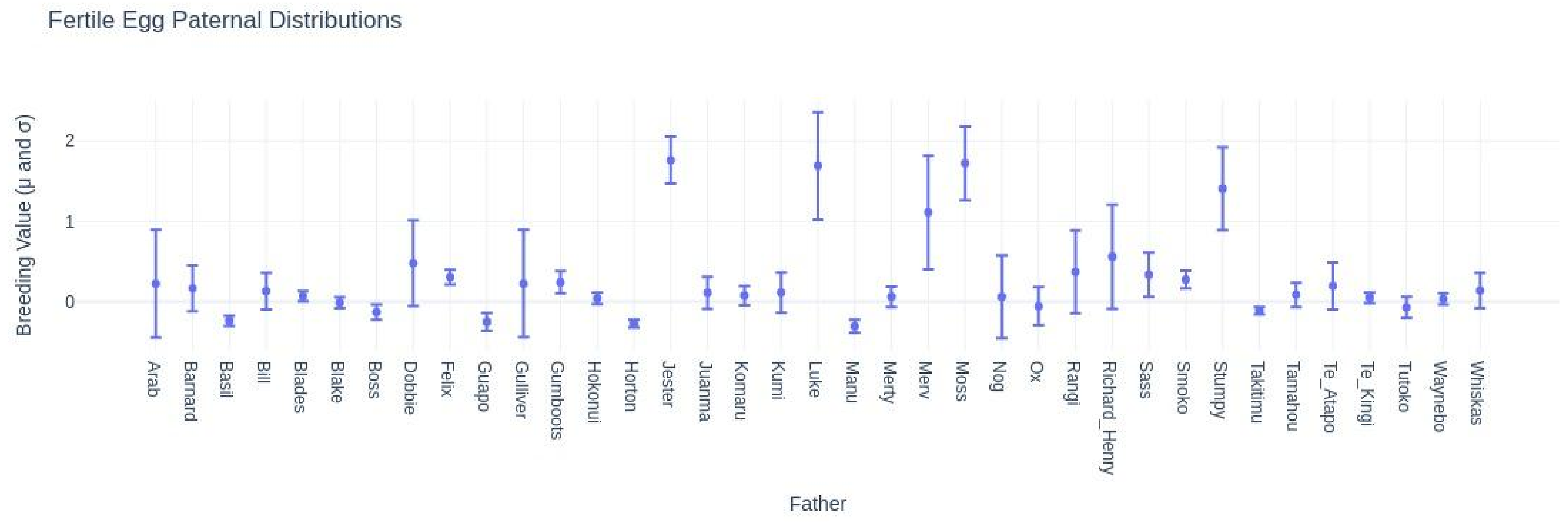


Fig S46. Distribution of paternal effect sizes in probabilistic model.

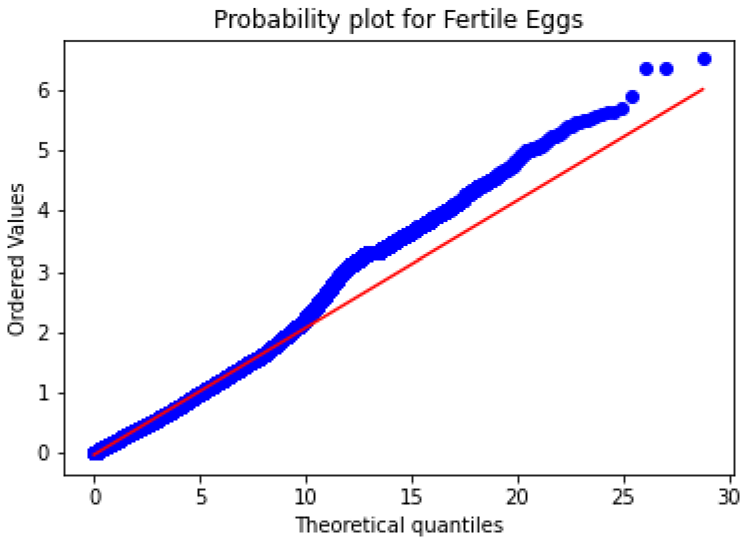


Fig S47. QQ-Plot for Fertile Eggs GWAS.

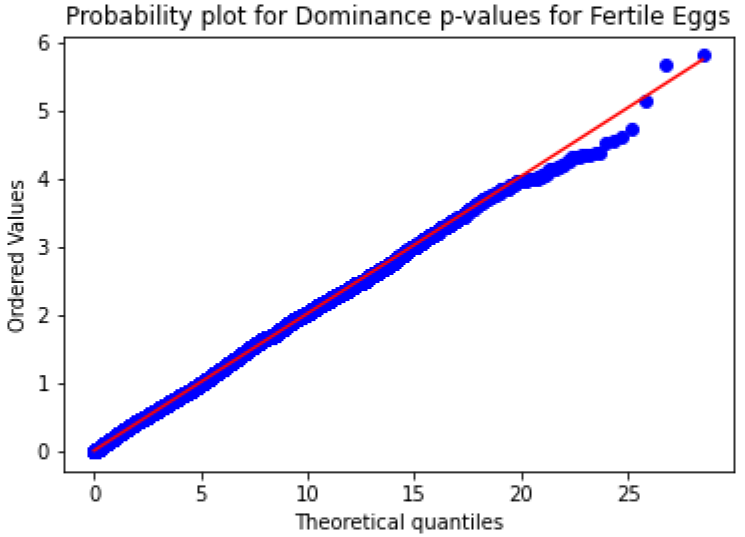


S48. QQ-Plot for Fertile Eggs GWAS. Dominance model.

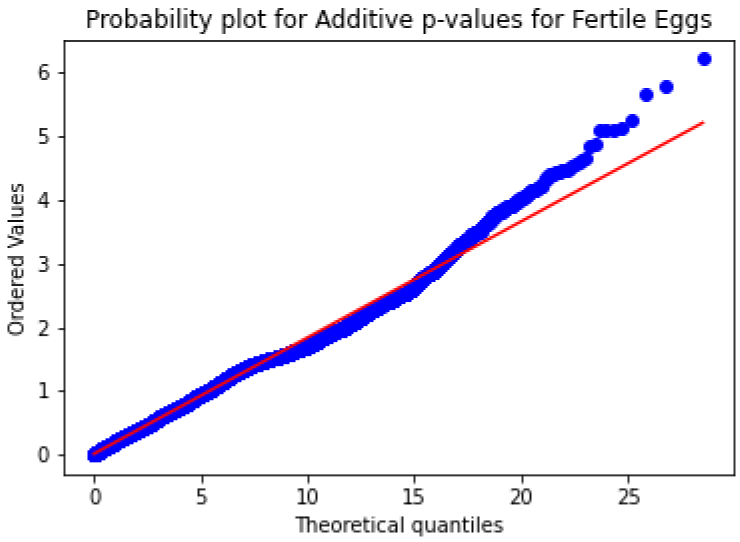


Fig S49. QQ-Plot for Fertile Eggs GWAS. Additive model.

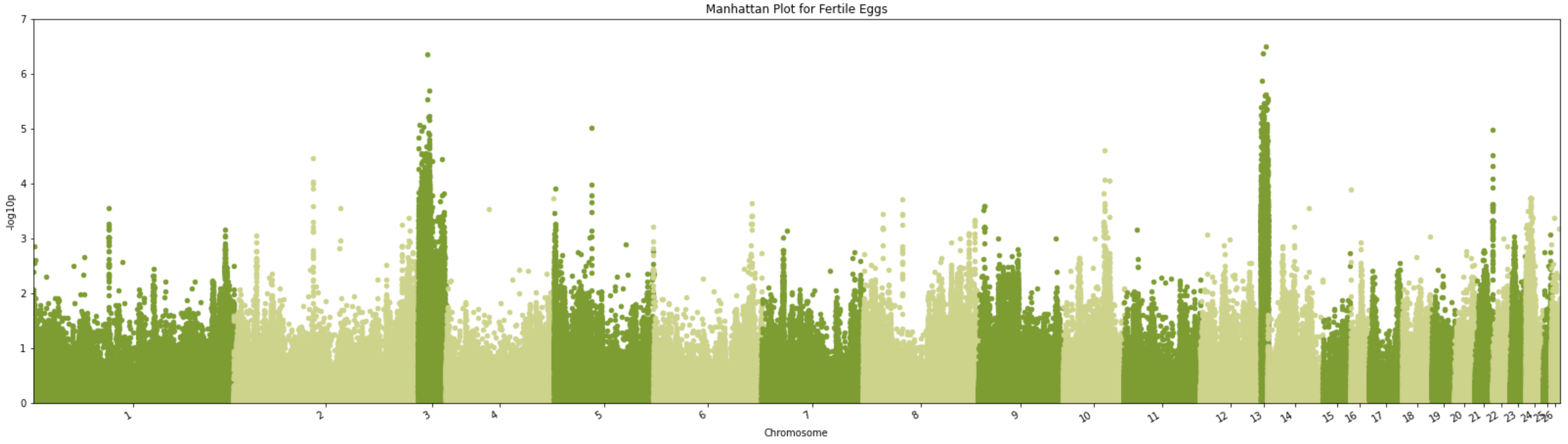


Fig S50. Manhattan plot -log10(p) using the marker p-value from TASSEL GWAS for Fertile Eggs for chromosomes 1 through 26 (Supplemental Table S2 for Chromosome mapping).

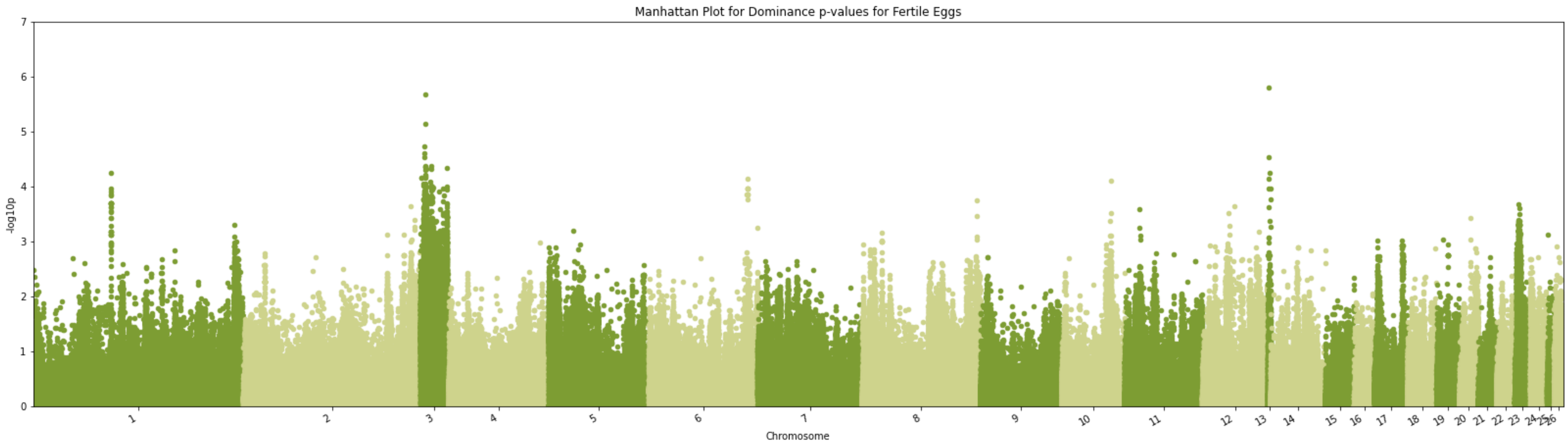


Fig S51. Manhattan plot -log10(p) using the marker p-value from TASSEL GWAS for Fertile Eggs for chromosomes 1 through 26 (Supplemental Table S2 for Chromosome mapping). Dominance model.

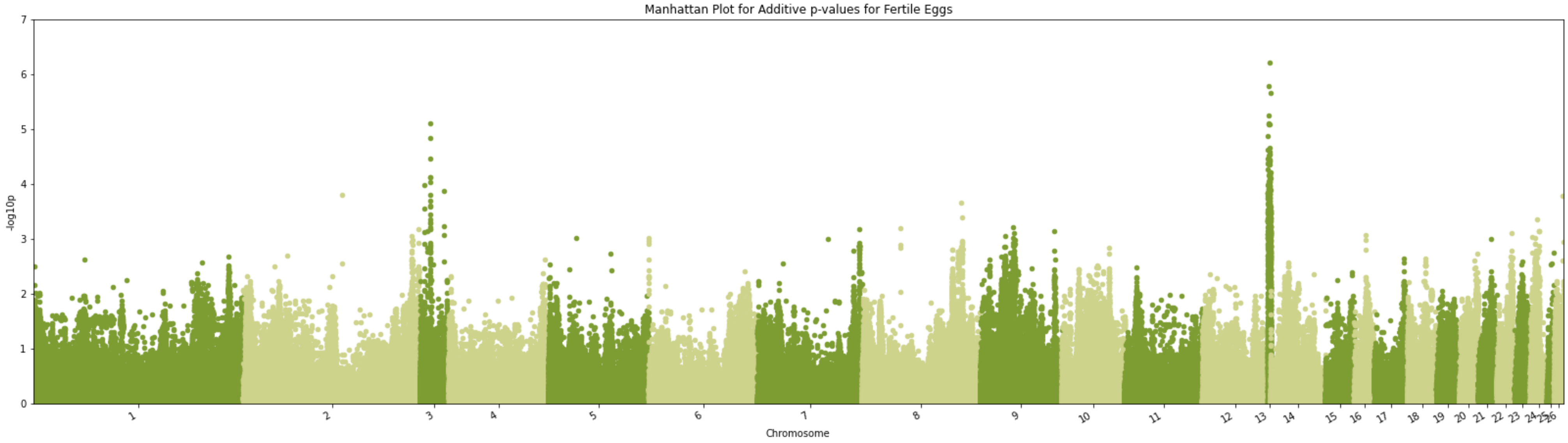


Fig S52. Manhattan plot -log10(p) using the marker p-value from TASSEL GWAS for Fertile Eggs for chromosomes 1 through 26 (Supplemental Table S2 for Chromosome mapping). Additive model.

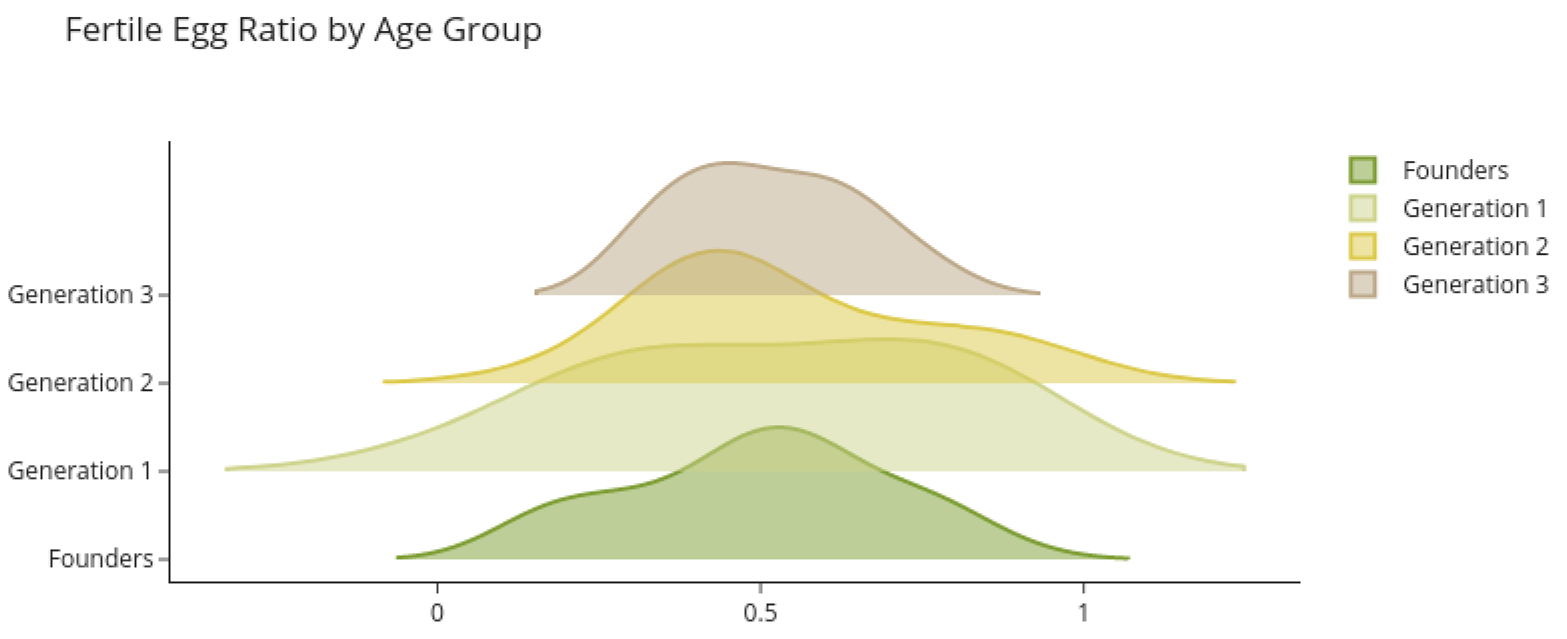


Fig S53. Predicted breeding values for fertile eggs by age group.

#### Embryo Survival

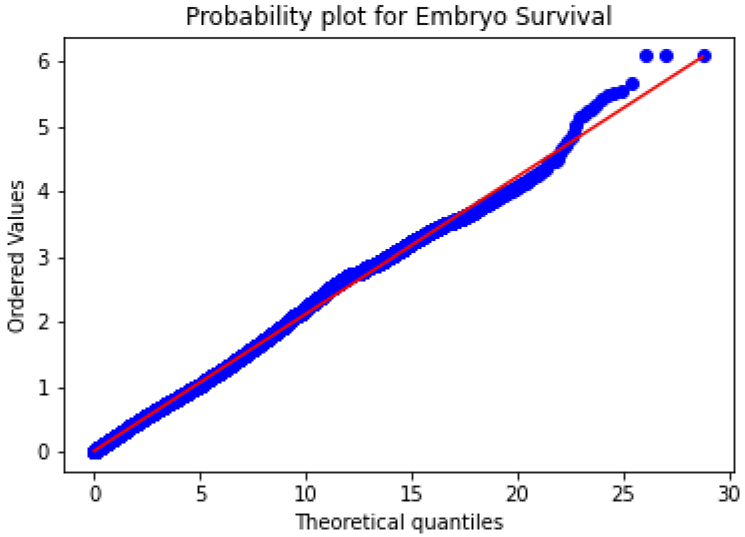


Fig S54. QQ-Plot for Embryo Survival GWAS.

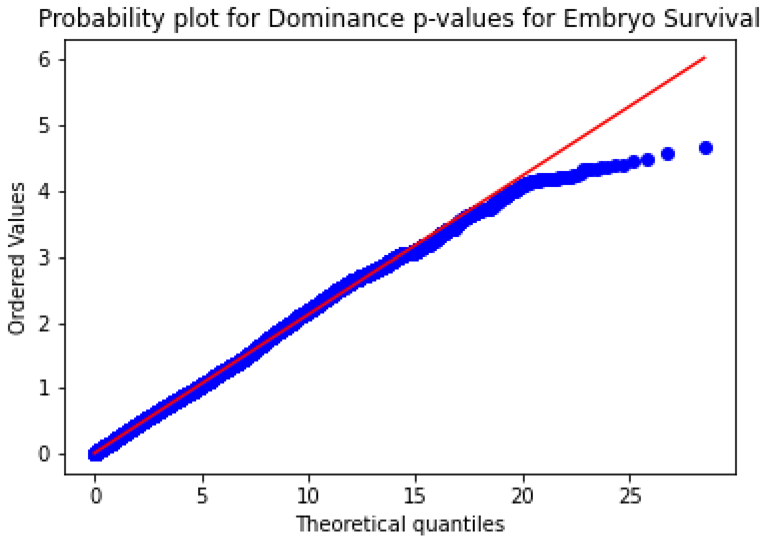


Fig S55. QQ-Plot for Embryo Survival GWAS. Dominance model.

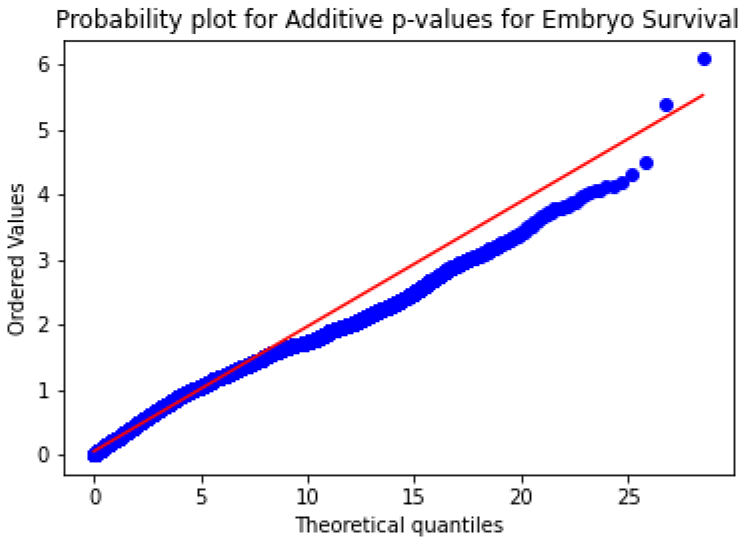


Fig S56. QQ-Plot for Embryo Survival GWAS. Additive model.

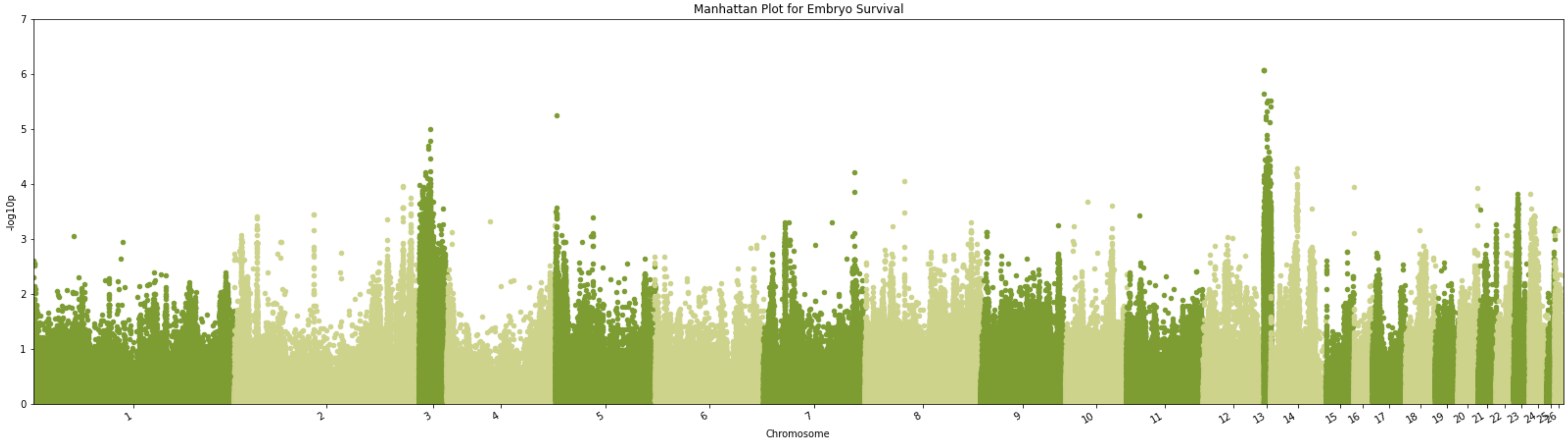


Fig S57. Manhattan plot -log10(p) using the marker p-value from TASSEL GWAS for Embryo Survival for chromosomes 1 through 26 (Supplemental Table S2 for Chromosome mapping).

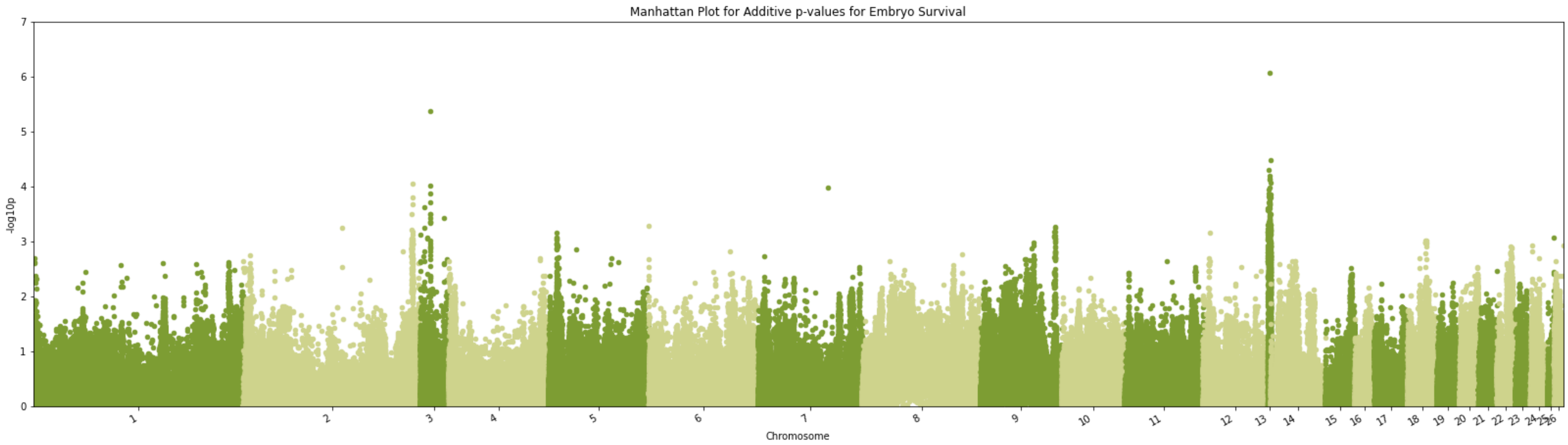


Fig S58. Manhattan plot -log10(p) using the marker p-value from TASSEL GWAS for Embryo Survival for chromosomes 1 through 26 (Supplemental Table S2 for Chromosome mapping). Dominance model.

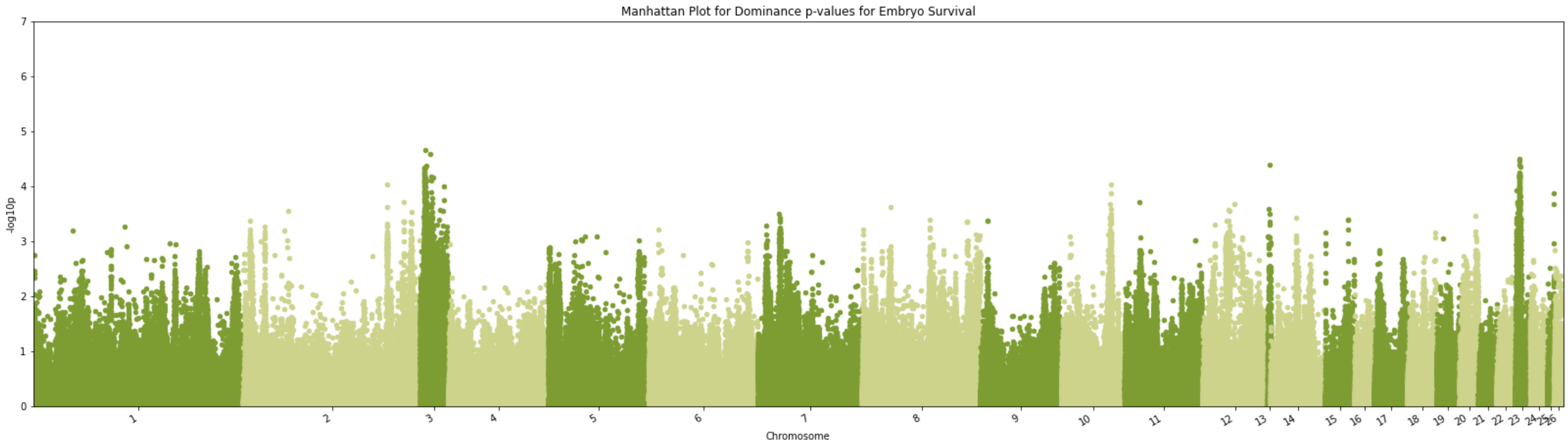


Fig S59. Manhattan plot -log10(p) using the marker p-value from TASSEL GWAS for Embryo Survival for chromosomes 1 through 26 (Supplemental Table S2 for Chromosome mapping). Additive model.

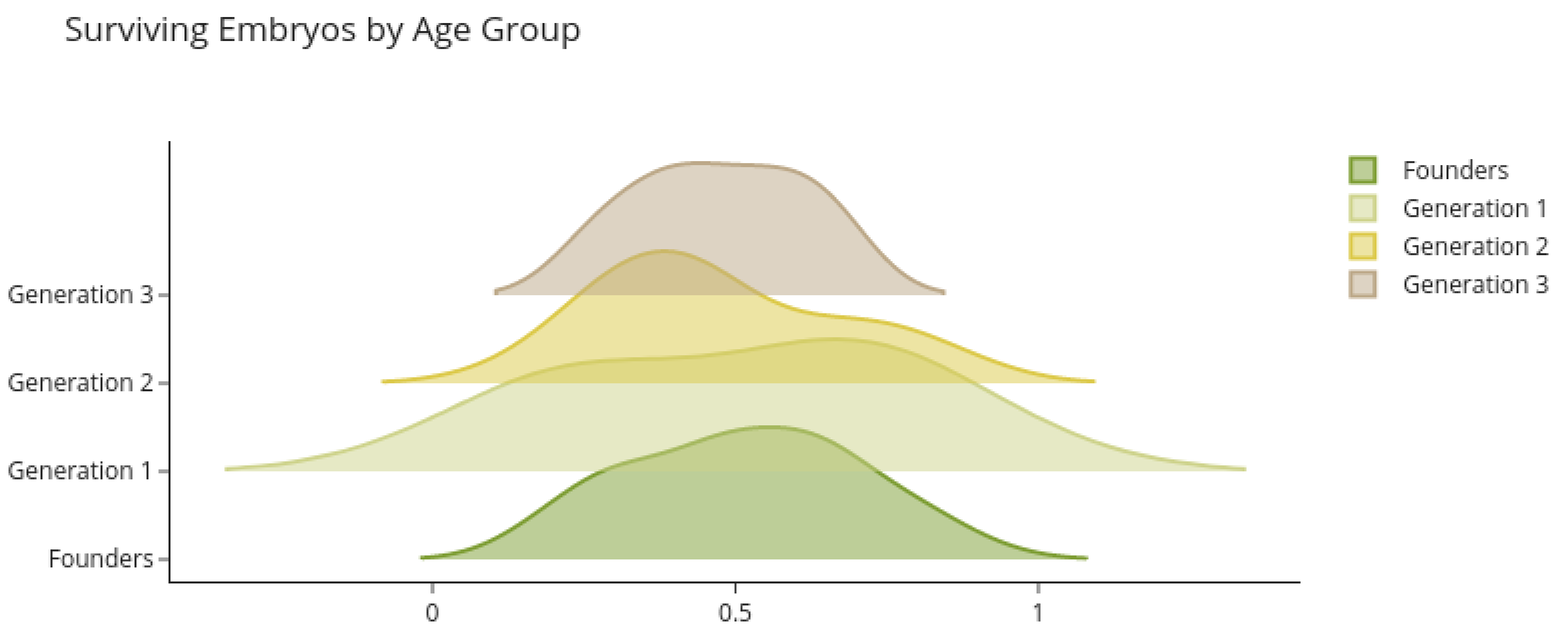


Fig S60. Predicted breeding values for embryo survival by age group.

#### Aspergillosis Susceptibility

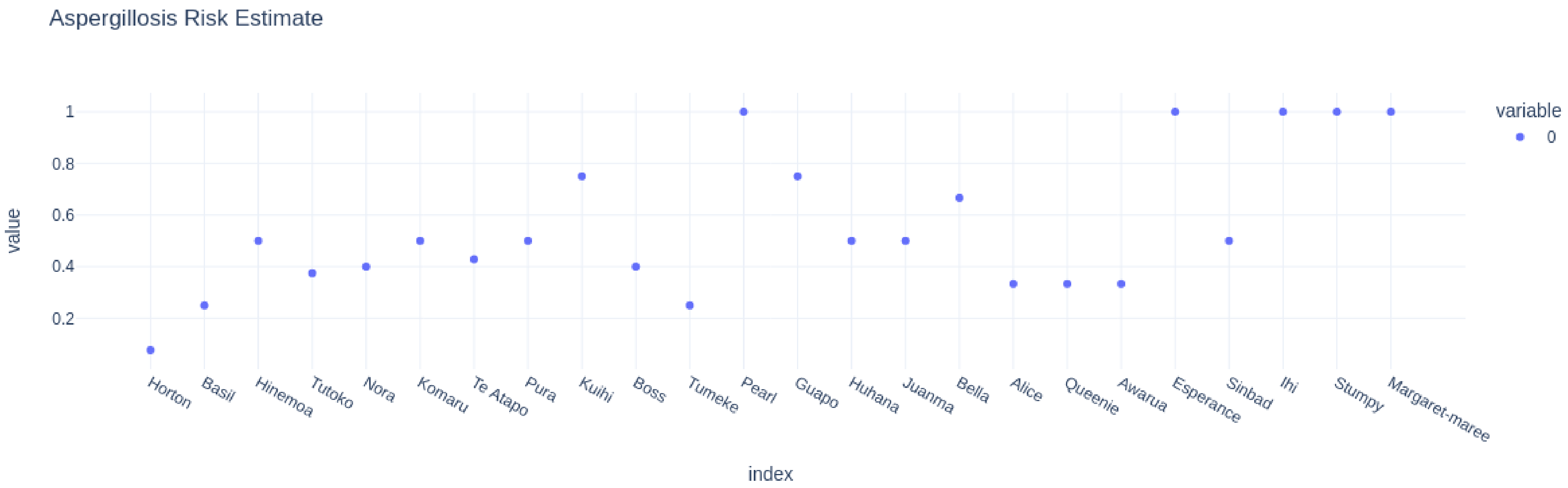


Fig S61. Risk estimate predicted from model.

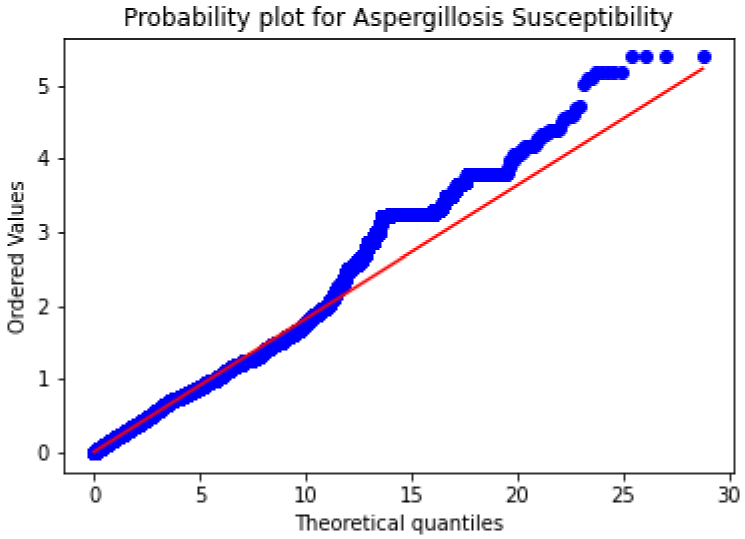


Fig 622. QQ-Plot for Aspergillosis susceptibility.

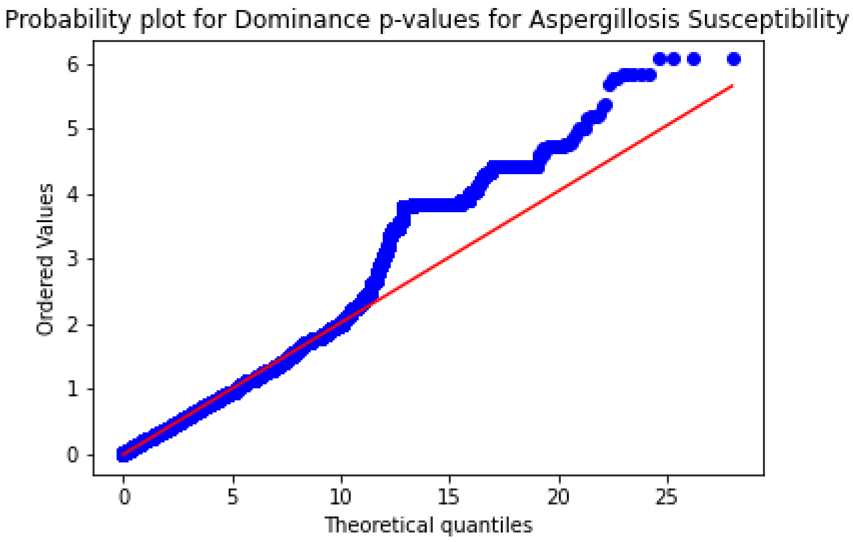


Fig S63. QQ-Plot for Aspergillosis susceptibility. Dominance model.

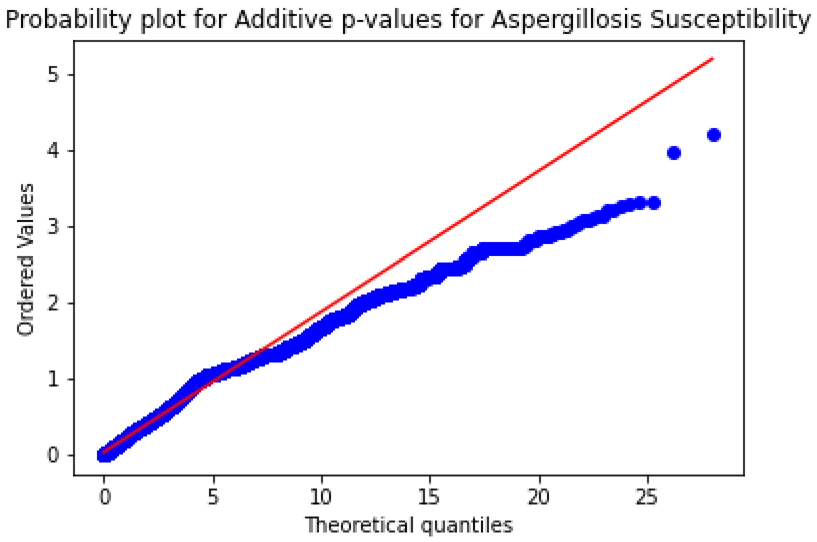


Fig S64. QQ-Plot for Aspergillosis susceptibility. Additive model.

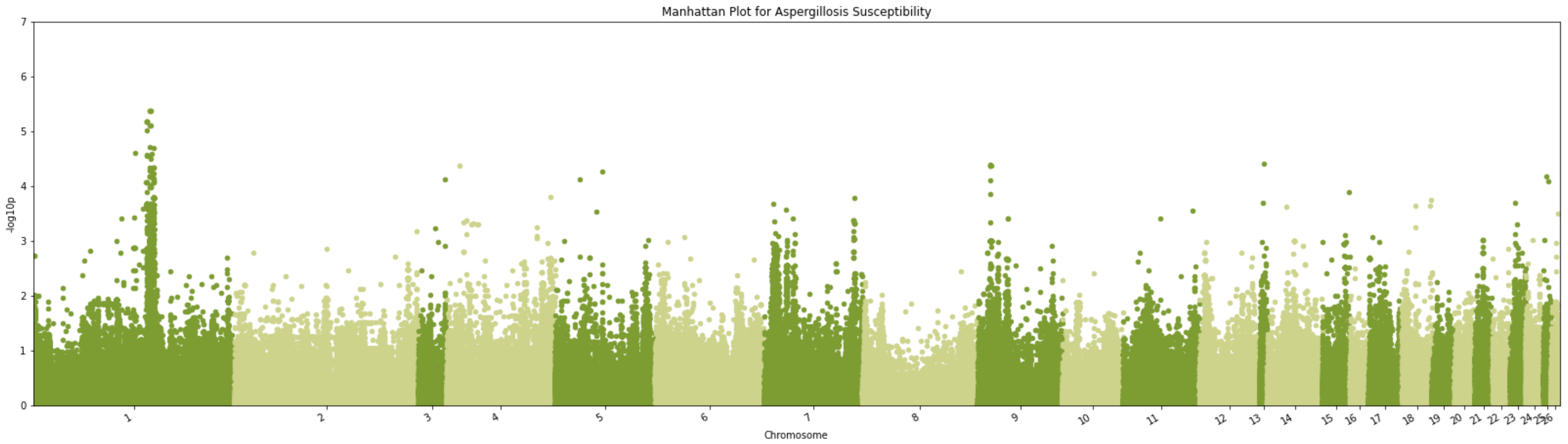


Fig S65. Manhattan plot -log10(p) using the marker p-value from TASSEL GWAS for Aspergillosis Susceptibility for chromosomes 1 through 26 (Supplemental Table S2 for Chromosome mapping).

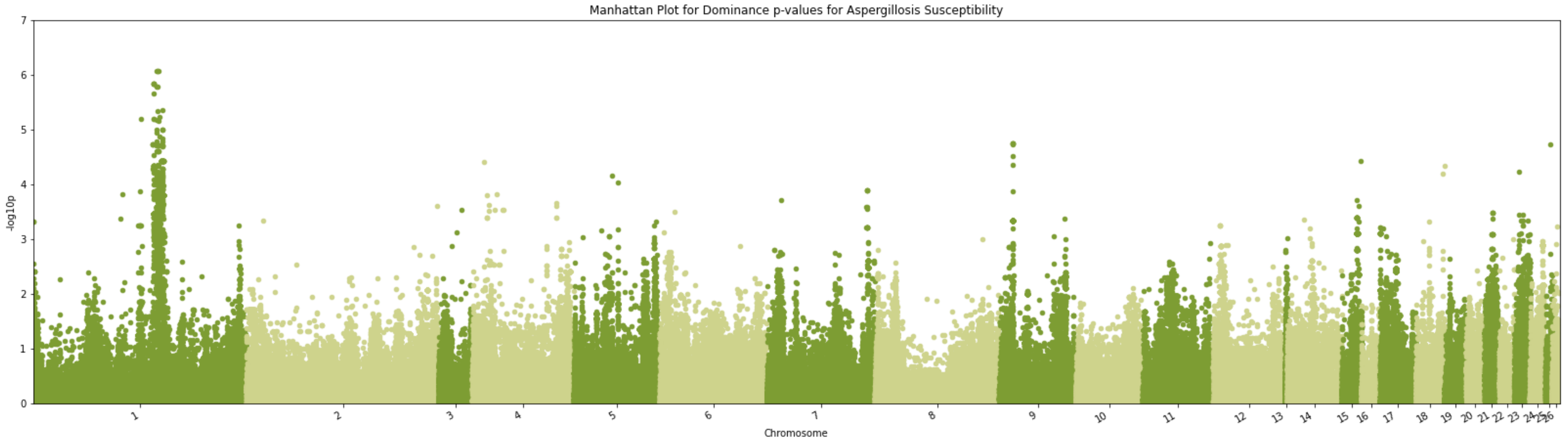


Fig S66. Manhattan plot -log10(p) using the marker p-value from TASSEL GWAS for Aspergillosis Susceptibility for chromosomes 1 through 26 (Supplemental Table S2 for Chromosome mapping). Dominance model.

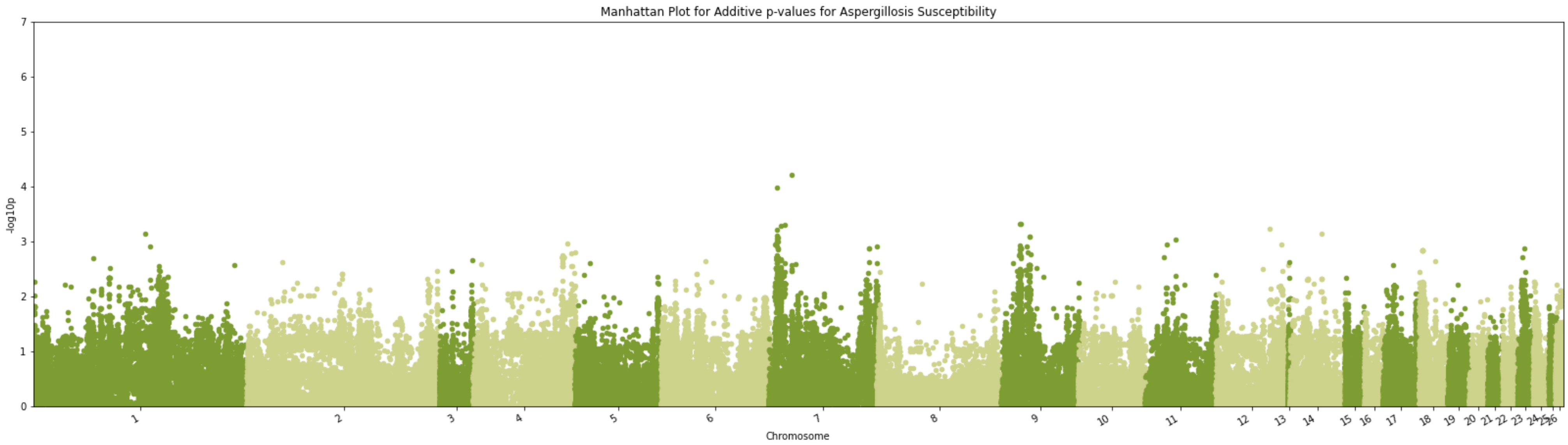


Fig S67. Manhattan plot -log10(p) using the marker p-value from TASSEL GWAS for Aspergillosis Susceptibility for chromosomes 1 through 26 (Supplemental Table S2 for Chromosome mapping). Additive model.

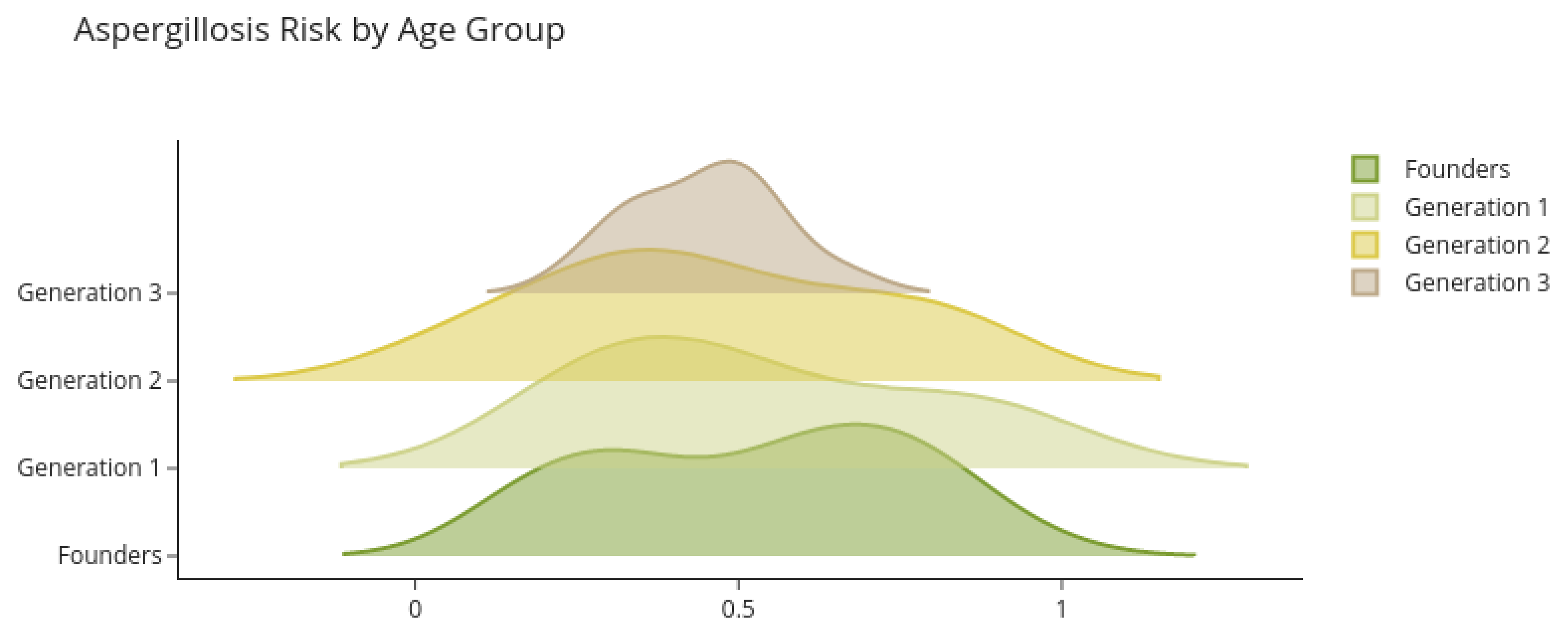


Fig S68. Predicted breeding values for Aspergillosis risk by age group.

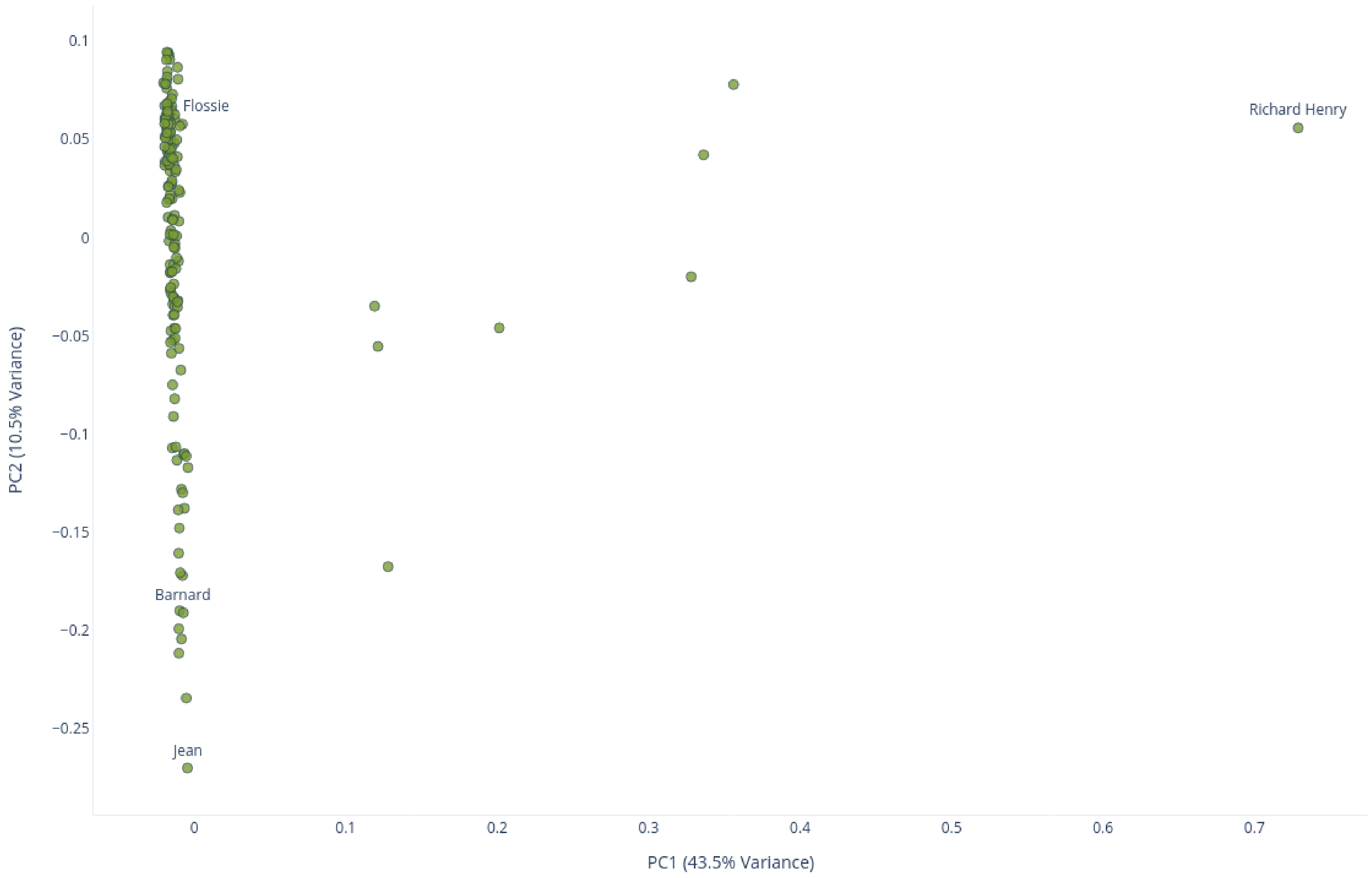


Fig S69. PCA of Our Population. Population PCA determined from biallelic SNPs with fewer than 20% missing allele calls and SNP quality >= 99. Half calls set to missing.

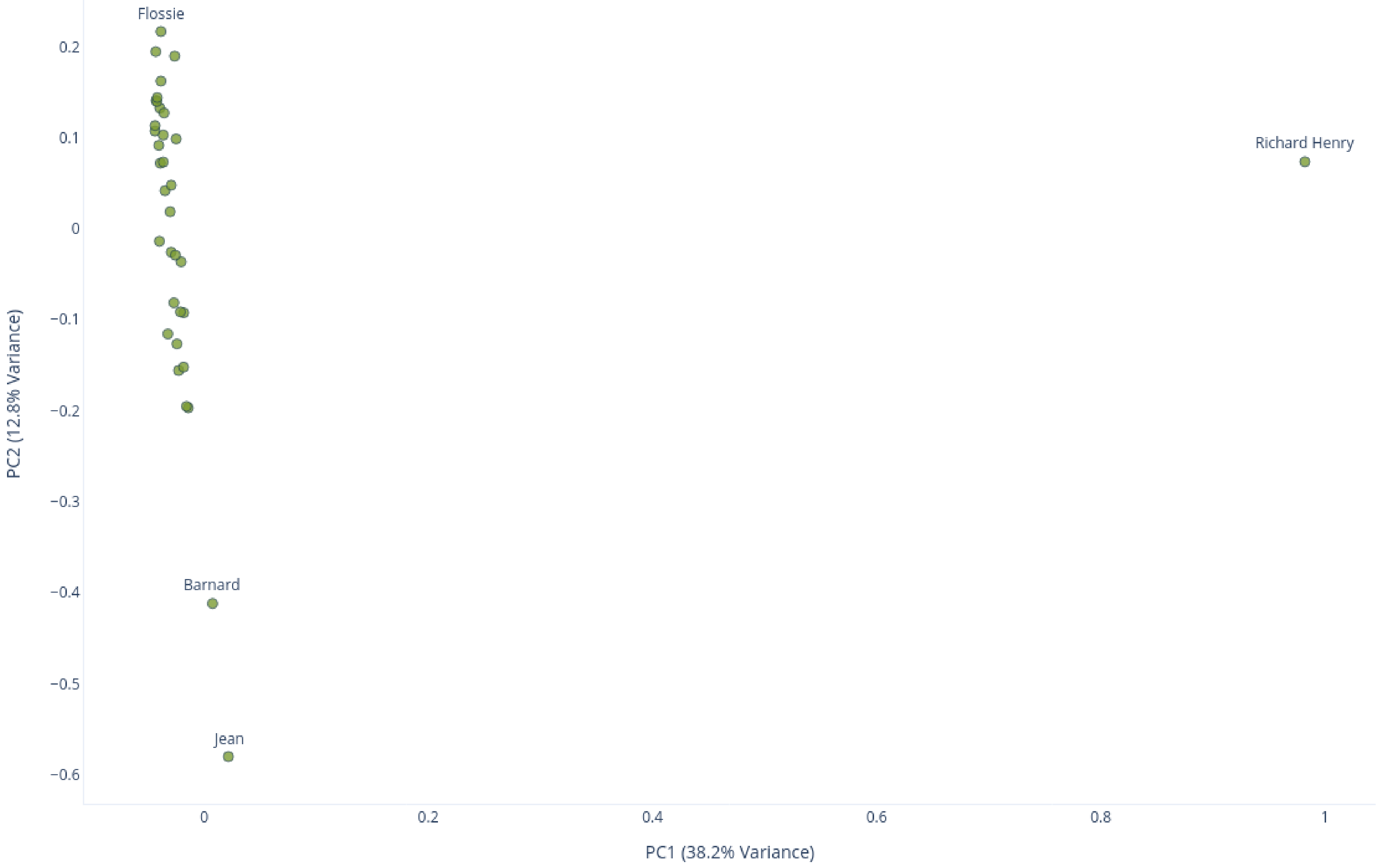


Fig S70. PCA of founders. Founder PCA determined from biallelic SNPs with fewer than 20% missing allele calls and SNP quality >= 99. Half calls set to missing.

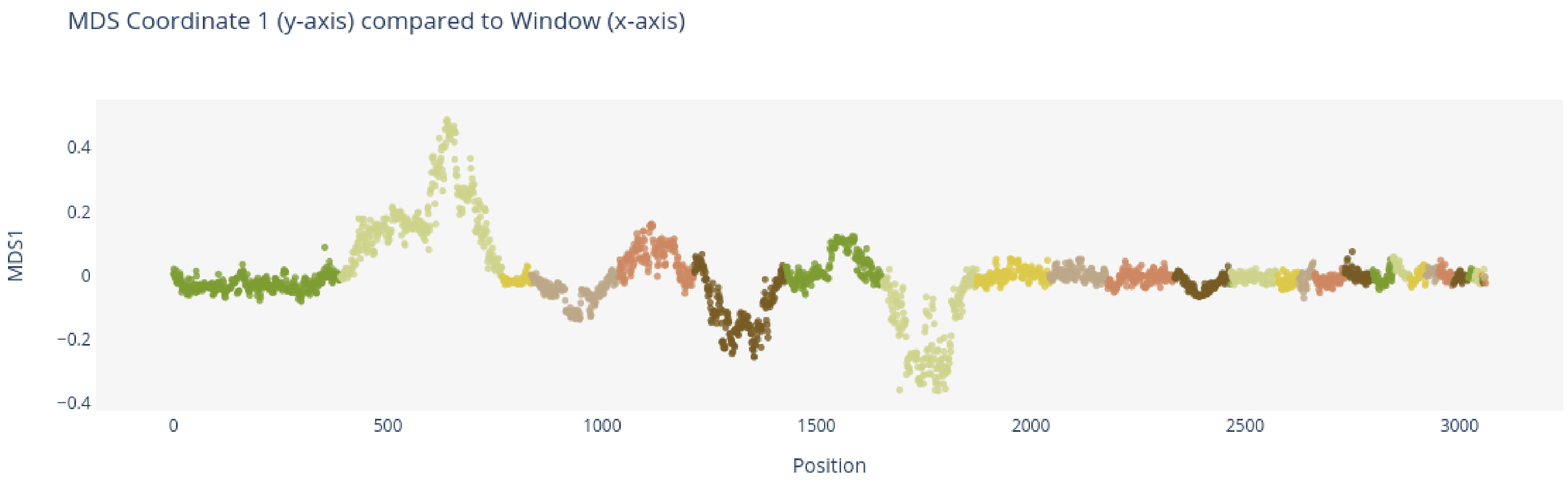


Fig S71. Local PCA MDS Coordinates 1. Each distinct color represents a different chromosome, starting with S1 (Table S2).

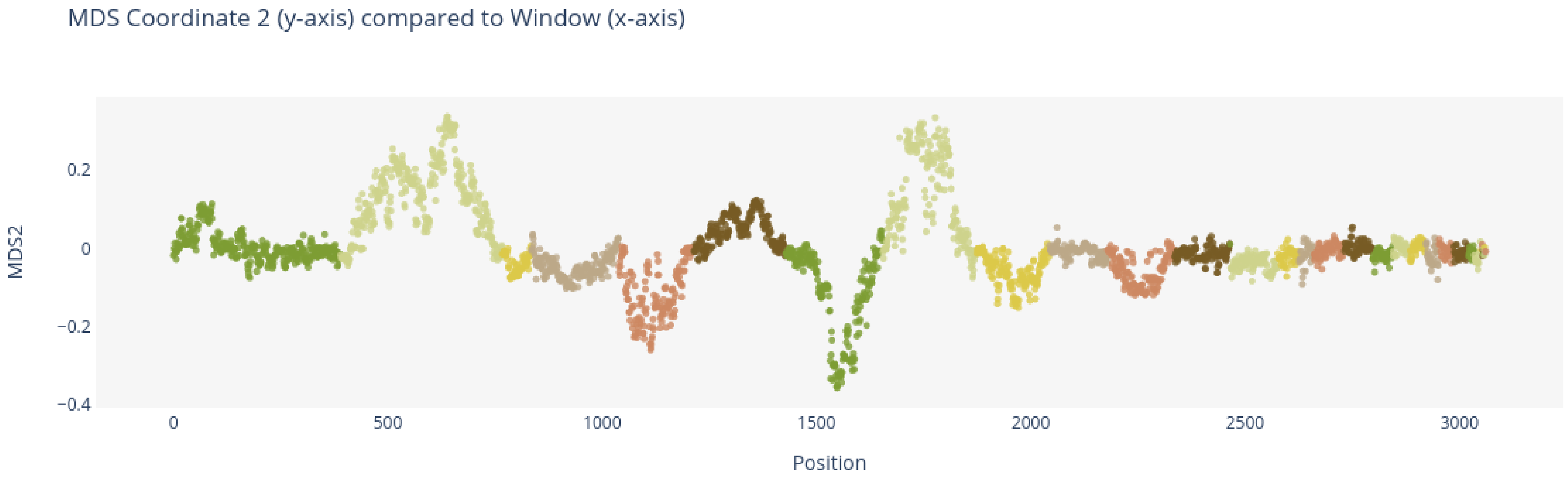


Fig S72. Local PCA MDS Coordinates 2. Each distinct color represents a different chromosome, starting with S1 (Table S2)
